# Nanopore dwell time analysis permits sequencing and conformational assignment of pseudouridine in SARS-CoV-2

**DOI:** 10.1101/2021.05.10.443494

**Authors:** Aaron M. Fleming, Nicole J. Mathewson, Cynthia J. Burrows

## Abstract

Nanopore devices can directly sequence RNA, and the method has the potential to determine locations of epitranscriptomic modifications that have grown in significance because of their roles in cell regulation and stress response. Pseudouridine (Ψ), the most common modification in RNA, was sequenced with a nanopore system using a protein sensor with a helicase brake in synthetic RNAs with 100% modification at 18 known human pseudouridinylation sites. The new signals were compared to native uridine (U) control strands to characterize base calling and associated errors as well as ion current and dwell time changes. The data point to strong sequence context effects in which Ψ can easily be detected in some contexts while in others Ψ yields signals similar to U that would be false negatives in an unknown sample. We identified that the passage of Ψ through the helicase brake slowed the translocation kinetics compared to U and showed a smaller sequence bias that could permit detection of this modification in RNA. The unique signals from Ψ relative to U are proposed to reflect the *syn-anti* conformational flexibility of Ψ not found in U, and the difference in π stacking between these bases. This observation permitted analysis of SARS-CoV-2 nanopore sequencing data to identify five conserved Ψ sites on the 3’ end of the viral sub-genomic RNAs, and other less conserved Ψ sites. Using the helicase as a sensor protein in nanopore sequencing experiments enables detection of this modification in a greater number of relevant sequence contexts. The data are discussed concerning their analytical and biological significance.

## Introduction

The epitranscriptome represents the collection of chemical modifications on mRNA strands that are essential for their biological functions in cells. Complete nuclease digestion and mass spectrometric analysis of cellular RNAs from many organism types have identified >170 different modifications, a large subset of which have been found in humans.^1–4^ One drawback to this analytical approach is the fact that sequence information regarding the sites modified, the specific RNA, and the extent of modification at a given site is lost. This information is essential to address how the epitranscriptome impacts RNA physiology. Methods developed for sequencing target modifications include high accuracy but low-throughput approaches such as SCARLET.^5^ At present high-throughput methods relying on modification enrichment and next-generation sequencing (NGS), best illustrated by the many *N^6^*-methyladenosine (m^6^A) sequencing approaches reported, have enabled many discoveries of the m^6^A epitranscriptome.^3,6^ Development of techniques to sequence other RNA modifications with high accuracy and in a quantitative fashion are desperately needed to continue our exploration of the epitranscriptome.

Pseudouridine (Ψ), the most abundant global RNA modification, is an isomerization product of uridine (U) found at ~7% of all Us in all RNAs except mRNA where Ψ is found at ~0.1% occupancy (Figure 1A).^7^ This modification is tied to critical RNA functions in cells such as reinforcing RNA secondary structure and regulation of translation, and Ψ levels change in response to oxidative, micronutrient, or heat-shock stress.^7–10^ Initially Ψ was located in RNA via digestion and chromatographic methods followed by mass spectrometric quantification approaches,^11–13^ and recently, high-throughput NGS using Ψ-specific chemistry has enabled sequencing the mammalian transcriptome for Ψ. Pseudouridine is specifically alkylated by the carbodiimide CMC to yield a stable and bulky adduct to stall reverse transcription, which is found by distinct sequencing read stops in comparison to a non-alkylated matched control.^10,14–16^ Alternatively, Ψ can be sequenced by the absence of reactivity with hydrazine while the parent U readily reacts to yield strand breaks detected during NGS.^17^ As a result of analyzing sequencing stops, these approaches cannot achieve read-through in order to sequence multiple sites in single strands; additionally, RNA structure-induced stops can yield false positives, and quantification of the modification is challenging to conduct accurately using these approaches.^18,19^ One chemical solution is to treat the suspect RNA with bisulfite to yield a stable sugar adduct on the Ψ nucleotide,^20,21^ which induces a deletion signature during cDNA synthesis, allowing more than one modification per strand to be sequenced. However, this approach does not yield quantitative deletions, and therefore, the extent of modification at suspected sites is challenging to obtain.

**Figure 1.**
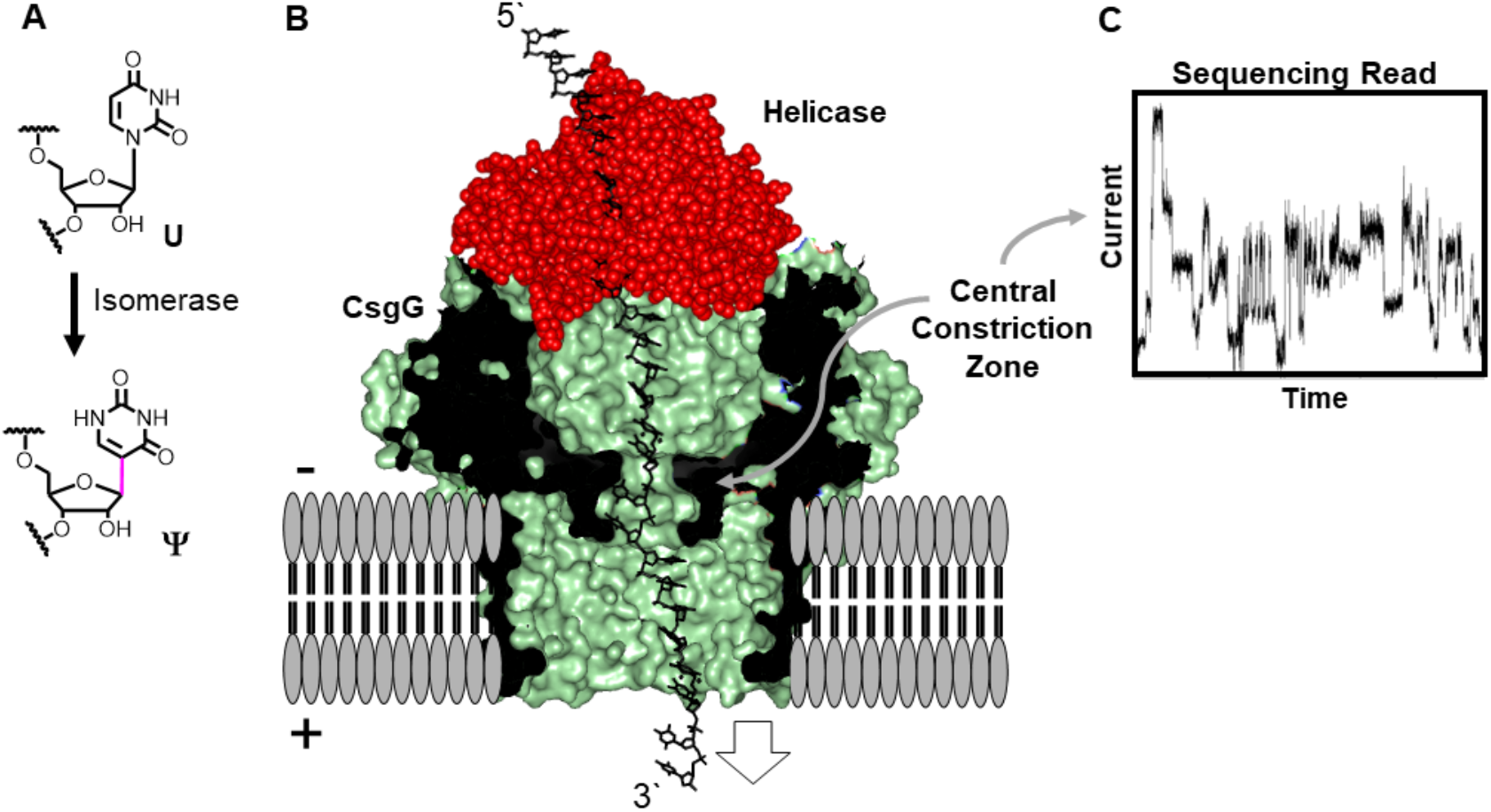
Direct RNA sequencing for Ψ by monitoring current vs. time traces in a protein nanopore-helicase platform. (A) Isomerization of U yields Ψ. (B) Structural depiction of the CsgG-helicase nanopore setup used in the R9.1.4 MinION/Flongle flowcells manufactured by ONT. (C) Example ion current vs. time trace. This figure was made using the PDB files 4UV3^22^ and 3UPU^23^ that were selected based on the description of this system in the literature.^24^

Third-generation sequencing (TGS) technologies utilizing the Oxford Nanopore Technologies (ONT) or the Pacific BioSciences waveguide platforms sequence single molecules of DNA or RNA that allow modifications to be directly observed.^25^ By directly sequencing DNA or RNA, steps that introduce biases in the workflow can be omitted such as the need for high-yielding and selective chemistry, reverse transcription, and/or PCR. The ONT or MinION™ device sequences RNA by ratcheting the strand 3’ to 5’ via a motor protein (e.g., helicase) through a protein nanopore sensor under an electrophoretic force (Figure 1B). The current modulates as the strand passes through the nanopore sensor in a characteristic way to call the individual nucleotides using a trained neural network algorithm (Figure 1C). Because the signal from the sensor results from molecular interactions between the RNA and the protein nanopore it is anticipated that epitranscriptomic modifications will interact differently to generate unique signatures for their identification. Direct nanopore sequencing applied to synthetic RNAs with 100% modification present (e.g., m^6^A, Ψ, N7-methylguanosine (m^7^G), 5-hydroxymethylcytidine, etc.) have demonstrated the feasibility of this approach.^26,27^ Sequencing of the 16s rRNA from *E. coli* with the MinION™ successfully called m^7^G and Ψ sites,^28^ and many examples of calling m^6^A in cellular RNAs have been reported demonstrating that direct sequencing for modifications can work.^27,29,30^ Lastly, long-range interactions observed in the nanopore sensor when the modification passes through the motor protein have been noted.^31,32^ Nanopore sequencing has great potential to enable complete and quantitative sequencing of the epitranscriptome.

Herein, synthetic RNAs were made with Ψ in 18 known human sequence contexts found in rRNA, tRNA, or mRNA. Sequencing of the contexts found Ψ can be called predominantly as a U or C, while the U:C base call ratio and associated errors are highly sequence-context dependent resulting in an incoherent Ψ signature. The raw ionic current and dwell time data from the sequencer were then analyzed to identify significant differences between populations of RNAs with either the parent U or Ψ at the target sites; however, the sequence context impacted the signal differences that again would bias data during sequencing of unknown samples. Inspection of the dwell time differences between the RNAs with U or Ψ found a long-range signature resulting when Ψ translocated through the active site of the helicase motor to yield a more sequence agnostic signature. We used the helicase dwell differences in conjunction with the other data analysis approaches to inspect publicly available nanopore sequencing data for the SARS-CoV-2 RNA sub-genomes for Ψ. The data analysis converged on five conserved sites with high confidence at which Ψ resides in the viral RNA sub-genomes near the 3’ end. The results are discussed concerning their analytical and biological impact.

## Methods

### RNA synthesis

The RNA transcripts were synthesized by two different approaches. The first used short RNA strands (~60 mers) that were synthesized by solid-phase synthesis to have the modifications studied in sequence contexts with U adjacent to Ψ. Second, in vitro transcription was used to synthesize longer RNAs utilizing duplex DNA templates containing the T7 promoter and a poly-A tail necessary for library preparation using the direct RNA sequencing kit from ONT (see Figure S1 for complete RNA sequences synthesized). In vitro transcription (IVT) was performed using the MEGAscript T7 transcription kit according to the manufacturer’s instructions. The IVT reactions were incubated overnight at 37 °C. Then, DNase I treatment was performed on all samples, followed by purification using Quick Spin Columns for radiolabeled RNA purification. To install Ψ, IVT was conducted in the presence of pseudouridine-5’-triphosphate (ΨTP) instead of uridine-5’-triphosphate (UTP). Success in the synthesis of the RNA transcripts was verified by agarose gel electrophoresis by comparison to a ladder of known lengths.

### Library preparation and nanopore sequencing

The poly-A tail containing RNAs generated by IVT were the input strands in the direct RNA sequencing kit (SQK-RNA002) from ONT. Before library preparation of the rRNA, the sample was treated with a 3’-poly-A tailing kit to generate strands that can be used in the direct RNA sequencing kit. The protocols were followed without changes and the library prepared samples were directly used for sequencing. The samples were applied to either the ONT MinION™ or Flongle™ flowcells using the R9.4.1 chemistry following the manufacturer’s protocol with an applied voltage of 180 mV. All sequencing experiments were conducted for 12 h, and as expected, the MinION™ produced >10x more reads (~300,000) per experiment than the Flongle™.

### Data analysis

The fast5 files were base called using Guppy v.4.0 to obtain the fastq sequencing read files used in the subsequent data analyses. The data were first analyzed by MultiQC implemented in the Master of Pores tool,^33^ to yield read statistics similar to those described in the literature (Figures S2).^34^ The fastq files were aligned to reference sequences using minimap2 to generate bam files.^35^ The bam files were indexed with Samtools and then visualized in IGV to obtain the base call frequency at the modification sites.^36,37^ The Eligos2 tool was used as described in the online manual on GitLab,^27^ and the Nanocompore and Tombo tools were used as described on their Github sites.^29,38^ The current and dwell time data were extracted, indexed to the reference, and analyzed using Nanopolish as described in the user manual.^39^ The data were plotted and analyzed in either python, Origin, or Excel for visualization.

### Publicly available data

The nanopore sequencing reads^40^ and Nanocompore analysis^41^ for the SARS-CoV-2 RNA genome and synthetic RNA void of chemical modifications have been reported and deposited in publicly available databases for analysis. The data were reanalyzed by the methods outlined in the present manuscript using the Wuhan-Hu-1 reference genome NC_045512.2 that was corrected to match the South Korea variant with four single nucleotide substitutions (T4402C, G5062T, C8782T, and T28143C) as described in the initial report of these data.^40^

## Results

The CsgG protein nanopore in the R9.4.1 sequencing flowcells has a 5-nt sensing zone (i.e., k-mer) for RNA;^42^ thus, the duplex DNA templates utilized for RNA synthesis by IVT were designed to install UTP or ΨTP centrally within a 5-nt window comprised of known pseudouridinylation sites in human rRNA, tRNA, and mRNA that only had one U nucleotide in the window (Figure S1).^43,44^ To study Ψ contexts that include a U, shorter RNAs were synthesized by solid-phase synthesis and then sequenced. Unique to the present studies is that the Ψ sites were separated by >25 nts to allow their individual study as they moved through the sequencer top to bottom (helicase to nanopore); this would be the most common scenario in which Ψ would be sequenced from biological samples except where they are clustered as described below. This approach contrasts with those that synthesize model mRNAs with all U sites converted to Ψ.^27^ In the sequence contexts bearing doubly or triply modified sites, 2 nt of the biologically adjacent sequence were maintained on either side of the modified region (Figure S1). Different RNA strands were sequenced with some redundancy in the contexts to test the reproducibility of the sequencer from one strand to the next as well as at different positions relative to the ends of the RNA. After library preparation using the direct RNA sequence kit from ONT, the samples were sequenced on either a MinION™ or Flongle™ flowcell using CsgG nanopores (R.9.1.4) to achieve ~300,000 or ~30,000 reads, respectively, and the fast5 reads were base called using the Guppy tool to yield fastq files for further analysis.

The sequencing reads were aligned to the reference using minimap2 to identify the base call identities for the 18 different sequence contexts studied. For the RNAs containing U, the called nucleotide was >90% U with the remainder called as C or A with sequence context dependency (Figure 2A). When Ψ was present, the nucleotide called was predominantly a mixture of C and U and a low amount of A or G, consistent with prior results.^26–28^ The new finding herein is the distribution of C and U called at the 13 singly-modified Ψ sites ranged from 10% to 97% C with the remainder predominantly called as U (Figure 2A). Inspection of the sequence-dependent base calling results for Ψ did not lead to an obvious sequence context trend. Demonstration of reproducibility in the base calls was achieved by sequence redundancy in the strand design to identify similar base calls for Ψ (Figure S3). The doubly and triply modified sites produced similar U vs. C base call signatures as the singly modified sites, as well as impacting base calling on the adjacent canonical nucleotides (Figures 2B and 2C). These data indicate that the Guppy algorithm, which was trained on canonical RNA nucleotides, when confronted with Ψ called the site as a C or U with dependency on the sequence context.

**Figure 2.**
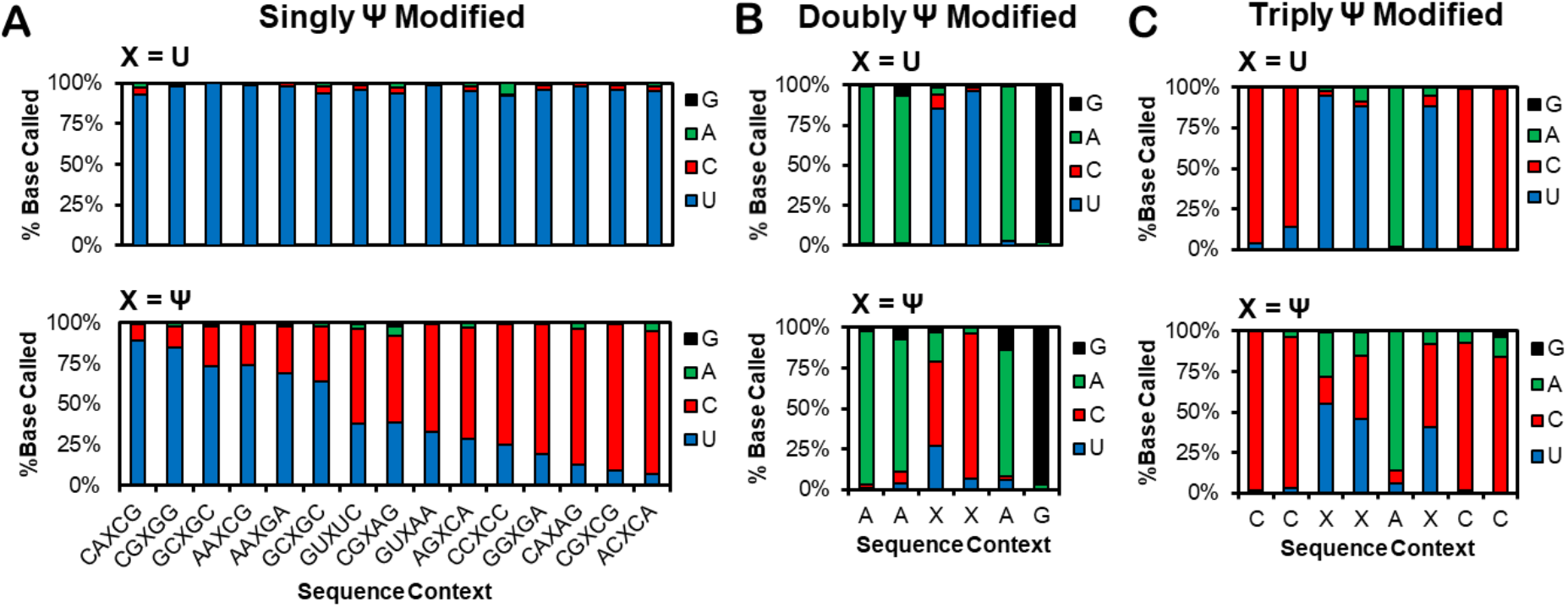
Base-calling frequencies from direct RNA nanopore data when U or Ψ are present in biologically relevant sequence contexts. The bases are called for U or Ψ in (A) singly, (B) doubly, or (C) triply modified contexts. The raw nanopore reads were base-called with Guppy v.4.0 followed by minimap2 alignment, IGV visualization and analysis of 30,000 reads.

The Eligos2 tool calls chemically modified sites in RNA from nanopore data by sample comparison of a modified strand with a matching strand depleted or void in the modification to inspect for differences in the error of specific bases (ESB) (Figure 3A and S4).^27^ Radar plots illustrate that the ESB values increase at Ψ sites and are impacted by adjacent nucleotides (Figure 3A). The ESB values allow calculation of an odds ratio (oddR) or *P*-value of statistical significance for the presence of modifications (Figures 3B and S4). Application of Eligos2 to locate Ψ when compared to a U-containing RNA of the same sequence identified observable oddR values at the modified sites (Figure 3B), consistent with the literature.^38^ The oddR values ranged from 4 to 240 for the singly Ψ-modified sites with dependency on the sequence context (Figure 3B). A trend was observed in which those Ψ sites called to a greater extent as C gave the higher oddR values (Figures 2A and 3B). The doubly and triply modified sites were also detectable by the Eligos2 tool (Figure S4). Comparison of the redundant Ψ-modified contexts in the RNAs studied found Eligos2 can yield similar statistical values for some of the modified sites selected for study and others were found vary considerably between the test cases (Figure S5). Using base-calling errors provides a means to locate Ψ in a sequence, but the corresponding signals are sequence context dependent that favorably bias detection of Ψ to sequence contexts that yield greater C base calls and base-calling error.

**Figure 3.**
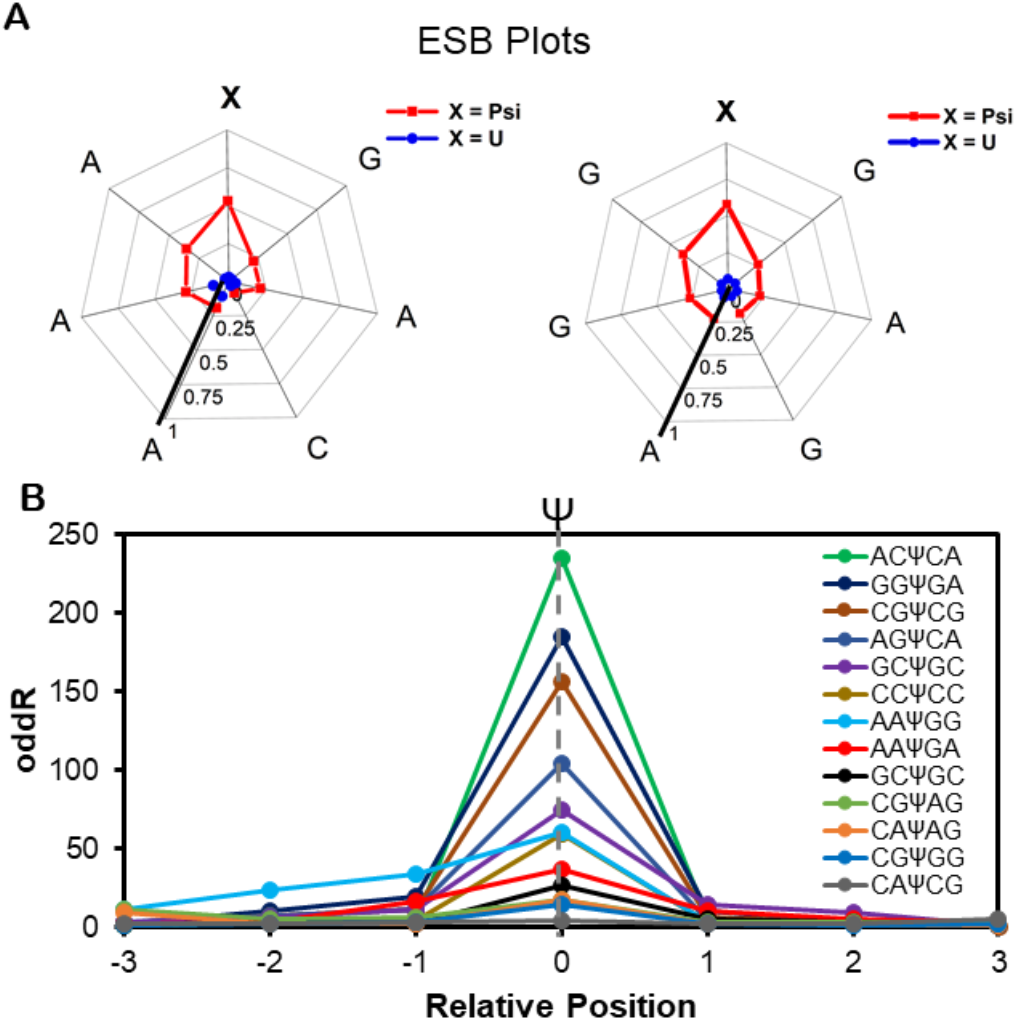
Eligos2 analysis to locate Ψ in direct RNA sequencing data shows sequence context-dependency in the magnitude of the oddR values. (A) Two example radar plots of ESB values for singly modified Ψ versus U sites in RNA. (B) A plot of oddR values that were found for the Ψ modified sites in the RNAs studied. More example ESB plots and the reproducibility plots for the oddR values found for Ψ are provided in Figures S4 and S5.

The ONT single-molecule sequencing platform reads the nucleotide sequence by using an electrophoretic force and a helicase brake to slowly move the strand through a small aperture protein pore.^24^ As the nucleotides pass through the constriction of the nanopore that is ~5 nt long (i.e., k-mer) for RNA,^42^ the ionic current is deflected with dependency on the sequence identity inside the aperture. Modified nucleotides have different sizes, shapes, and/or hydrodynamic properties that permit changes in the current and/or dwell times to be detected compared to the canonical forms, resulting in their identification. Thus, the raw nanopore data can be analyzed for the presence of RNA modifications.

Interrogation of the current intensity and dwell times requires aligning the base-called data to the events using either Tombo or Nanopolish to inspect the population of single-molecule reads at each point (Figures S6 and S7). The Nanocompore tool takes in the Nanopolish event alignments and provides the ability to compare the populations on modified RNAs against a matched population void in the modification. The tool compares the two samples in a pairwise fashion using the Kolmogorov-Smirnov test to report *P*-values of statistical differences between the RNA populations. The *P*-values are -log transformed to visualize sites with a significant difference by an increased value. Thirteen of the biologically relevant Ψ-containing sequence contexts were analyzed for differences in signal when the modification passes through the protein nanopore sensor (Figure 4A). The single-molecule events were compiled to make histograms of the current intensities or dwell times for the U or Ψ samples at each sequence read frame (Figures 4B and 4C; top or blue = U, bottom or red = Ψ). Inspection of the current histograms for U vs. Ψ as the suspect site moves through the 5-nt window of the sensor for each sequence context identified significant sites based on the Nanocompore analysis (Figures 4D and S7). The key observation regarding the current-level differences is that Ψ impacts the signal but the position at which the impact is most significant within the known 5-nt window is dependent on the sequence context.

**Figure 4.**
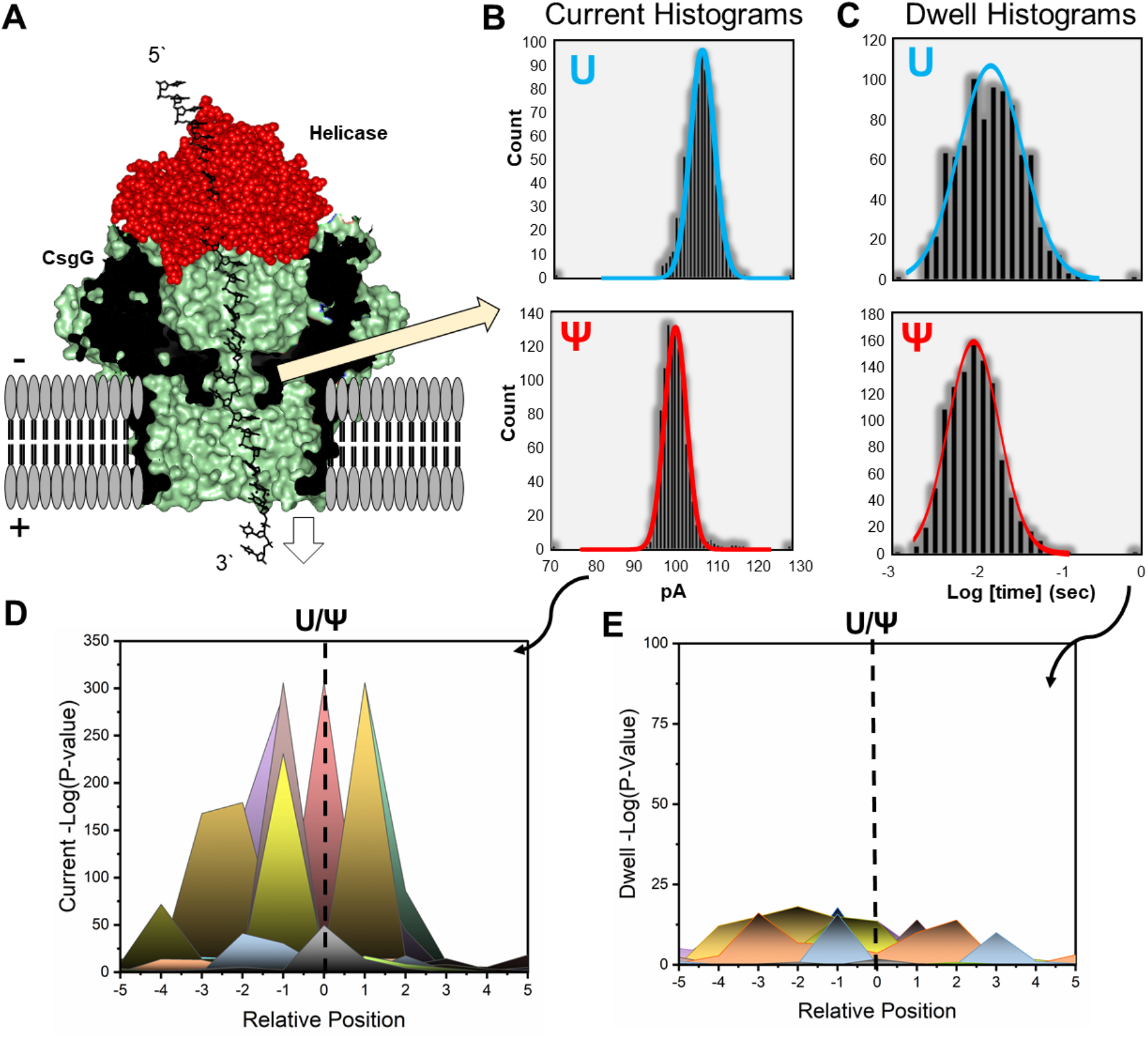
The current intensity and dwell time analysis of RNA with U or Ψ passing through the protein nanopore sensor. (A) A representation of the helicase-nanopore sequencing set up for illustrating where the data are analyzed. (B) Example current histograms and (C) dwell time histograms for U or Ψ in the nanopore sensor, in which the distributions were analyzed with Nanocompore to identify sites of statistical differences between the populations by pairwise analysis using the Kolmogorov-Smirnov test. The *P*-values from the statistical test were -log transformed to visualize the results of the test by increased signal at those most different at each site based on (D) current and (E) dwell time. The analysis was conducted across 10 nt in which the modification could span the 5-nt window of the protein nanopore sensor region. The plots were constructed from >800 data points obtained from Nanopolish extraction of the currents and dwell times from the raw fast5 data files. More example histograms can be found in Figure S7.

Analysis of the dwell time differences between U vs. Ψ in the contexts studied found a similar observation; Ψ can change the dwell time compared to U in the nanopore sensor but the position in the known 5-nt window at which the difference is maximal is dependent on the context. Because of this variability in the maximal difference in signal, the resolution to call the modified site in an unknown sequence is 9 nt (i.e., k-mer = 5 nt window flanking both sides of a centrally located modification). The redundant sequence contexts sequenced and analyzed were compared to evaluate the reproducibility of the -log transformed *P*-values to find they are poorly reproducible (Figure S8). This is not surprising because of sampling errors leading to different levels of statistical significance.

During the inspection of the differences in current levels and dwell times between the RNAs with U or Ψ, a long-range change to the dwell time population analysis was observed 10-11 nts 3’ to the suspect sites. Sequencing RNA on the ONT platform occurs in the 3’ to 5’ direction (Figure 5A),^24^ and therefore, we propose the long-range difference found only in the dwell time analysis is a result of the modification impacting the helicase activity (Figures 5B-5C). The 10-11 nt registry difference between the helicase active site and the nanopore sensor is supported by a similar observation previously reported between 10-12 nt.^31^ More specifically the current analysis at positions 10-11 3’ to the modification site when it is in the helicase did not significantly change based on Nanocompore analysis (Figure 5D); however, the dwell times produced a long-range signature for sequencing Ψ observed for each sequence context (Figure 5E).

**Figure 5.**
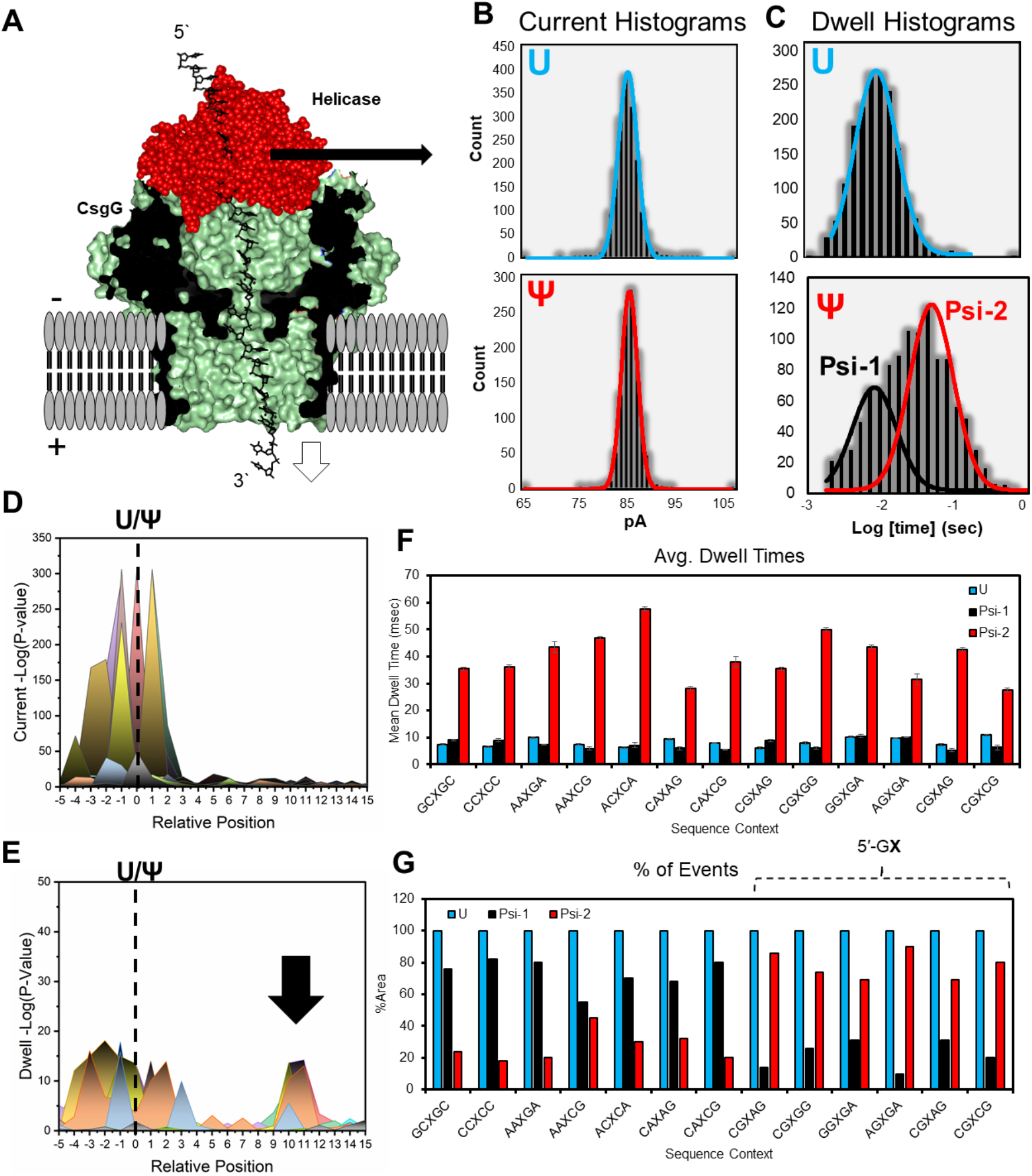
Passage of Ψ through the helicase active site impacts the read dwell time compared to U that permits detection of the epitranscriptomic modification. (A) Schematic of the helicase-protein nanopore setup. Example (B) current and (C) dwell time histograms for a U or Ψ in the active site of the helicase. Stacked plots of -log(*P*-Values) from the Nanocompore analysis for the complete passage of the suspect sites through the nanopore setup looking at statistical differences in the (D) current and (E) dwell times. (F) The median dwell times for the site most statistically significant based on Nanocompore analysis when it resides in the helicase. (G) Plot of the areas for the median dwell time distributions. The analyses were conducted on >1000 single-molecule measurements. Additional data are provided in Figure S9.

The dwell time distributions for helicase stalling at the maximally different sites when plotted on a log axis were found to be Gaussian distributed for the U populations and bimodal Gaussian distributed for the Ψ populations (psi-1 and psi-2; Figures 5B-5E). The distributions were fit to Gaussian equations (r^2^ > 0.95) allowing determination of the average event time for the populations (Figures 5B and 5C). First, the U sites were processed by the helicase with average event times in the range of 6-13 msec (Figure 5F blue bars). The event time sub-population labeled psi-1 gave an average time similar to the U population (Figure 5F black bars or psi-1). In contrast, the second sub-population for Ψ labeled psi-2 produced a longer average time of 10-100 msec (Figure 5F red bars or psi-2). The psi-2 sub-populations were >3-fold longer in average time than the psi-1 sub-populations for the same events. This observation found the helicase activity on Ψ as a substrate is split into two populations that results from the modified nucleotide interacting with the active site of the helicase in two different conformations.

The two Ψ helicase activity sub-populations were then integrated to determine whether the populations change as a function of sequence context (Figure 5G). The analysis found 5’-GΨ sequence contexts gave >70% of the total area as the psi-2 sub-population with a larger average dwell time (Figure 5G). In contrast, the psi-2 sub-population for all other contexts were found to have the psi-2 population existing with <30% of the events (Figure 5G). The Ψ impact on the dwell time while residing in the helicase active site was observed consistently in the redundant sequence contexts studied (Figure S10). The ability to use dwell time differences when Ψ passes through the helicase active site provides another means for base modification sequencing that does not rely on the protein nanopore sensor.

The dwell time signature found for Ψ provides two advancements for sequencing modifications using the nanopore setup: (1) This minimizes the window in which to call a modification down to 2 nts, and (2) it provides a secondary approach to call modifications that is not reliant on error-prone signals from the protein nanopore sensor that were found to be sequence-context dependent (Figures 2 and 3). The long-range dwell analysis was always present for Ψ, and when combined with base calling error analysis, it results in greater confidence to call modifications and their locations in the nanopore sequencing data.

Nanopore sequencing of RNAs with U or Ψ within biologically relevant contexts found differences in the bases called (Figure 2), differences in base-calling errors (Figure 3), differences in the currents and dwell times when the sites resided in the nanopore sensor (Figure 4), and differences in the dwell times when the sites passed through the active site of the helicase (Figure 5). Only the long-range dwell time differences provide the ability to detect Ψ consistently in all sequence contexts, albeit with a weaker signal. We propose using a hybrid analytical approach of base-calling errors from nanopore sensor-derived signals using Eligos2 analysis coupled with dwell time signatures derived from helicase activity differences using Nanocompore/Nanopolish analysis to permit high confidence calling of Ψ in RNAs.

This proposed approach was applied to publicly available nanopore sequencing data deposited for the SARS-CoV-2 RNA genome.^40,41^ In the deposited data, the modified RNA was obtained from SARS-CoV-2 infected cells, and the non-modified matched control was generated by IVT.^40,41^ The base-called fastq files were analyzed with Eligos2 to locate sites of base-calling errors between the samples, and the fast5 files were analyzed with Nanocompore/Nanopolish to inspect for currents and dwell time differences in the nanopore sensor and helicase proteins. A noteworthy point about coronaviruses is that during replication of their genomes, large populations of sub-genomic RNAs (sgRNAs) are generated, in which these shorter RNAs code for conserved structural proteins (spike protein [S], envelope protein [E], membrane protein [M], and nucleocapsid protein [N]), as well as for key accessory proteins (3a, 6, 7a, 7b, 8, and 10).^40^ The analysis described in the text to locate Ψ inspected the sgRNAs or transcriptional regulatory sequences (TRSs) for the structural proteins S (TRS-S), M (TRS-M), and E (TRS-E), as well as the accessory protein 3a (TRS-3a), which are the longest of the population of sgRNAs. One more noteworthy point is that homopolymer runs can yield signals that masquerade as modifications,^27,45^ and therefore, the SARS-CoV-2 analysis described removed homopolymer runs >4 nts; these sites may be modified^40^ but were removed because of the known issues with the sequencer that were verified in the RNAs studied (Figure S11).

Using TRS-S (length = 8407 nt) as an example, the base-calling error analysis to report oddR values identified 111 sites with a value ≥ 3 based on the data herein (Figures 3, 6A, and S12). The current analysis of TRS-S found 810 sites with a -log(*P*-value) threshold ≥ 50 and 125 sites with a threshold set to ≥ 100, a value selected to match the number of modified sites found with the base calling error analysis (Figures 6A and S12). The nanopore data analysis approach described in the present work found five high confidence Ψ sites in TRS-S, five in TRS-3a, five in TRS-E, and five in TRS-M (Figures 6C red labels; and S12-S15). In TRS-3a, -E, and -M, five of the identified peaks were at the same location (U27,164, U28,039, U28,759, U28,927, and U29,418); therefore, TRS-S was inspected at those positions to find weak base calling error signals (oddR < 3) and weak long-range dwell times for Ψ that support modification at all of these sites except U28,039 with lower confidence (Figure 6C gray labels). The analysis found U nucleotides in the SARS-CoV-2 TRSs that are modified to Ψ with conservation through the sub-genomic RNAs.

**Figure 6.**
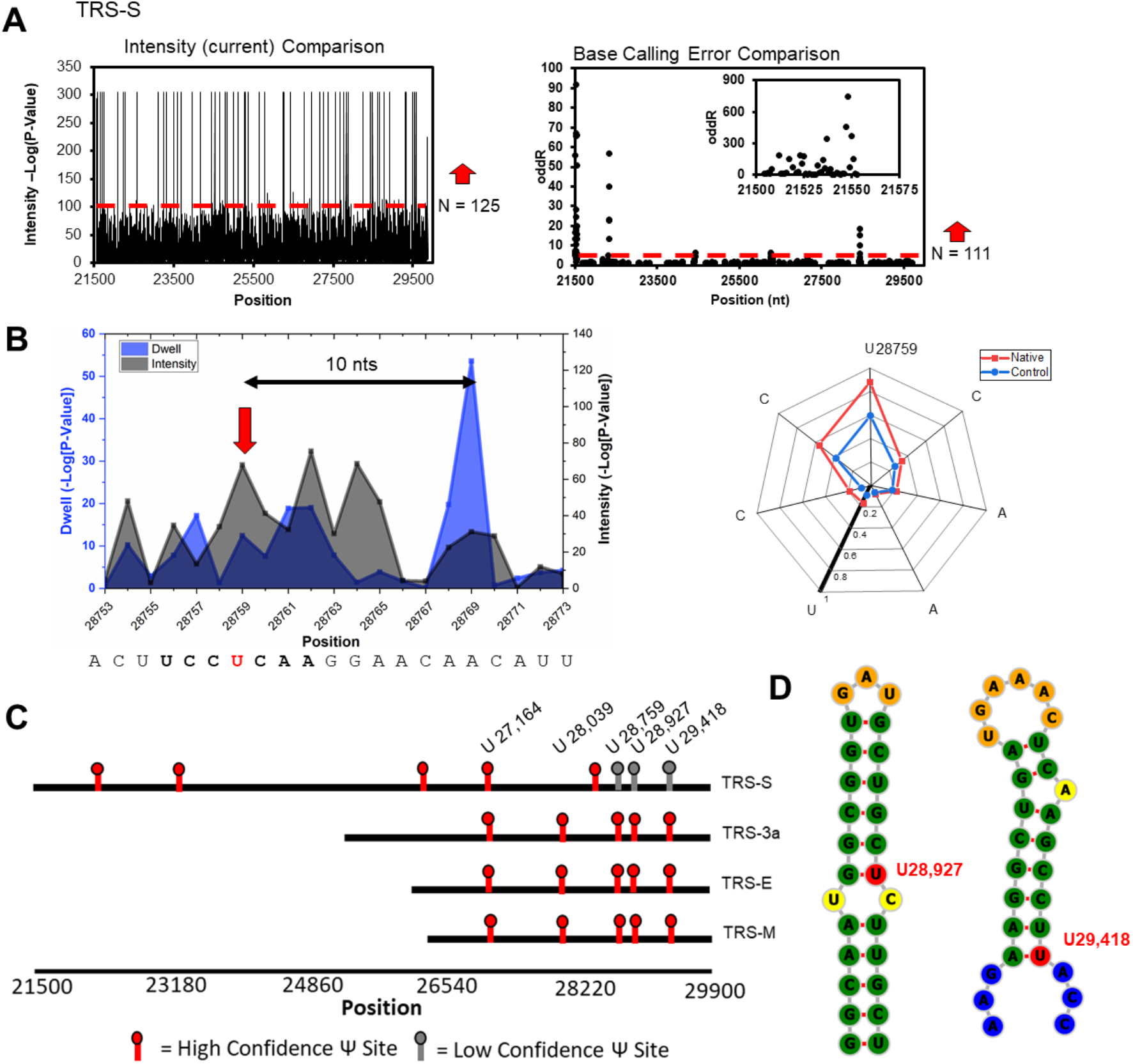
Analysis of nanopore sequencing reads for the SARS-CoV-2 RNA sub-genomes for Ψ. (A) Interrogation of the SARS-CoV-2 RNA extracted from cell culture with modifications against an IVT-generated genome without modifications to find statistically significant differences in current intensity and base calling error for the TRS-S sub-genome. (B) As an example, U27,164 in TRS-S yields a base-calling error found by Eligos2 and a long-range dwell signature found by Nanocompore/Nanopolish analysis. (C) Plot illustrating the Ψ sites found in the analysis. (D) RNAfold analysis of the region flanking U28,927 and U29,418 showing the local secondary structure is favorable as a substrate for PUS1 in comparison to a literature report.^44^ Data for the other sub-genomes is provided in Figures S11-20 and the full RNAfold analysis is found in Figure S21.

The knowledge of these five conserved sites of Ψ, led us to inspect the other TRSs in SARS-CoV-2 to determine whether their occupancy spanned all sub-genomes. The subgenomes progressively decrease in length, and therefore, if the sequence had the conserved Ψ site it was found to be modified in every TRS (Figure S16-S20). The sequence contexts for the five identified sites were U27,164 = 5’-AG**X**GA, U28,039 = 5’-AG**X**AG, U28,759 = 5’-CC**X**CA, U28,927 = 5’-GC**X**CU, and U29,418 = 5’-CU**X**AC (Figure S21); in the first two sites of the list, they have the 5’-GΨ context that likely gave strong dwell time signatures permitting their detection, and the other three had a 5’-pyrimidine that in general produced stronger base-calling errors (Figure 3). To reiterate, the analysis of the SARS-CoV-2 RNA sequencing data using nanopores identified five conserved pseudouridinylation sites on the 3’ end of the genome fragments. The base-calling error and dwell time analysis hybrid approach enabled this discovery out of the noisy nanopore data. The hybrid analysis approach did find other nucleotides that could be modified but they were not explored further because standards for these were not studied herein (Figures S11-20); however, there are two noteworthy examples of possible m^6^A modification sites in its established consensus motif found at A24,420 in TRS-S and A27,334 in TRS-3a; whether these are bonified modification sites is not known.

## Discussion

Direct RNA sequencing with nanopores has the potential to locate epitranscriptomic modifications via current levels, dwell times, and the associated base-calling errors. At present, the signatures for all modifications are not known and studies are needed to address what the signals are, as well as what the biases and limitations are to the data analysis. In the present work, Ψ was synthetically incorporated by IVT in RNA at known locations in 18 different human-relevant sequence contexts found in rRNA, mRNA, and tRNA (Figure S1). The Ψ sites were spaced >25 nt apart to study them one at a time as they pass from the helicase to the nanopore sensor, an overall distance that spans ~17 nt from the entry of the helicase to the exit of the k-mer sensing zone in the protein nanopore. Pseudouridine is base called by Guppy predominantly as U or C, consistent with other studies,^26^ and the present work found the ratio is dependent on the local sequence context (Figure 2). The base-calling error for Ψ was greater than U permitting detection of the modification (Figure 3); however, the base-calling errors were sequence context dependent, similar to the base-calling differences. Two extreme examples found in the data illustrate the challenges in using base-calling data for RNA modification sequence; in the 5’-AC**X**CA (**X** = U or Ψ) context, the U:C ratio is 7:88 with a base-calling error analysis giving an oddR value of 233, while the 5’-CA**X**CG context had a U:C ratio of 90:10 and an oddR value of 4, both when 100% Ψ is present at position **X** (Figures 2 and 3). In real samples, this approach will systematically favor observation of high error and high C calling sites over those that fit a profile similar to the reactant U resulting in huge biases to the data especially at sites that are not quantitatively modified. This is a problem because Ψ can reside in all possible sequence contexts.

A similar sequence dependency was observed for Ψ when inspecting the raw currents and dwell times as the modification passed through the protein nanopore sensor (Figure 4); moreover, the position within the k-mer window for which Ψ impacted the current and/or dwell time to the greatest extent was dependent on the sequence context, resulting in a 9-nt ambiguity of the location of a modification in a real sample. These analytical approaches work for modifications like *N^6^*-methyladenosine that are favorably deposited in reproducible sequence contexts,^3,6^ but fail for Ψ that can exist in all possible sequence contexts.^44^

The observation that Ψ impacts the dwell time as it passes through the helicase active site compared to U alleviates some of the challenges for detection, especially when this analytical approach is used in tandem with other detection strategies, as we propose in the present work. In all sequence contexts, Ψ produced an observable signal not seen for U in the helicase, which is a slowing of the helicase processing activity by ≥ 3-fold for a subpopulation of the events (Figure 5). The longer dwell-time subpopulation distribution was greatest for 5’-GΨ sequence contexts leading these contexts to be the easiest in which to detect Ψ. Nonetheless, in all sequence contexts studied (Figure 5F), the signal was present yielding a positive signal to locate Ψ in the strand. Unlike the other methods that did not report consistent values on replicate studies and in the case of the nanopore sensor did not yield signals with single-nucleotide resolution, the helicase stalling leading to a dwell time signature detects Ψ within a 2-nt window created by the helicase active site. With appropriate controls for sequence contexts, this could provide quantitative information on the extent of epitranscriptomic modification at a suspected Ψ site.

Pseudouridine creates four differences in the RNA strand compared to U that likely led to the sequencing signatures observed: (1) Uridine has a hydrogen bond donor site at N3 while Ψ can hydrogen bond at N1 and N3, and both have the hydrogen bond acceptor sites at *O^2^* and *O^4^* (Figure 1A). The Ψ N1 hydrogen shows long-lived bonding with the phosphodiester backbone to introduce rigidity in duplex RNA, and the bond likely exists in single-stranded RNA albeit with a shorter lifetime.^46^ (2) The glycosidic bond angle for Ψ shows a slight *syn* preference while U adopts the *anti* conformation almost exclusively.^47^ (3) Pseudouridine is more hydrophilic than U, and (4) Ψ stacks with adjacent bases better than U.^48^ These physical differences likely result in the ability to differentiate Ψ from U in the nanopore sequencer. In the CsgG protein nanopore, calls of Ψ as U or C with sequence context dependency may result from the hydrogen bond and *syn/anti* conformation differences that impact interactions with the nanopore. We propose the U vs. C base-calling ratio is a result of the *syn* vs. *anti* conformation of the Ψ heterocycle. The *syn* vs. *anti* conformational distribution shows strong sequence context dependency as observed in the wide range of U:C base calling ratios observed (Figure 2), and this provides a molecular understanding to the incoherent signal in the nanopore from this isomer of the U nucleotide.

As for the helicase, the literature suggests that a mutant form of the T4 phage Dda helicase is used in the ONT platform.^24^ The mechanism by which Ψ would slow Dda unwinding compared to U in the active site would be at an interaction with the base. Helicases such as Dda predominantly interact with nucleic acids via the backbone, although one π-stacking interaction between Phe97 and the base occurs,^23^ assuming this amino acid was not mutated in the helicase used. All four Ψ-U differences discussed may contribute to the helicase differentiation of Ψ, while the local rigidity imposed by N1-H of Ψ and its better π stacking are likely the dominant forces leading to the slower helicase processing kinetics. Further, the 5’ G effect likely occurs from more stabilized pi stacking that can compete with Phe97 to slow the helicase translocation along the RNA strand. Lastly, the reason why Ψ within the helicase active site results in two different average time populations is again a consequence of the *syn/anti* conformation distribution found for this heterocycle. This observation suggest one face of Ψ π stacks better with Phe97 than the other face. Details of the helicase mutations are needed to better address the Ψ vs. U difference that enabled U and Ψ differentiation.

The helicase stalling at Ψ permitted analysis for this modification in the noisy sequencing reads for the SARS-CoV-2 RNA genome with greater confidence. Focusing on the sub-genomic TRSs, there exist five conserved Ψ sites on the 3’ end of the RNA sub-genomes (Figures 6C and S21). The structure of RNA guides where Ψ is installed by pseudouridine synthases (PUSs) that are stand-alone enzymes such as PUS1, PUS7, and TRUB1.^14–16^ Each of the five sites with 50-nts of flanking native sequence were selected and submitted for RNAfold analysis to identify their predicted folding (Figure 6D and S21).^49^ The predicted folds for regions around U28,759, U28,927, and U29,418 have folds that place the U at the base of a hairpin or in a bulge, which are structures previously found to be PUS1 substrates (Figure 6D and S21).^44^ As for the other two sites, U27,164 is in a large single-stranded loop and U28,039 is near the middle of a long duplex RNA adjacent to a G:U wobble base pair (Figure S21). Studies on cells infected with SARS-CoV-2 while knocking down or out the PUSs are needed to truly define the Ψ writer(s) in the viral RNA.

Inspection of other RNA viruses (e.g., the flaviviruses Zika and HCV) by LC-MS have found Ψ exists at ~1-2% of the U nucleotides;^50^ the occupancy of Ψ in SARS-CoV-2 is not known at present but may exist at a similar level as the flaviviruses considering their similar replication cycle and RNA-based genomes. Selecting TRS-E as an example, the hybrid approach used to locate possible Ψ nucleotides found five that represents 0.5% of the U nucleotides in this sequence; in contrast, Eligos2 calls 3.7% of the U nucleotides as modified (oddR ≥ 3) and Nanocompore found 6.1% of the modified k-mers had a U nucleotide defined as a (-log(*P*-Value) ≥ 100 (Figure S13). The Nanocompore results report on 5-nt k-mers, so the high value likely exists because of other chemical modifications in the genome. The oddR approach of modification calling by Eligos2 implies there will be false positives in the data set, and therefore, inflating the number of modified nucleotides. The approach herein is most likely an underestimate but reports on those sites that give complementary positive signals, and therefore, these are sites most anticipated to be modified at high levels and have the greatest likelihood of biological significance.

High occupancy Ψ sites in the SARS-CoV-2 RNA genome may have a biological function. The pseudouridinylation of viral RNA is hypothesized to be a mechanism by which the virus hijacks host enzymes to avoid the immune response. Support for this is the use of Ψ or its N1-methyl derivative for this purpose in mRNA vaccines,^51^ as well as the possibility that Ψ serves the same function in HIV and flaviviruses.^52,53^ If Ψ is used by SARS-CoV-2 and possibly other coronaviruses during the infection cycle to minimize immune stimulation, this points to new schemes for intervention;^54^ additionally, if Ψ is found to be essential for infectivity of the virus, an inspection of host genetic variants for pseudouridine synthase enzymes that impact activity may reveal more clues as to why host outcomes from SARS-CoV-2 infection are variable beyond other comorbidity factors.

## Conclusions

The epitranscriptome is currently at center stage for the discovery of critical details regarding RNA regulation in biology and is being fueled by advancements in sequencing technology. A nanopore sequencing platform comprised of two proteins, a nanopore sensor and a helicase brake, enabled us to directly sequence RNA for Ψ, which is the most common RNA modification. Two key sequencing features were identified: (1) The protein nanopore sensor produced a wide range of signals for Ψ with dependency on the sequence context and position in the 5-nt sensing zone of the CsgG protein (Figures 2, 3, and 4). Some contexts gave robust signals and others were nearly in the noise that would yield false negatives when inspecting unknown samples. (2) The helicase employed to regulate the speed of translocation was found to have sensing capabilities for Ψ by stalling on the modification but not the parent U. The stalling results in a long-range dwell signal 10-11 nt 3’ to the protein nanopore as the modification passes through the helicase active site (Figure 5). This signal was always present but most pronounced in 5’-GΨ sequence contexts. Knowledge of base-calling errors and helicase dwell-time signatures permitted the analysis of the SARS-CoV-2 RNA sub-genomes for Ψ (Figure 6). Analysis of the viral RNA identified five conserved Ψs on the 3’ end of the fragments. The local structures for three of the modified sites are similar to those previously identified as PUS1 sites (Figures 6D and S21);^44^ however, the writer enzyme(s) is/are not known. Using the literature as a guide, we propose these Ψs are beneficial to the virus by aiding in avoidance of the immune response to favor replication.^51–53^ The findings herein expand our knowledge of the viral epitranscriptome regarding Ψ that can be exploited for future interventions and understanding of individual host responses to SARS-CoV-2 viral infection.

## Data Availability

The data are available upon request.

## Supplementary Data

Supplementary Data are available.

## Acknowledgments

The authors are thankful to Dr. Yun Ding and Ms. Andrea Taillacq for their assistance with the computational tools.

