## Supplemental Data for "Nanopore dwell time analysis permits sequencing and conformational assignment of pseudouridine in SARS-CoV-2"

**Item** **Page**

**Figure S1.** Sequences of RNA synthesized with site specific Ψ. S3

**Figure S2**. Read statistics obtained from MultiQC analysis of the strands studied. S5

**Figure S3**. Reproducibility analysis of base calling of the same sequence context. S6

**Figure S4**. Additional Eligos2 data for sequencing Ψ in synthetic RNAs with known modification sites. S7

**Figures S5**. A plot of Eligos2 reproducibility for oddR values. S10

**Figure S6**. Tombo current-level analyses for Ψ (red) vs. U (black). S11

**Figure S7**. Additional plots for the current and dwell time analysis using Nanopolish and Nanocompore. S14

**Figure S8**. Reproducibility in the Nanocompore analysis. S20

**Figure S9**. Dwell-time histograms for U vs. Ψ in the helicase active site. S21

**Figure S10**. Reproducibility of the helicase dwell signatures. S24

**Figure S11**. Homopolymers can give long-range false positive signals in Nanocompore and Eligos2 analysis. S25

**Figure S11**. Data Analysis for TRS-S from SARS-CoV-2. S26

**Figure S12.** Data analysis for TRS-3a from SARS-CoV-2. S30

**Figure S13.** Data analysis for TRS-E from SARS-CoV-2. S35

**Figure S14.** Data analysis for TRS-M from SARS-CoV-2. S38

**Figure S15.** Data analysis for TRS-6 from SARS-CoV-2. S41

**Figure S16.** Data analysis for TRS-7a from SARS-CoV-2. S44

**Figure S17.** Data analysis for TRS-7b from SARS-CoV-2. S46

**Figure S18**. Data analysis for TRS-8 from SARS-CoV-2. S48

**Figure S19**. Data analysis for TRS-N from SARS-CoV-2. S50

**Figure S20**. Summary of the Ψ epitranscriptomic sites found in SARS-CoV-2. S52

**Figure S21**. Predicted folding for the region flanking the conserved Ψ sites in the TRSs from SARS-CoV-2. S53

**References** S54

**Figure S1.** Sequences of RNA synthesized with site specific Ψ.

**Strand 1**

5`GAGCACAGGACCAGAC**GCΨGC**ACAGAGCCGAAGCACAGCAGACCAGA**CCΨΨAΨCC**AG
AAGACGAGACCA**AAΨGA**CCAGAAGCCGAAGCACAGACGA**AAΨΨAG**CCAGACGGACAACA
GCAGAGACCGAAG**CGΨGG**GCAGACACGCAGCGACAGAGCAGCA**GGΨGA**GGACC**AGΨCA**
GGACAACAGAAAACAAAAAAAAAA

**Strand 2**

5`GAGCAGCACGAGACGA**GGΨGA**CACGACAGAGAGCGGACGC**AGΨCA**CGACCGACGAAC
ACGCA**GCΨGC**CAGACAAAGAGAACGCAGCACGA**CGΨAG**CGACGCAGACGGCGCAGCGA
G**CAΨAG**CACGCACGCAGCCACGCACAGAC**CGΨCG**CCAGCCGCAGCAGCACGACA**CAΨC**
**G**CGACGGCACGGAGCGGACGCACGACGAGCACAAAACAAAAAAAAAAAA

**Strand 3**

5`GAGCACAGGACCAGAC**GCΨGC**ACAGAGCCGAAGCAACCAGA**CCΨCC**AGAAGACGAGA
CCA**AAΨGA**CCAGAAGCCGAAGCACAGACGA**CGΨAG**CCAGACGGACACAGAGAAAG**AAΨ**
**CG**GCAGACACGCAGCGACAAAAGAGCCCGAGCAGCAGCA**ACΨCA**GGACGGACAGACAAC
CCCAGGCAAGAGCACGGCACAAAACAAAAAAAAAAAA

**Strand 4**

5`-AUAGAGUGCUAGUGUA**GUΨUC**AUACUCUGCAGAUACU**GUΨAA**CACAGAGAAAAAAAA

AAA

Ψ = pseudouridine, bold = site modified and adjacent sequence context, underlined sites are 4-nt repeats studied.

The sequences were also synthesized with the canonical form of the modified nucleotide present.

Sequence Contexts Studied Derived from Human RNAs

| Sequence Context | Location/Consensus |
|---|---|
| **Strand 1** | |
| GCΨGC | 5.8s rRNA 53-57 |
| CCΨΨAΨCC | 28s rRNA 1764-1771 |
| AAΨGA | 28s rRNA 1729-1733 |
| AAΨΨAG | 18s rRNA 812-817 |
| CGΨGG | tRNA Ψ-loop consensus |
| GGΨGA | 18s rRNA 404-408 |
| AGΨCA | tRNA site |
| **Strand 2** | |
| GGΨGA | 18s rRNA 404-408 (repeat) |
| AGΨCA | tRNA (repeat) |
| GCΨGC | 5.8s rRNA 53-57 (repeat) |
| CGΨAG | Potential mRNA PUS1 Site |
| CAΨAG | Potential mRNA PUS1 Site |
| CGΨCG | Potential mRNA PUS1 Site |
| CAΨCG | Potential mRNA PUS1 Site |
| **Strand 3** | |
| GCΨGC | 5.8s rRNA 53-57 (repeat) |
| CCΨCC | Potential mRNA PUS1 Site |
| AAΨGA | 28s rRNA 1729-1733 (repeat) |
| CGΨAG | Potential PUS7 Site (repeat) |
| AAΨCG | Potential RPUSD2 Site |
| ACΨCA | Potential RPUSD2 Site |
| **Strand 4** | |
| GUΨUC | Potential mRNA TRUB1 site |
| GUΨAA | Potential mRNA TRUB1 site |

There exist repeats of some of the sequence contexts in the pseudouridine-containing strands that were used to address whether position in the strand impacted the sequencing results. These sites are labeled with repeat in the location/consensus box.

**Figure S2**. Read statistics obtained from MultiQC analysis of the strands studied.

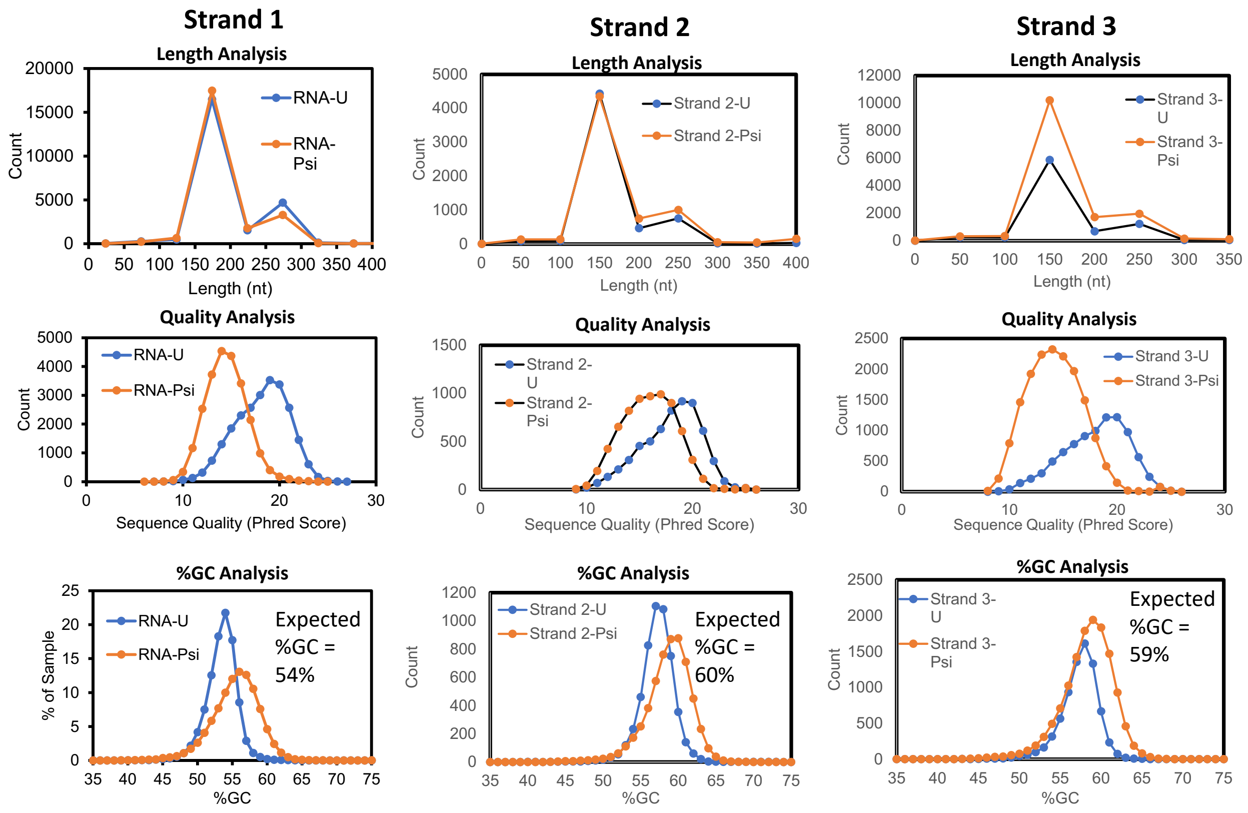


Strand 4 did not provide enough reads to conduct this analysis.

**Figure S3**. Reproducibility analysis of base calling of the same sequence context.

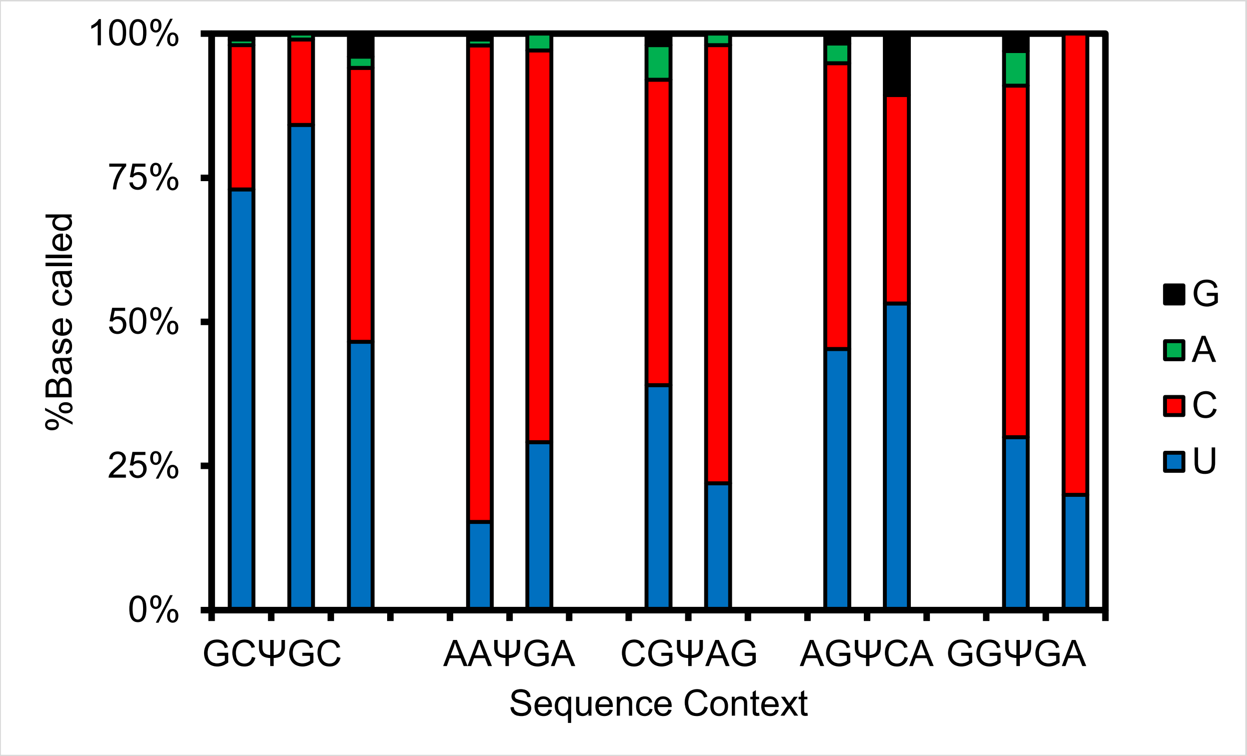


The GCΨGC context (left most data) was studied three independent times.

**Figure S4**. Additional Eligos2 data for sequencing Ψ in synthetic RNAs with known modification sites.

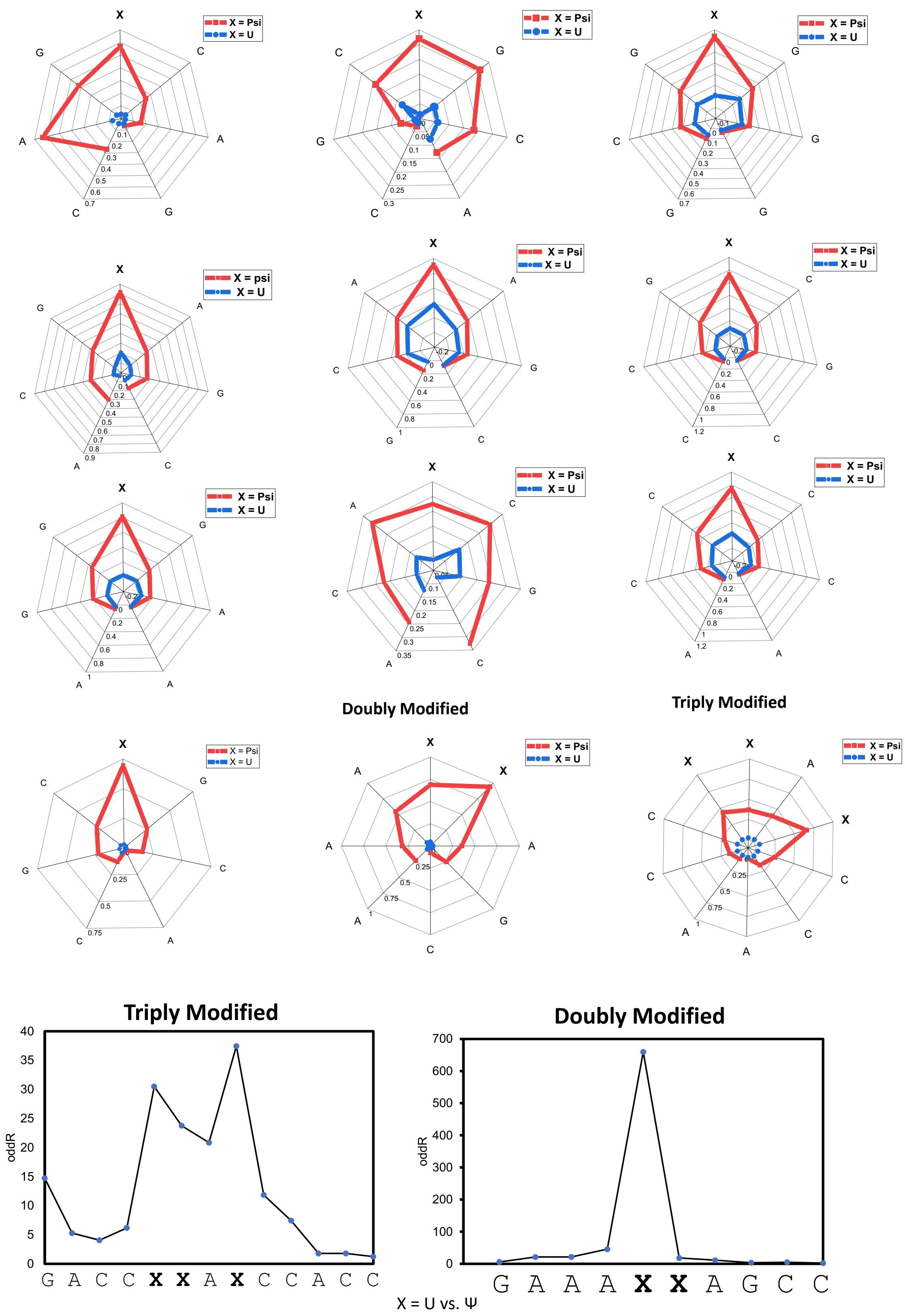

The plots below show Eligos2 data plotted as oddR or -log(*P*-value) to illustrate both tests give similar results, but the oddR values better define the position of the Ψ nucleotide.

### Strand 1 (X = U or Ψ)

#### Odds Ratio (oddR)

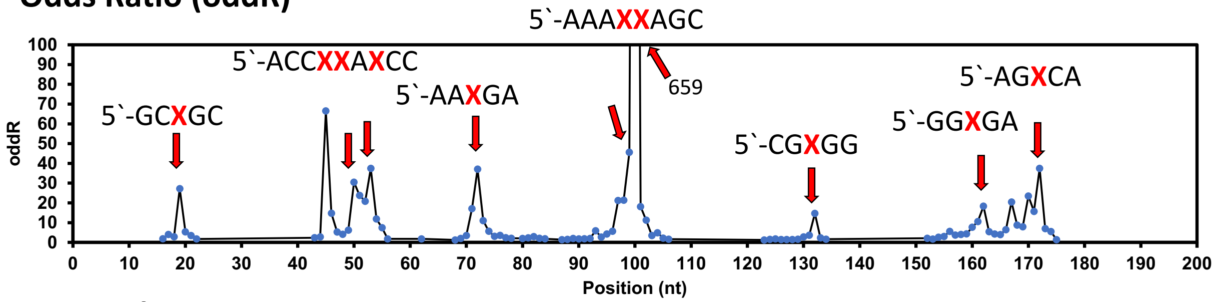


#### P-Values

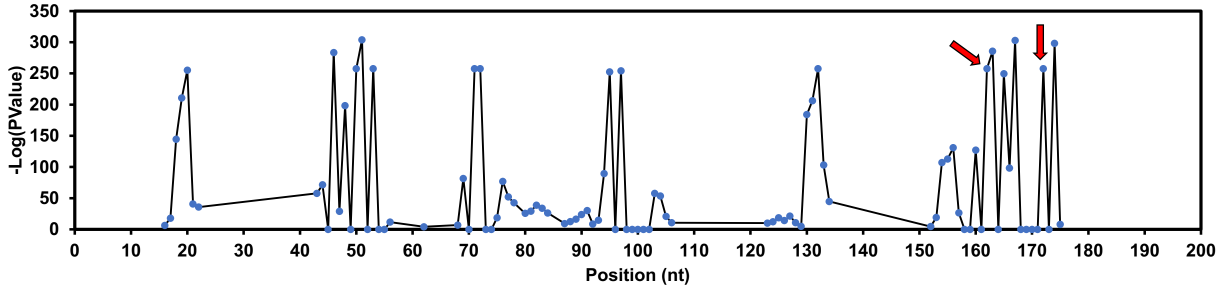


### Strand 2 (X = U or Ψ)

#### Odds Ratio (oddR)

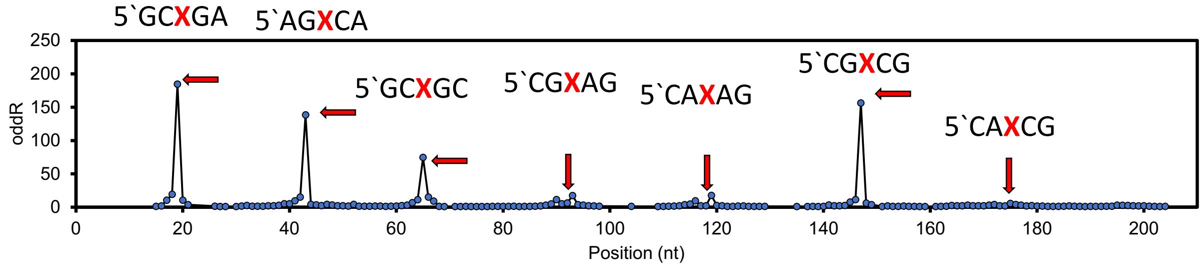


#### P-Values

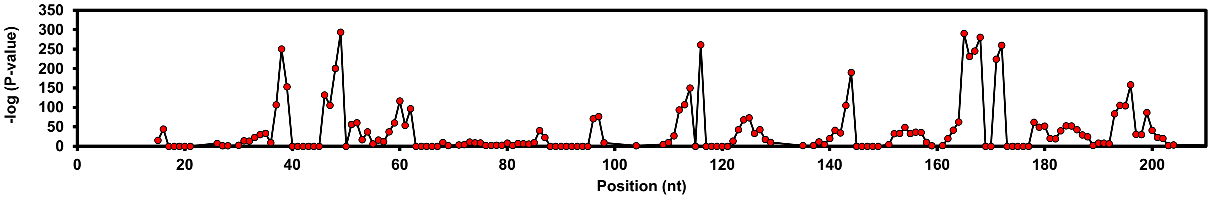

### Strand 3 (X = U or Ψ)

#### Odds Ratio (oddR)

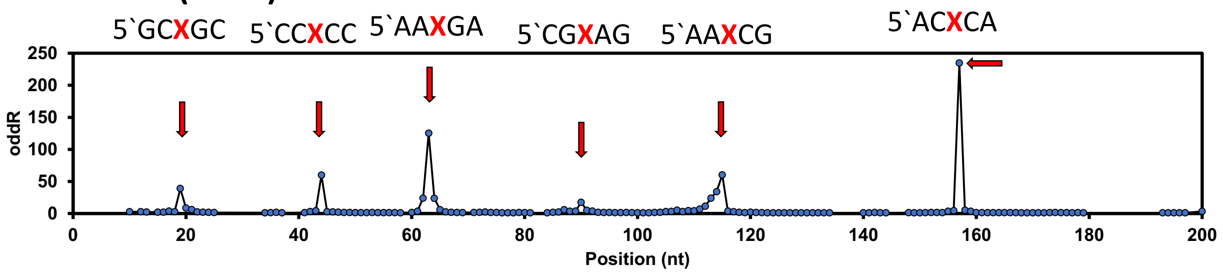


#### P-Values

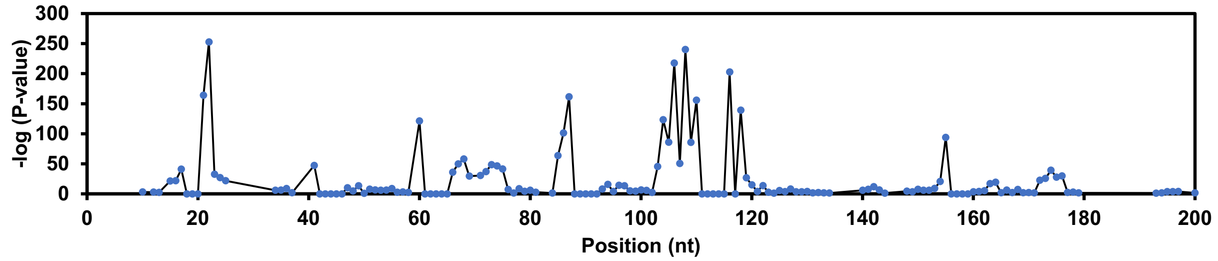

**Figures S5**. A plot of Eligos2 reproducibility for oddR values.

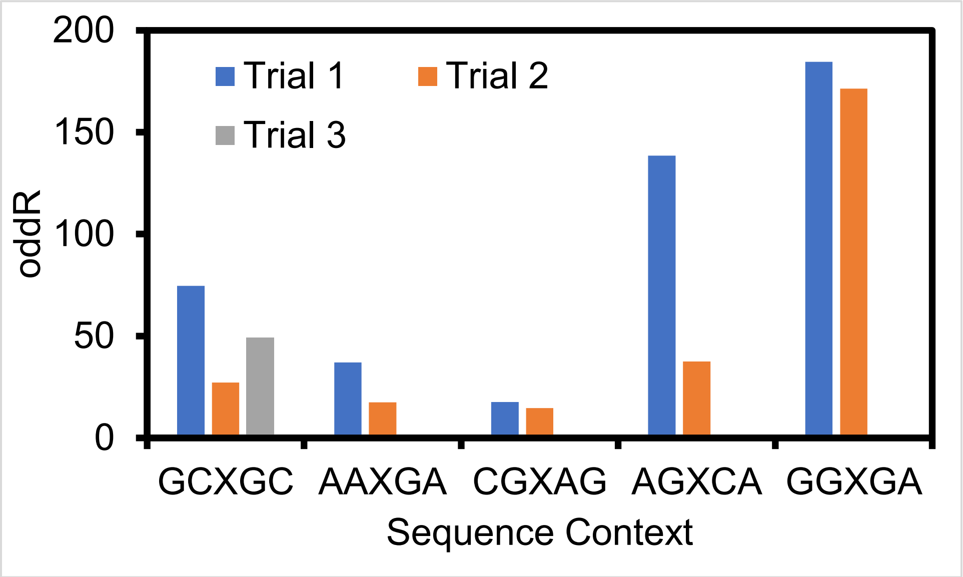


The sequence context GCXGC (where X = U or Ψ) was studied three independent times.

**Figure S6**. Tombo current-level analyses for Ψ (red) vs. U (black).

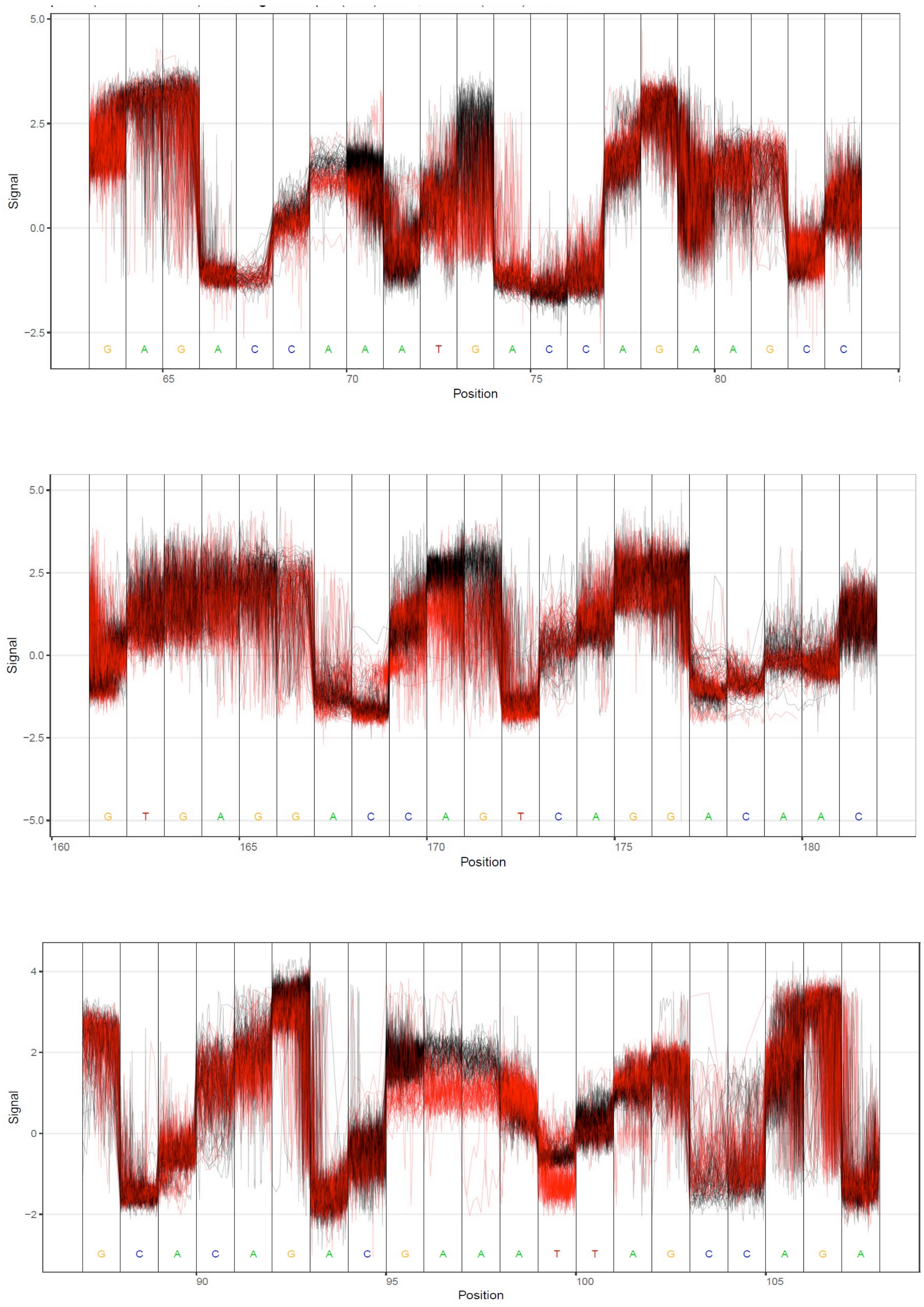

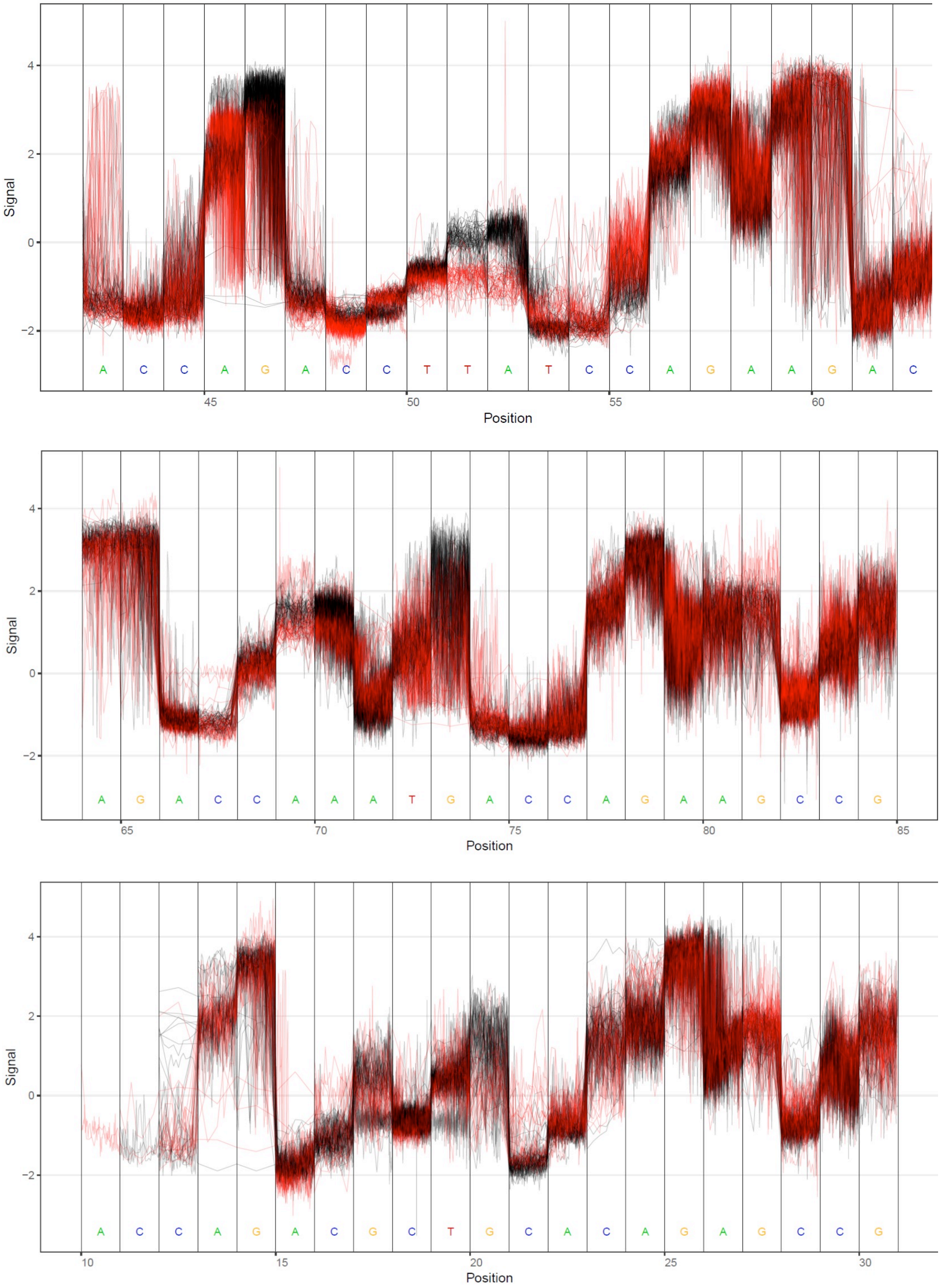
Signal
4
2
0
−2
A C C A G A C C T T A T C C A G A A G A C
45
50
55
60
Position
Signal
4
2
0
−2
A G A C C A A A T G A C C A G A A G C C G
65
70
75
80
85
Position
Signal
4
2
0
−2
A C C A G A C G C T G C A C A G A G C C G
10
15
20
25
30
Position

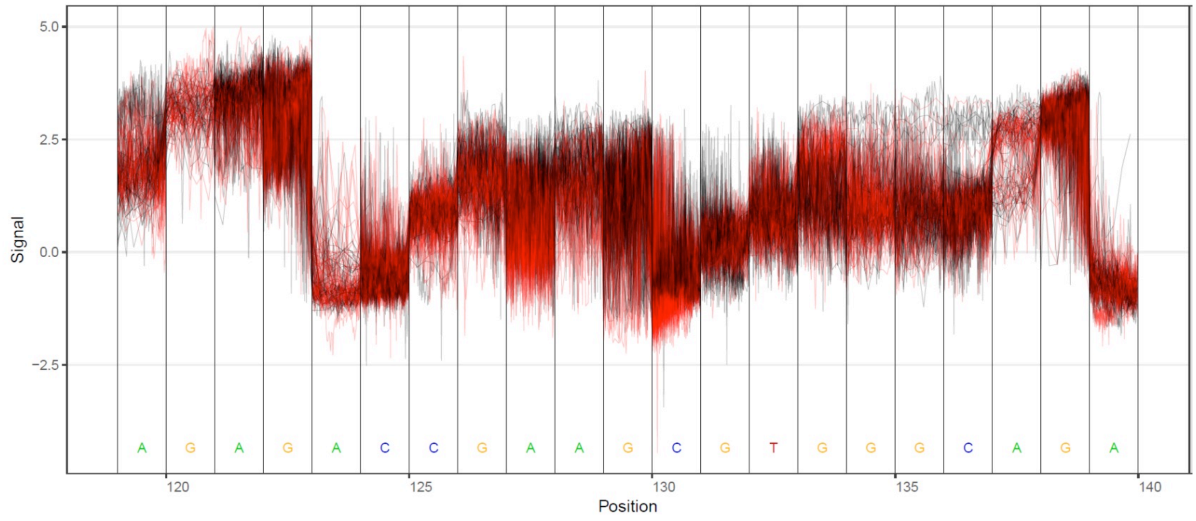
5.0
2.5
0.0
−2.5
Signal
A G A G A C C G A A G C G T G G G C A G A
120
125
130
135
140
Position

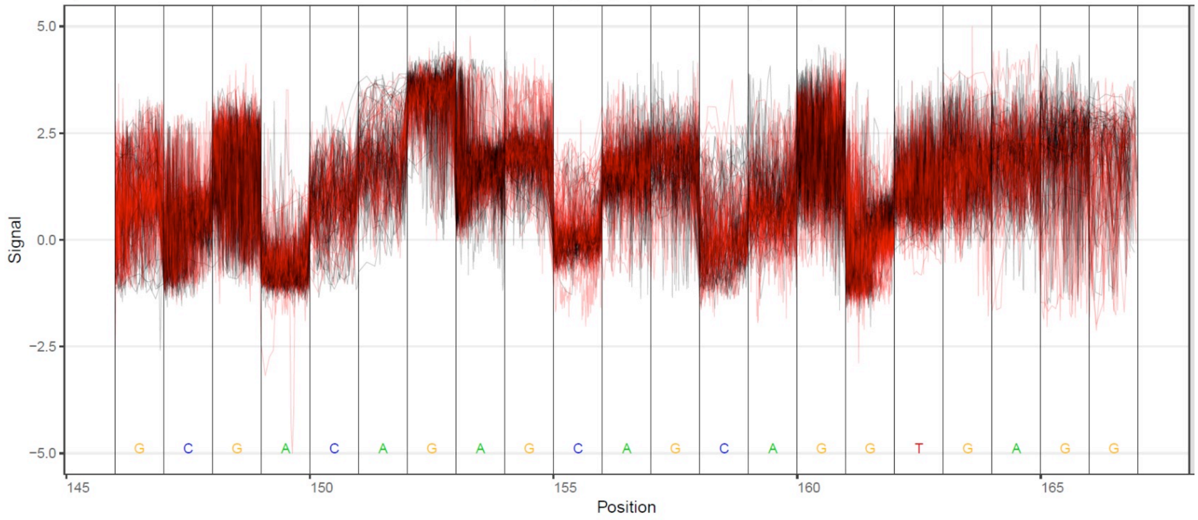
5.0
2.5
0.0
−2.5
−5.0
Signal
G C G A C A G A G C A G C A G G T G A G G
145
150
155
160
165
Position

**Figure S7**. Additional plots for the current and dwell time analysis using Nanopolish and Nanocompore.

The first block of data shows the passage of a Ψ (yellow) or U (dark blue) nucleotide through a 5-mer window of the sensor protein while monitoring the current intensity (top block of plots) or dwell times (bottom block of plots). These data were plotted from the Nanopolish eventalign files.

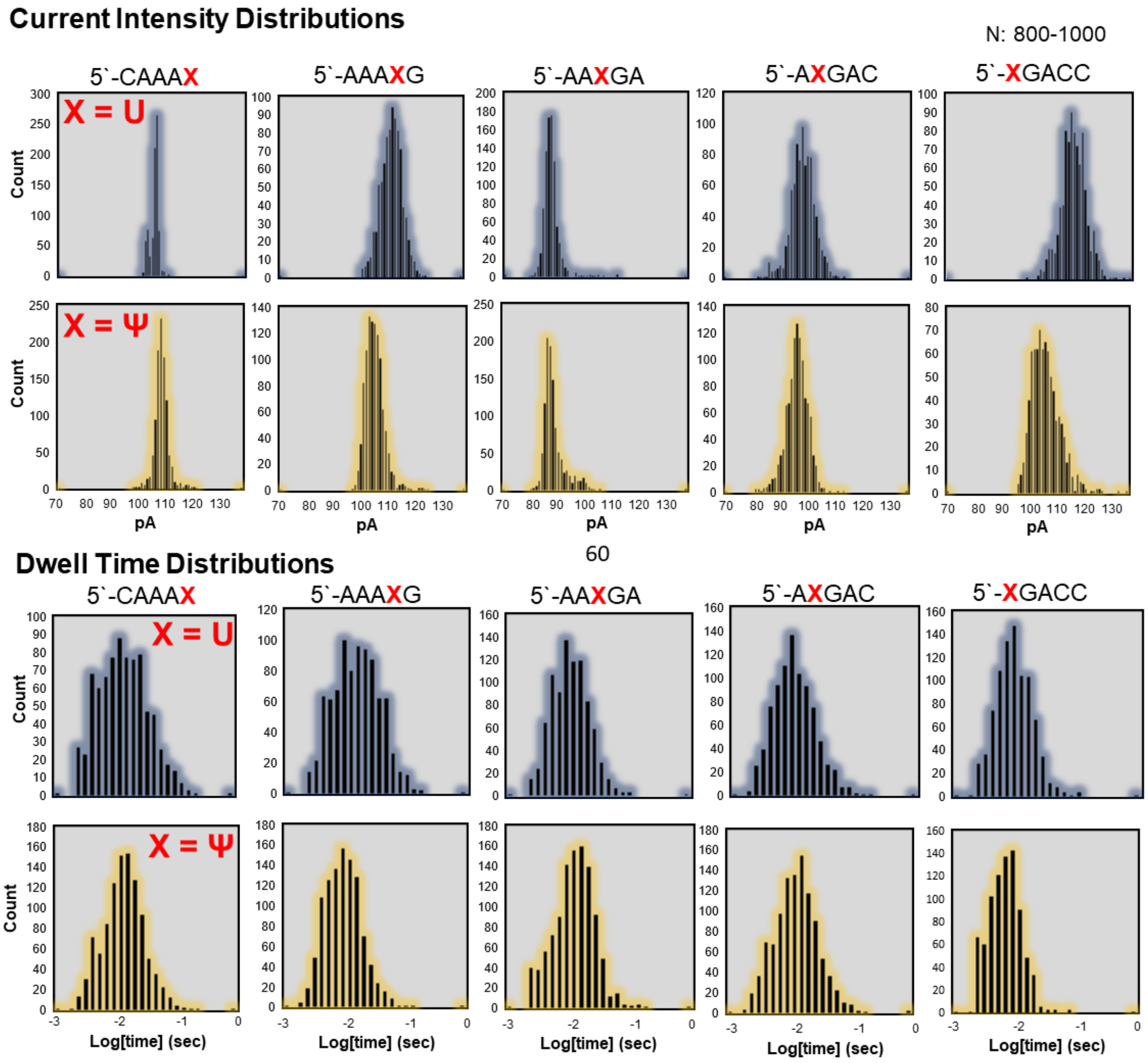

The following plots were created using Nanocompore to show the passage of U (aqua) or Ψ (light red) through the nanopore from the helicase to the sensor protein while monitoring current intensities (left block) or dwell times (right block).

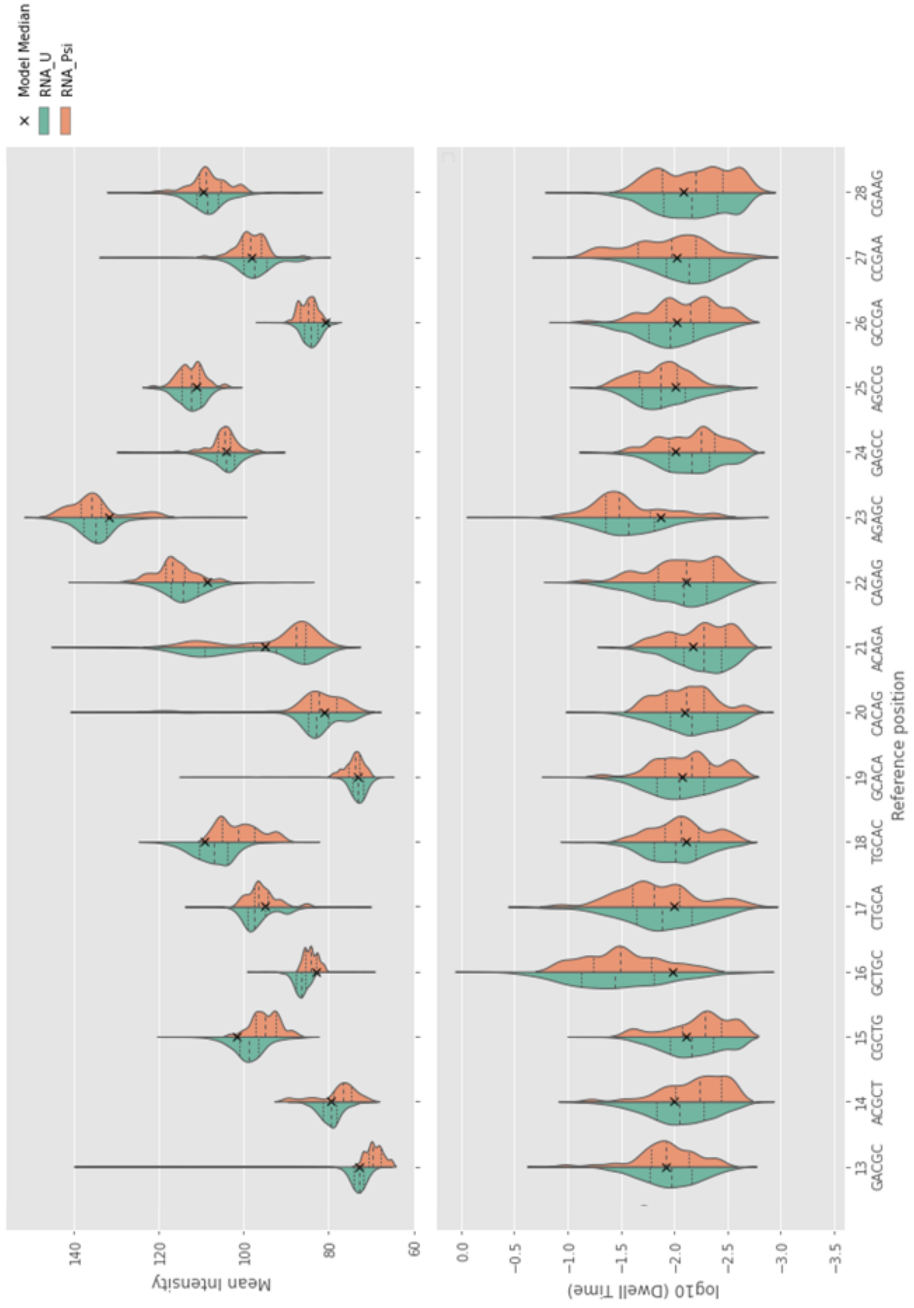

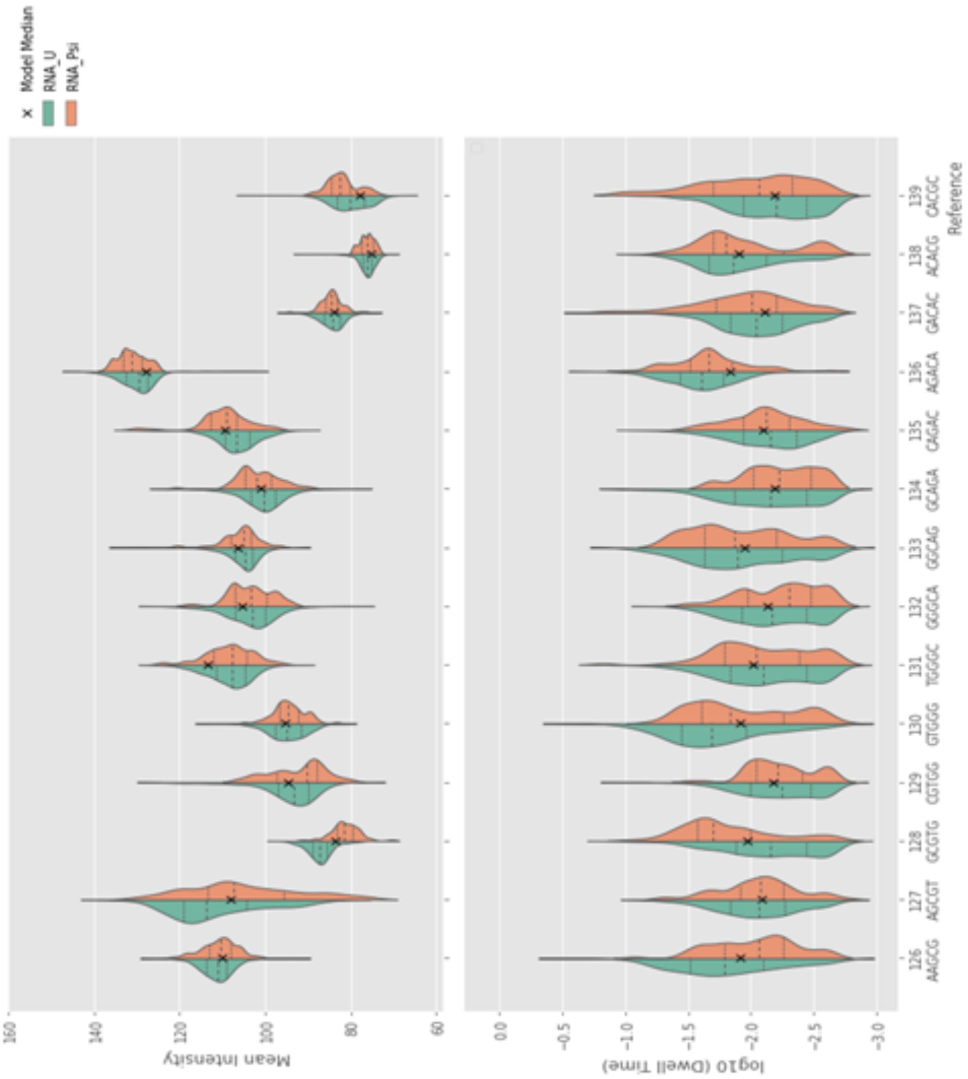
Model Median
RNA_U
RNA_Psi
Mean Intensity
log10 (Dwell Time)
Reference
126 AAGCG
127 AGCGT
128 GCGTG
129 CGTGG
130 GTGGG
131 TGGGC
132 GGGCA
133 GGCAG
134 GCAGA
135 CAGAC
136 AGACA
137 GACAC
138 ACACG
139 CACGC

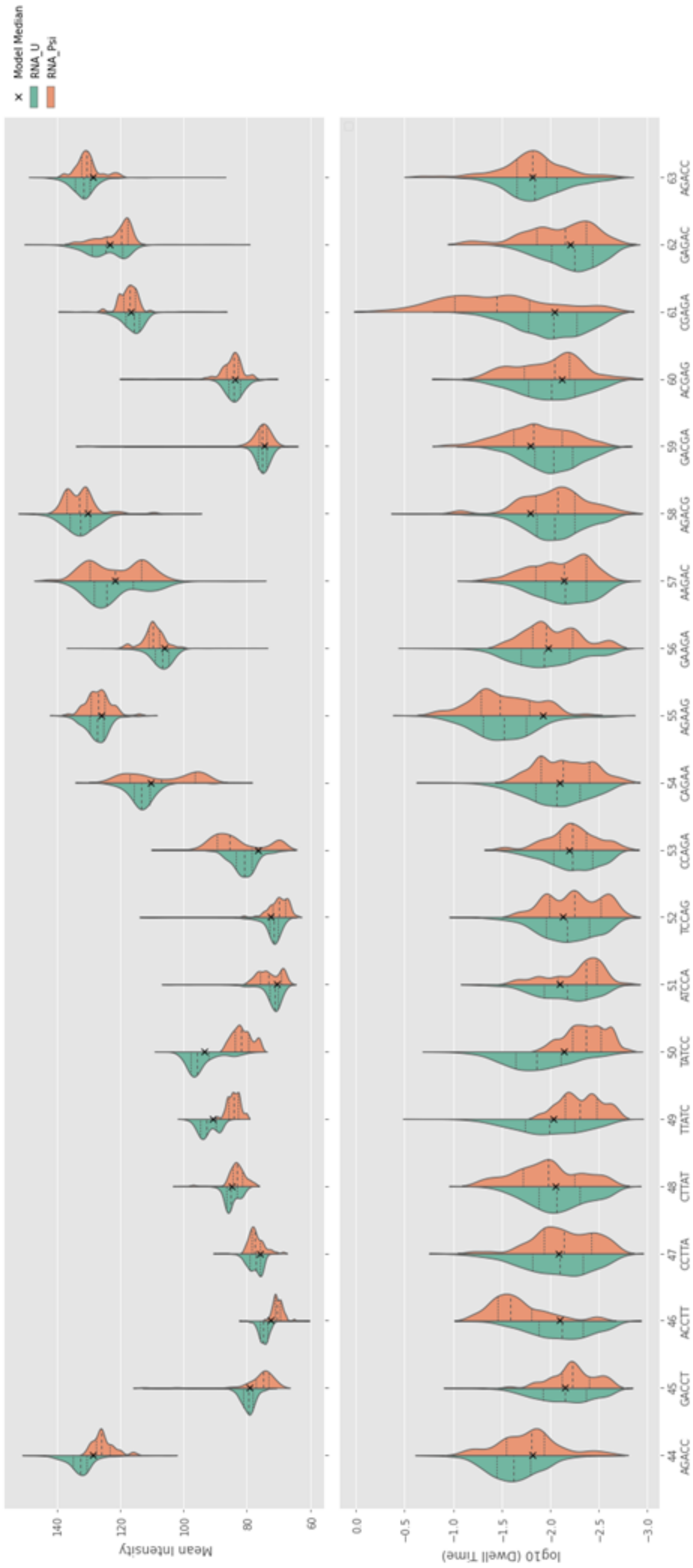

Model Median
RNA_U
RNA_Psi
Mean Intensity
log10 (Dwell Time)
Reference position
44 AGACC
45 GACCT
46 ACCTT
47 CCTTA
48 CTTAT
49 TTATC
50 TATCC
51 ATCCA
52 TCCAG
53 CCAGA
54 CAGAA
55 AGAAG
56 GAAGA
57 AAGAC
58 AGACG
59 GACGA
60 ACGAG
61 CGAGA
62 GAGAC
63 AGACC

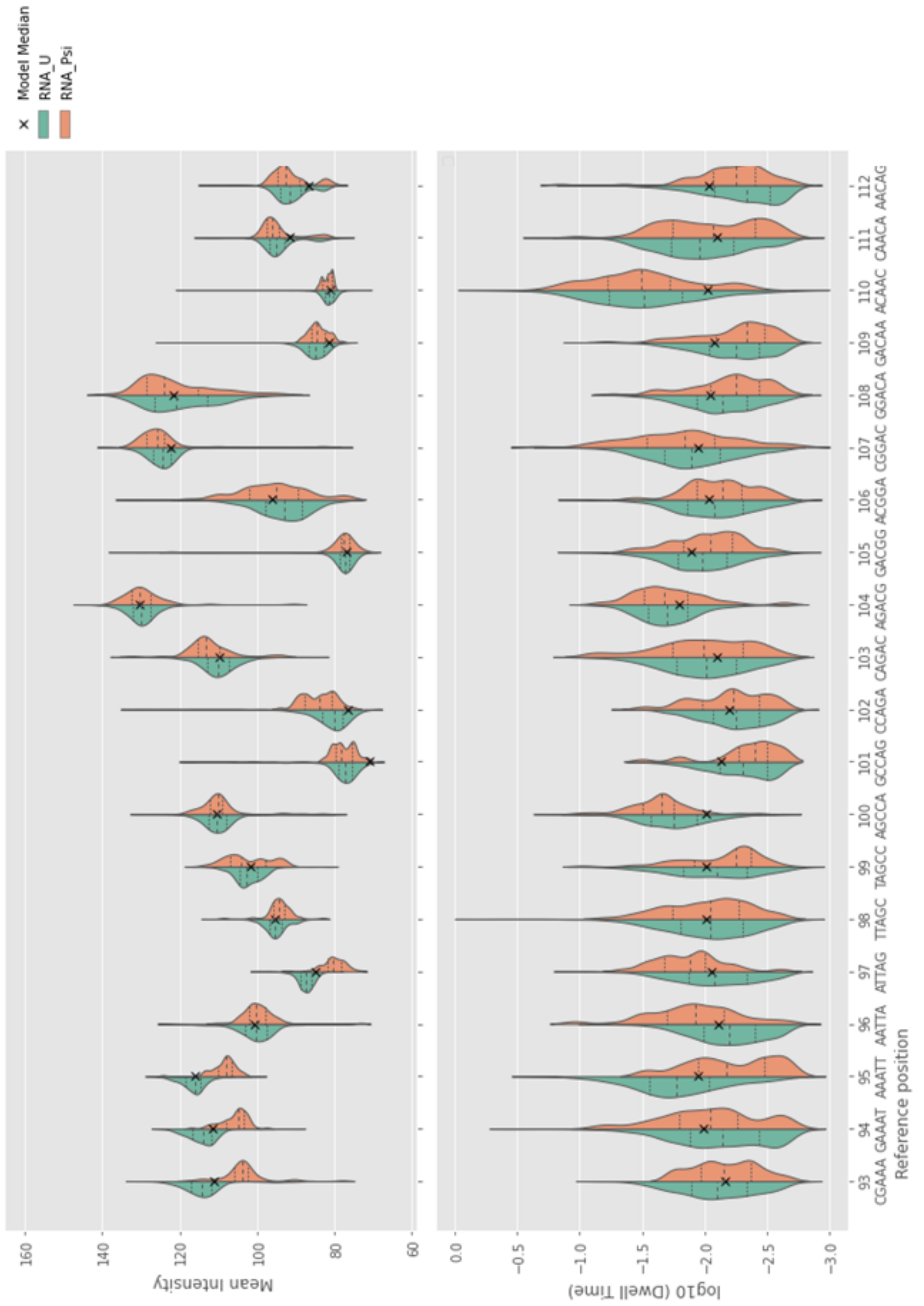

Model Median
RNA_U
RNA_Psi
Mean Intensity
160
140
120
100
80
60
log10 (Dwell Time)
0.0
−0.5
−1.0
−1.5
−2.0
−2.5
−3.0
93 CGAAA
94 GAAAT
95 AAATT
96 AATTA
97 ATTAG
98 TTAGC
99 TAGCC
100 AGCCA
101 GCCAG
102 CCAGA
103 CAGAC
104 AGACG
105 GACGG
106 ACGGA
107 CGGAC
108 GGACA
109 GACAA
110 ACAAC
111 CAACA
112 AACAG
Reference position

The plots below are for the Nanocompore statistical test results comparing RNAs with U or Ψ at defined positions to show where the greatest changes were (red arrows) and whether there was a long-range dwell time effect (black arrows) observed.

### Strand 1 (X = U or Ψ)

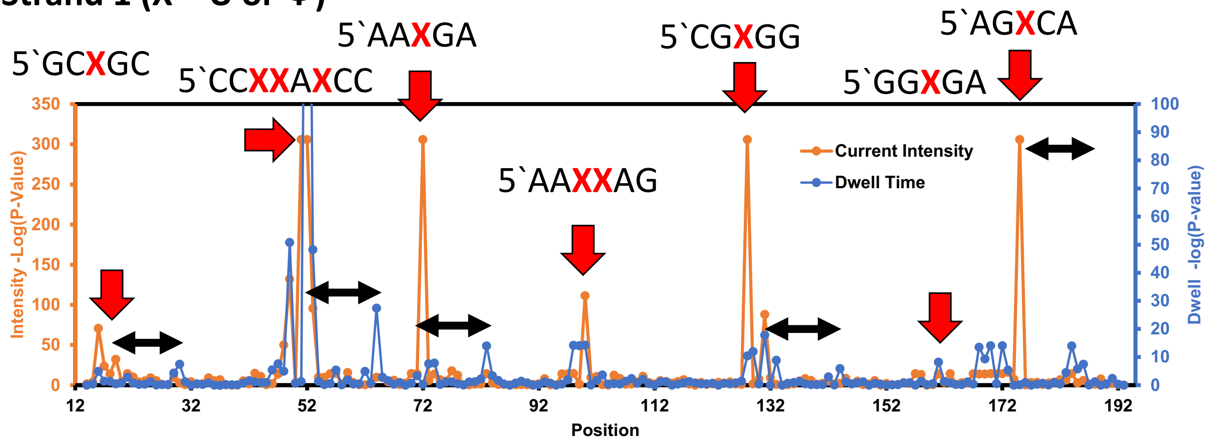


### Strand 2 (X = U or Ψ)

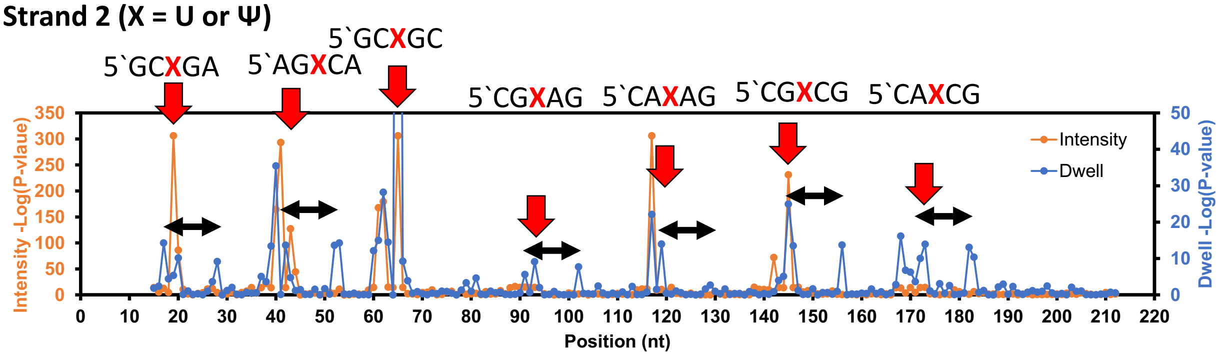


### Strand 3 (X = U or Ψ)

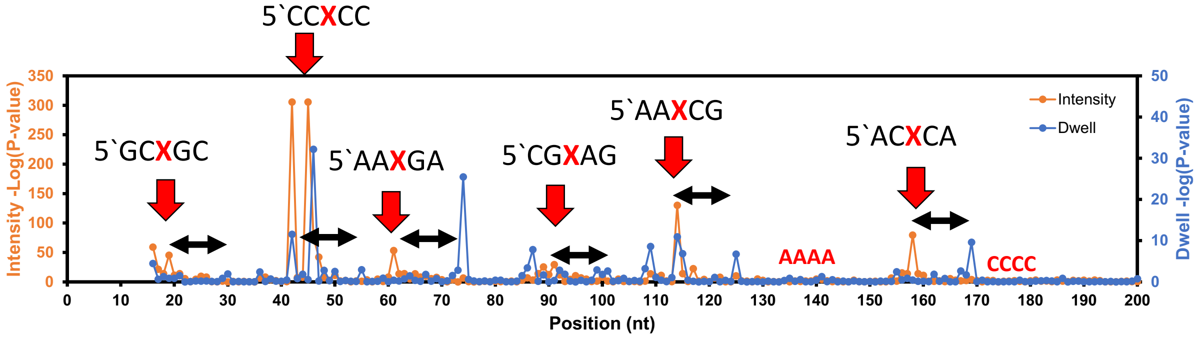

**Figure S8**. Reproducibility in the Nanocompore analysis.

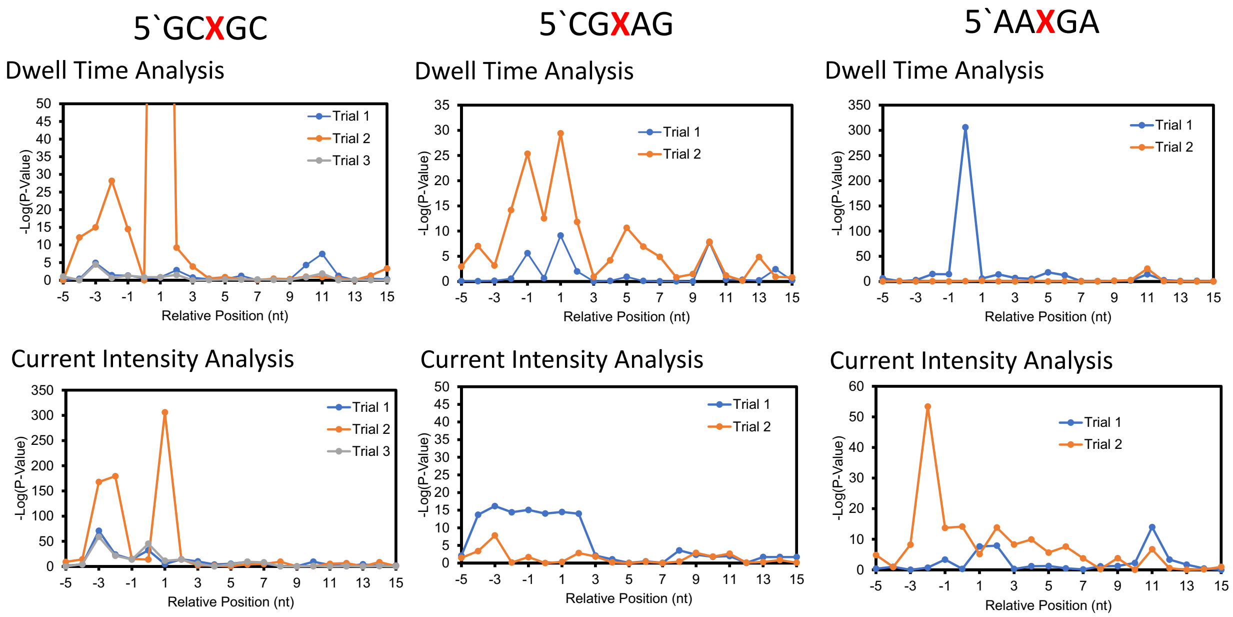

**Figure S9**. Dwell-time histograms for U vs. Ψ in the helicase active site.

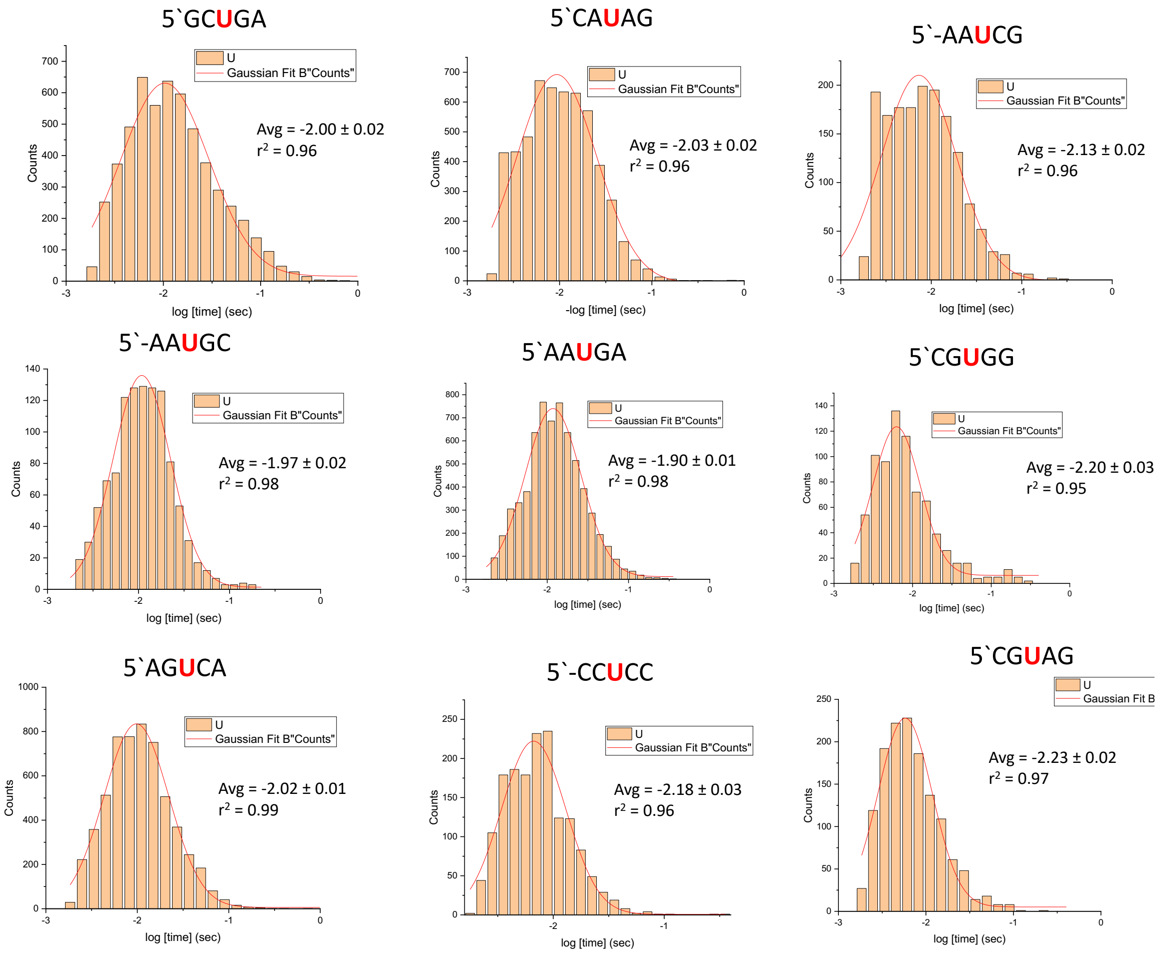

5`CG Ψ GG

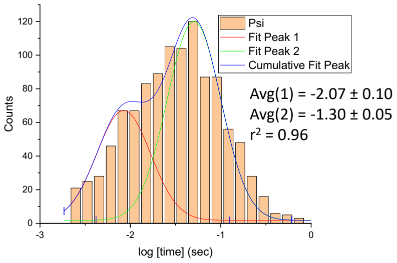


5`-CC Ψ CC

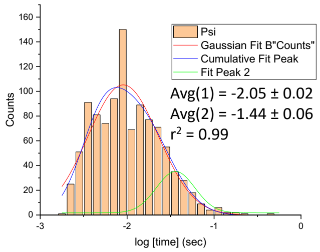


5`-AAΨGC

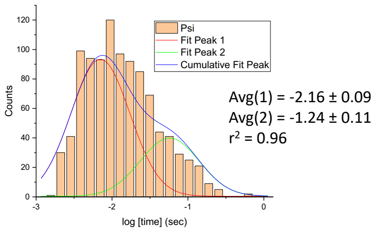


5`CG Ψ AG

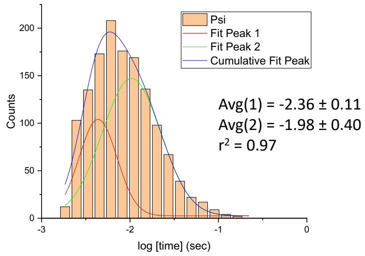


5`CG Ψ CG

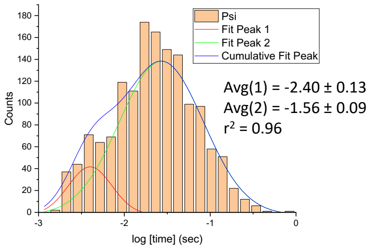


5`GC Ψ GA

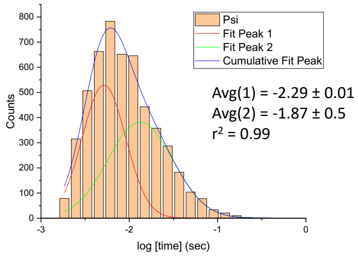


5`AG Ψ CA

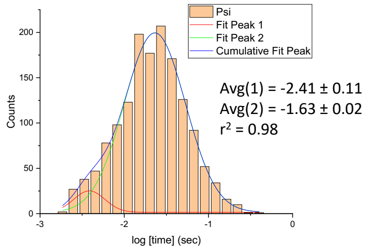


5`AG Ψ GA

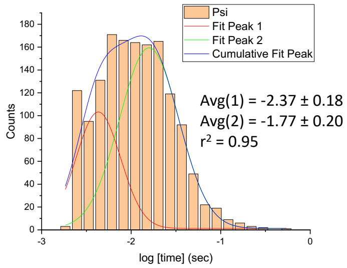


5`AA Ψ GA

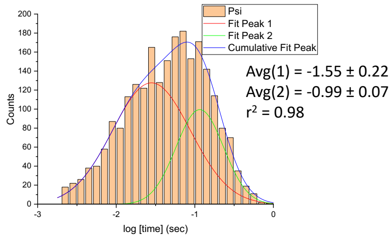

5`AA Ψ $_1$ Ψ AG

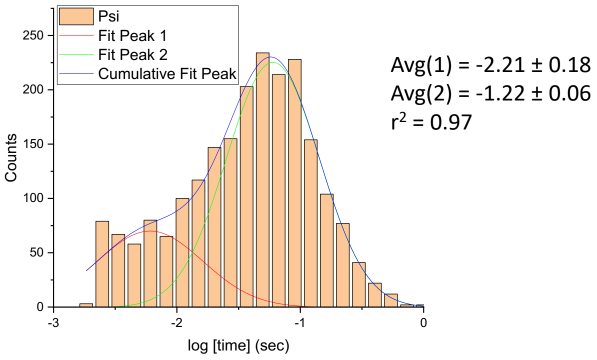


5`AA Ψ Ψ $_2$AG

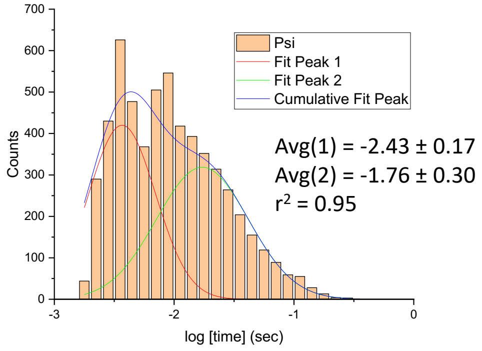


5`CCΨΨA Ψ CC

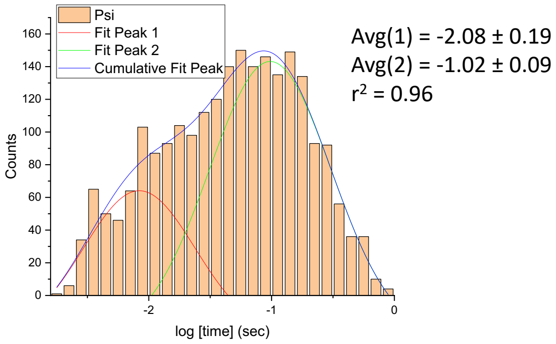

**Figure S10**. Reproducibility of the helicase dwell signatures.

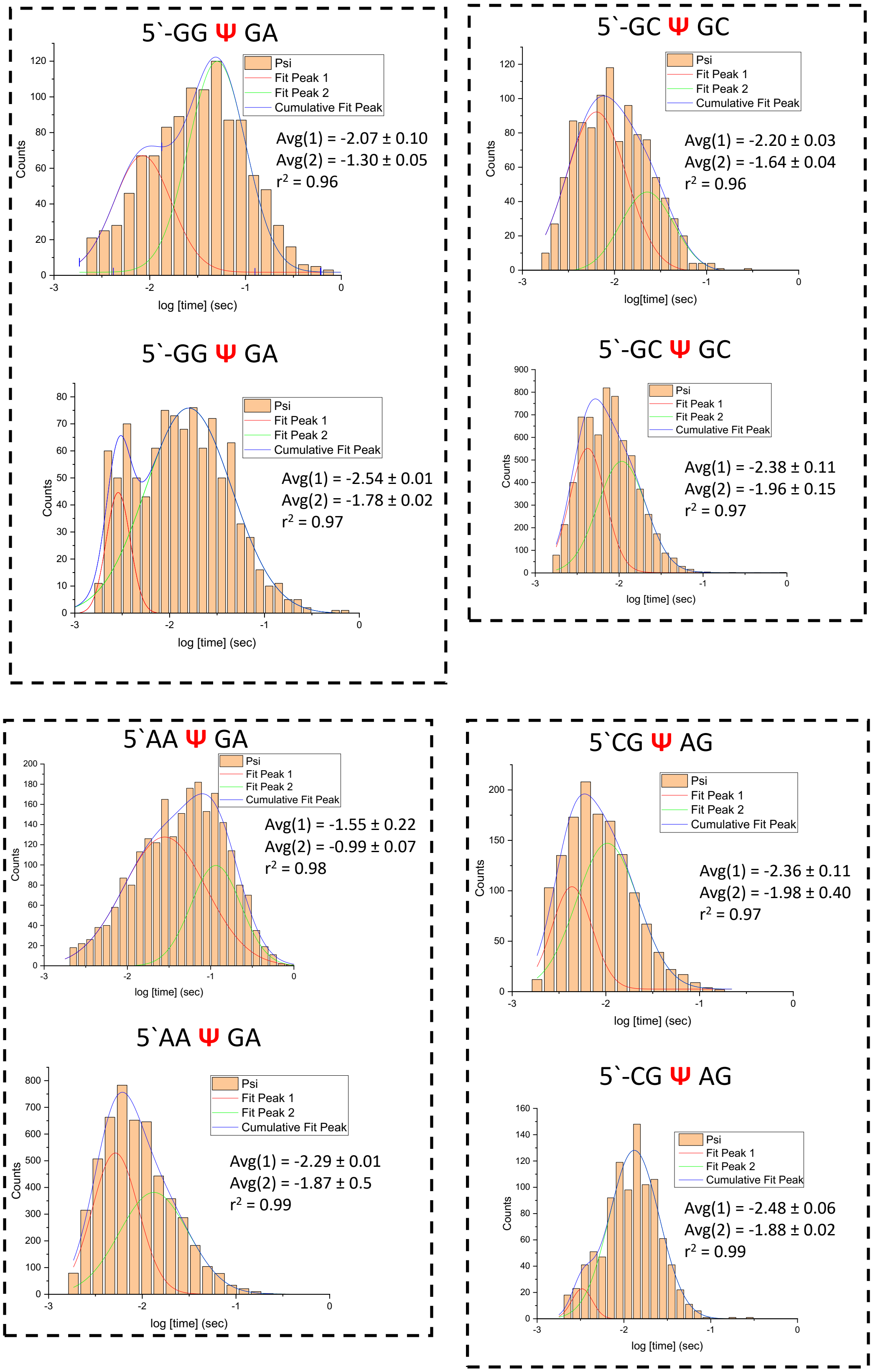

**Figure S11**. Homopolymers can give long-range false positive signals in Nanocompore and Eligos2 analysis.

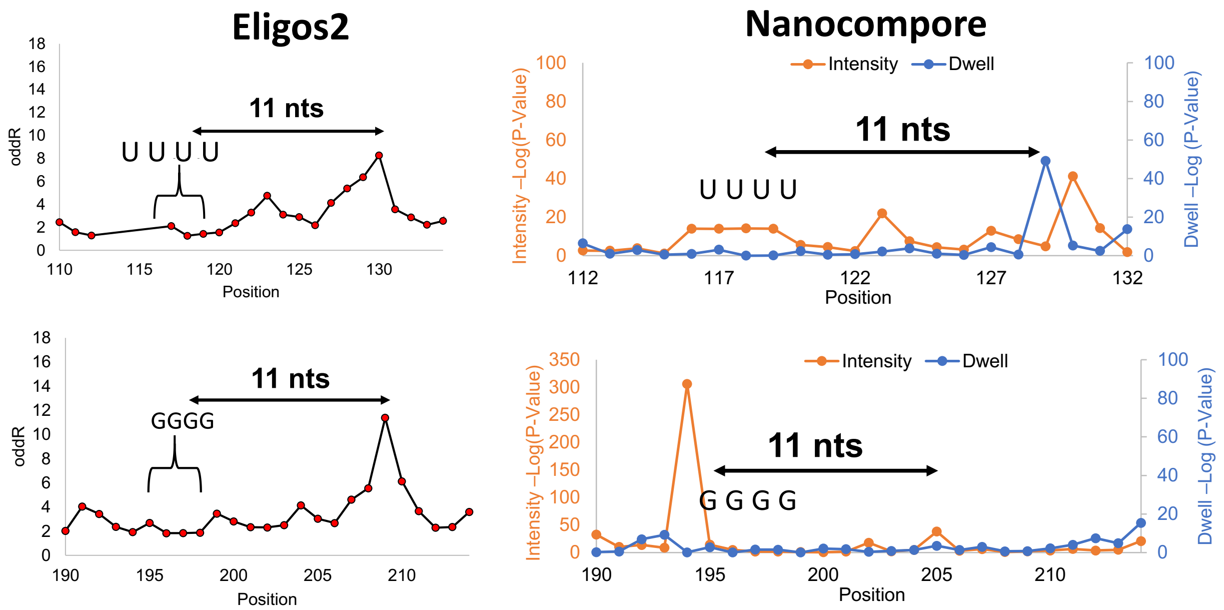


Strand studied with a poly-U track and poly-G track (underlined), studies with poly-C and poly-A were conducted with strand 2. The poly-U and poly-G tracks gave the most pronounced signals that masqueraded as modifications.

5`GAGUAUAGGAUUAGAUAGAUGGCGGAGUUGAAGUAUAGUAGAUUAGAGUCAGAGAAG
AUGAGAUUGAGGUCGGUUAGAAGUUGAUGUAUAGAUGAUG5AGUUAGAUGGAUAGUAAU
UUUAGUAGAGAUUGAAGGUCAAGUAGAUAUGUUAGUAGACCGGUGAUGAGGUGAUAUU
GUCGUGGAUAUUAGAUAUAUGGGGAGAUGAUAGUAGAGGAUUGAAAACAAAAAAAAAAA
A

**Figure S11**. Data Analysis for TRS-S from SARS-CoV-2.

In each block of data, the top plot is for the Nanocompore analysis and the bottom plot is for the Eligos2 analysis to compare the cellular SARS-CoV-2 RNA to the IVT RNA from publicly available data.[1,2] In the Nanocompore plots, the statistically significant differences in the current intensity are shown in orange and the dwell times are shown in blue. In the Eligos2 plots, the significant nucleotide is shown in the red data point. Each block of data is for a single site found by the Nanocompore analysis, long-range dwell impact (black arrow with distance marked), and Eligos2 analysis.

#### U22322 in TRS-S

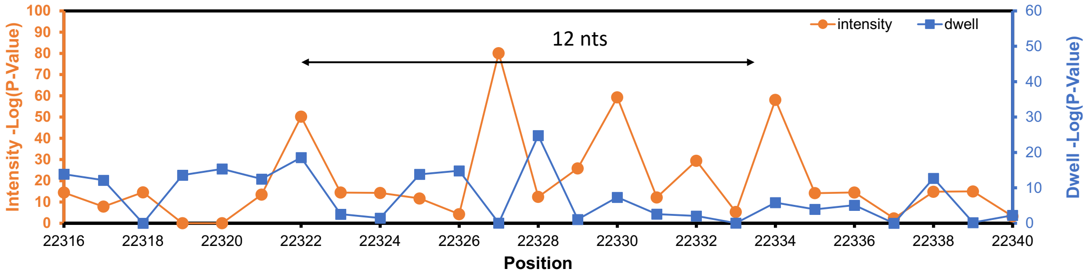


G G U **G A U** U **C U U C** U U C A G G U U G G A C A G

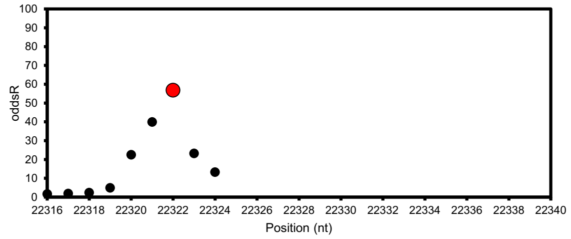


#### U23317 for TRS-S

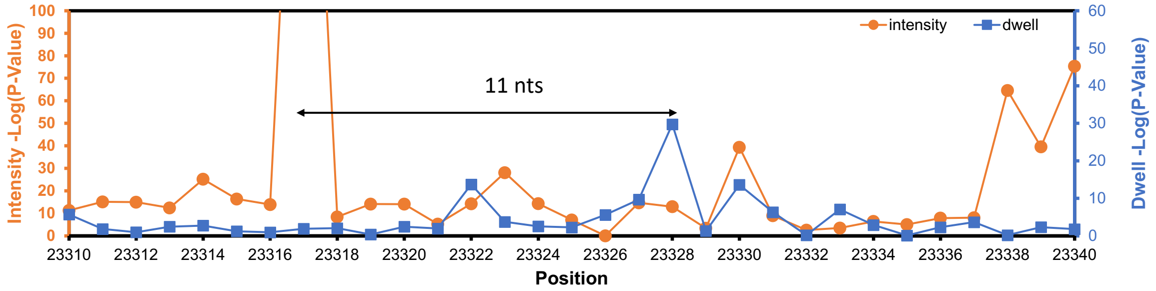


A G A U U C U U G A C A U U A C A C C A U G U U C U U U U G C

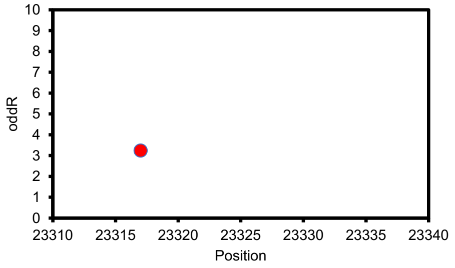

### A24420 in TRS-S

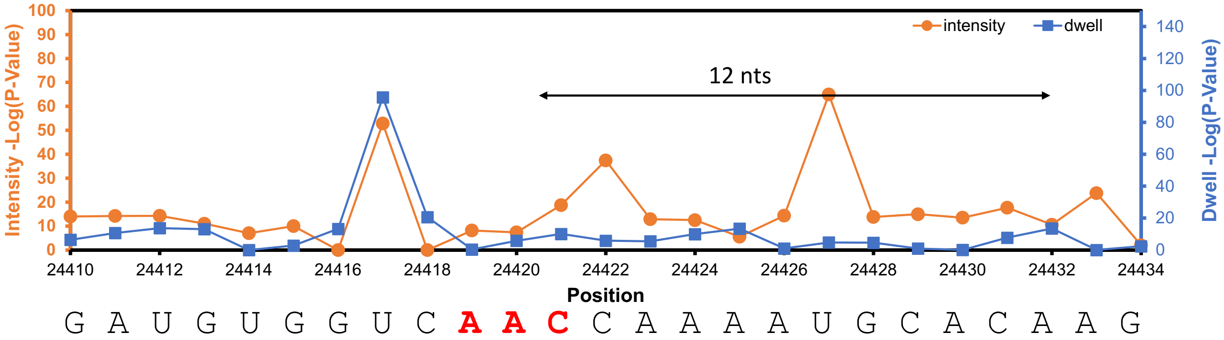


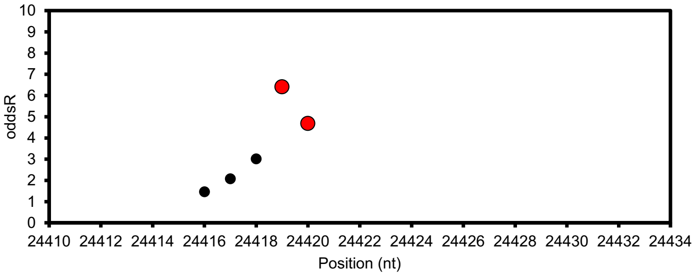


### U27164 in TRS-S

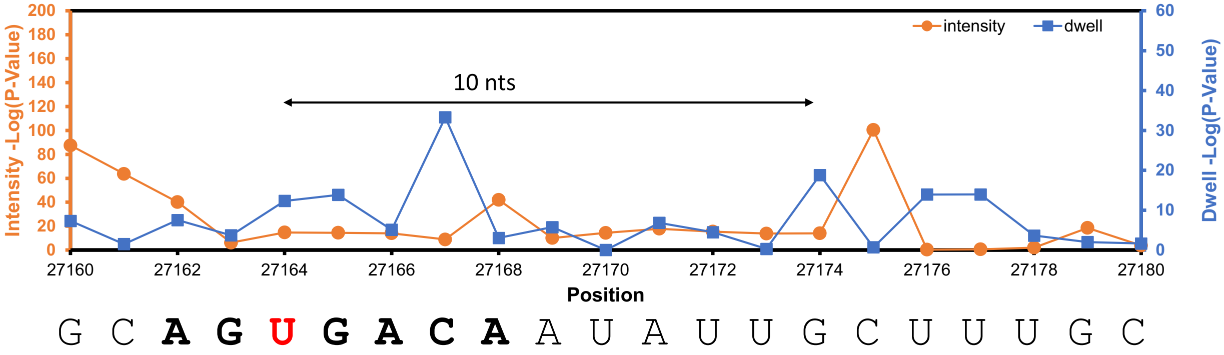


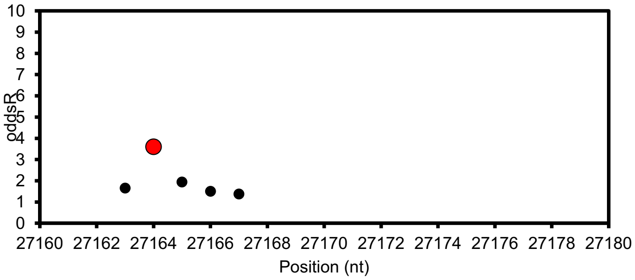

### U28417 in TRS-S

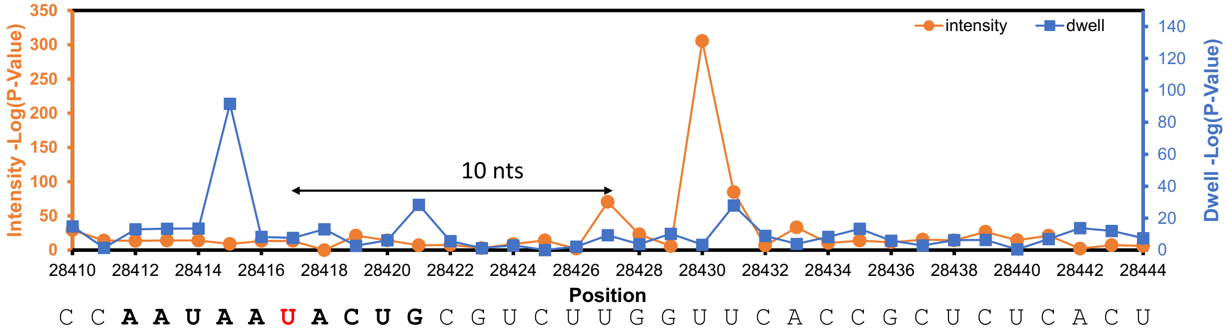


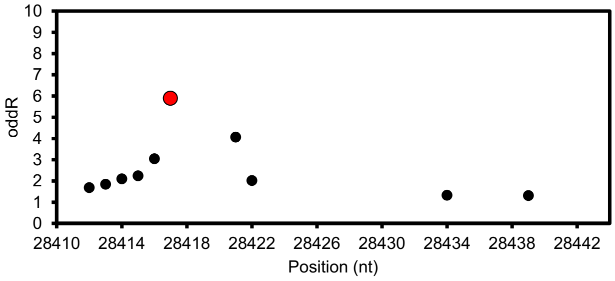


### U28759 in TRS-S

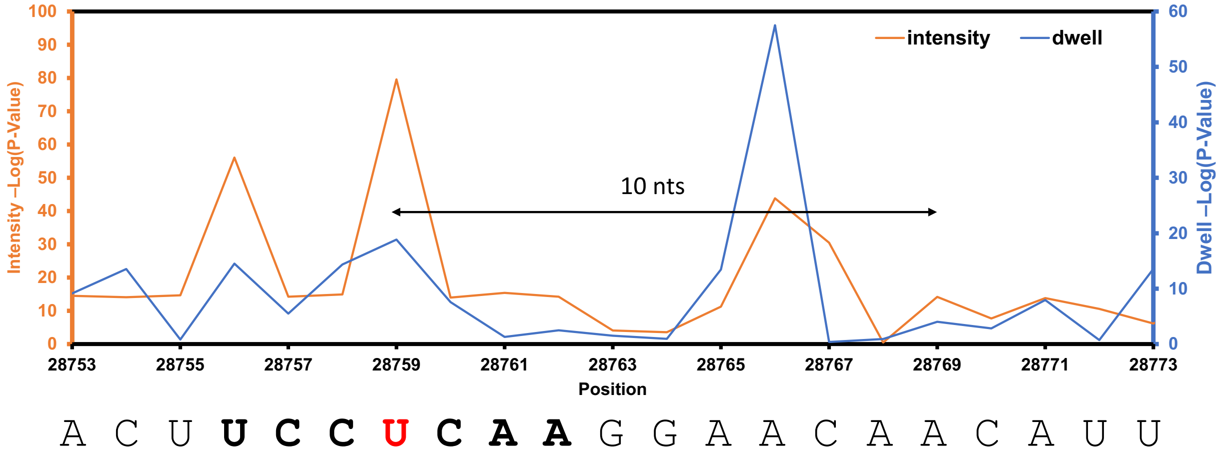


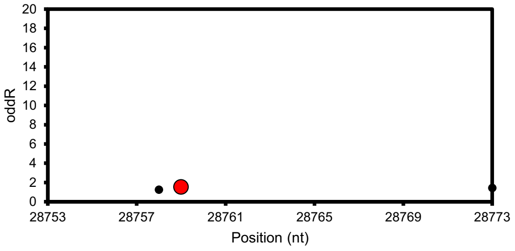

### U28927 for TRS-S

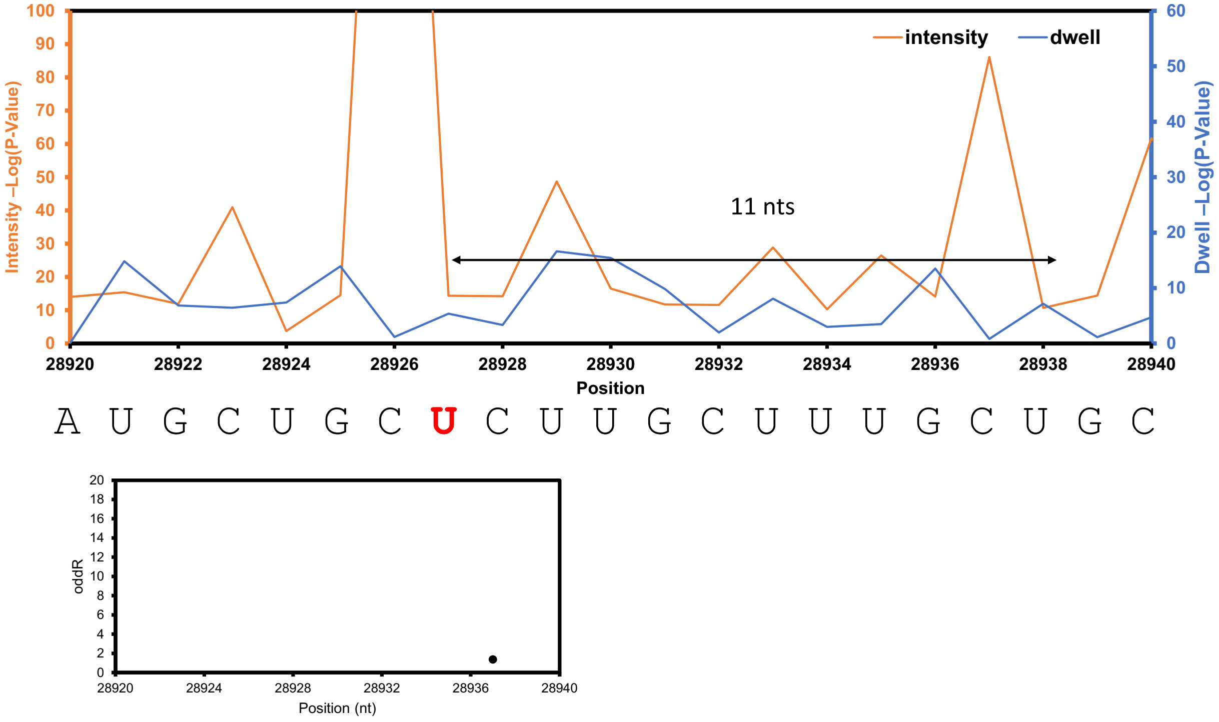


### U29418 for TRS-S

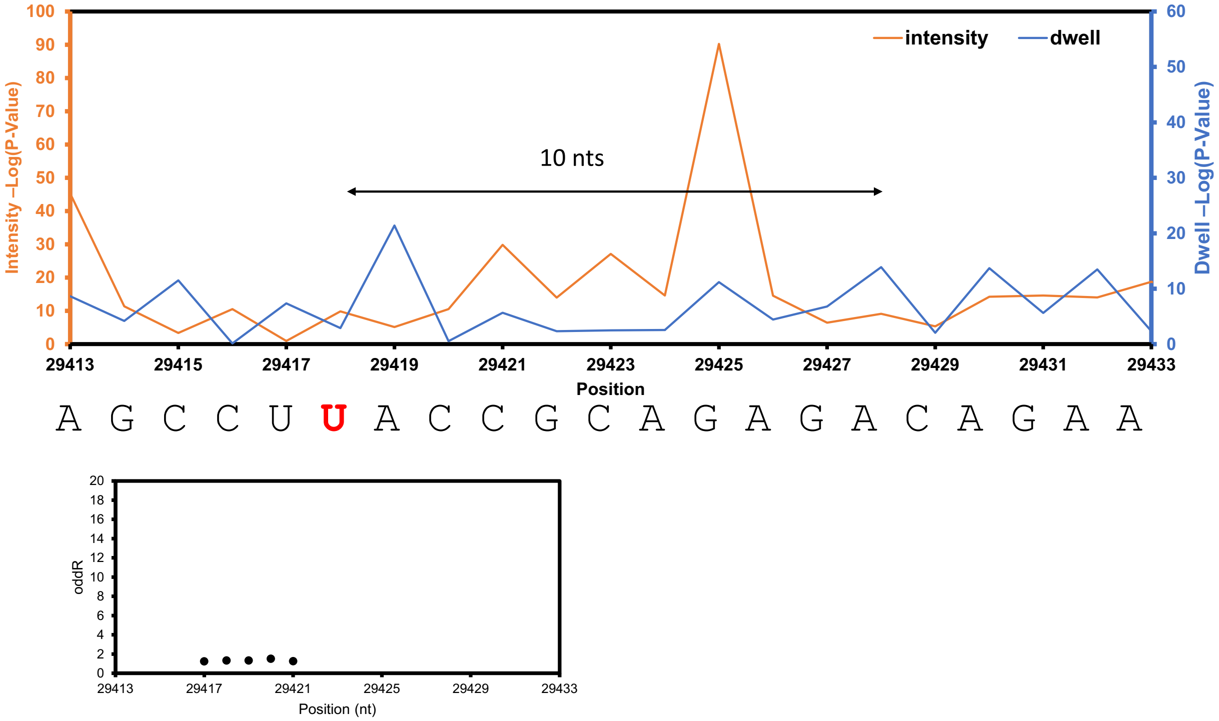

**Figure S12.** Data analysis for TRS-3a from SARS-CoV-2.

In each block of data, the top plot is for the Nanocompore analysis and the bottom plot is for the Eligos2 analysis to compare the cellular SARS-CoV-2 RNA to the IVT RNA from publicly available data.[1,2] In the Nanocompore plots, the statistically significant differences in the current intensity are shown in orange and the dwell times are shown in blue. In the Eligos2 plots, the significant nucleotide is shown in the red data point. Each block of data is for a single site found by the Nanocompore analysis, long-range dwell impact (black arrow with distance marked), and Eligos2 analysis.

#### U27164 for TRS-3a

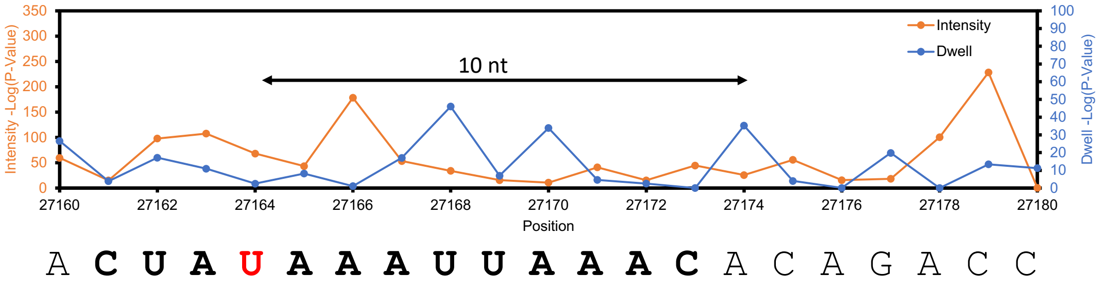


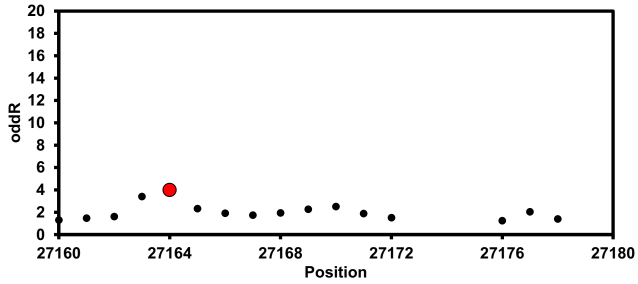


#### A27333-27334 in TRS-3a

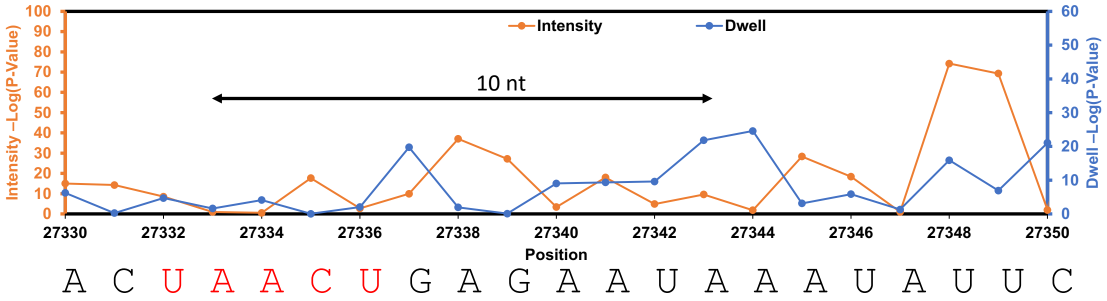


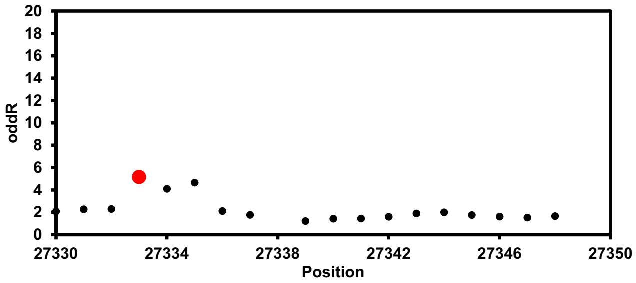

### C27600 in TRS-3a

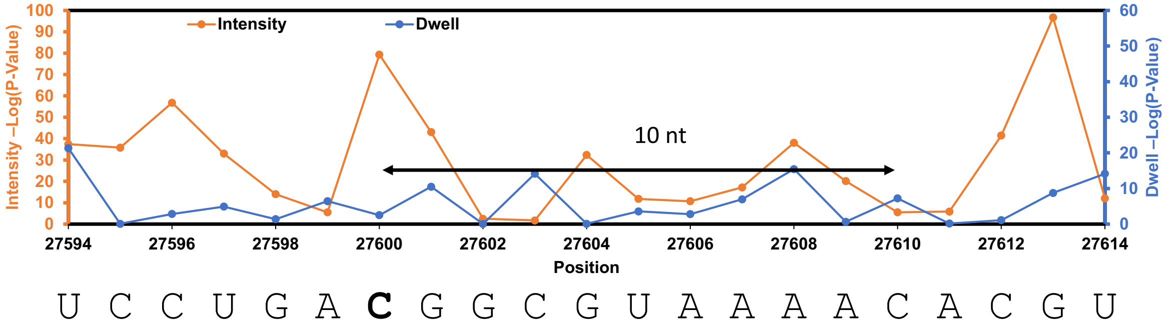


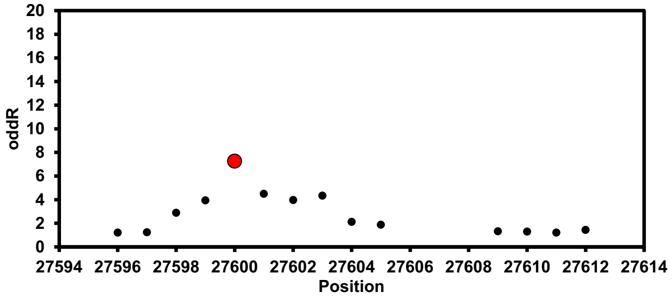


### U28039 and A28047 for TRS-3A

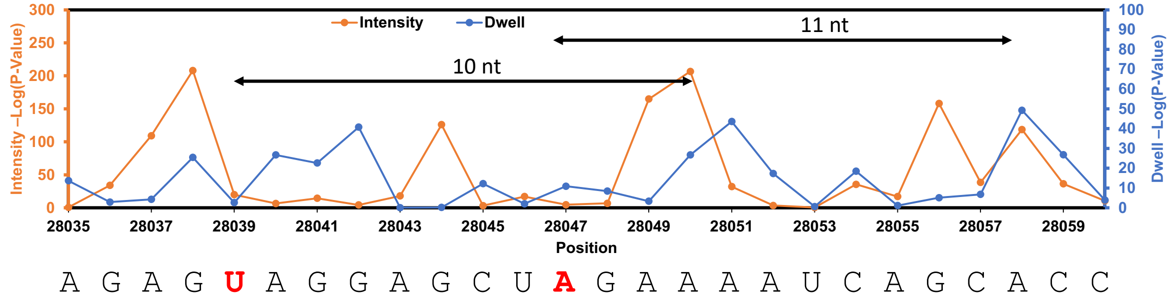


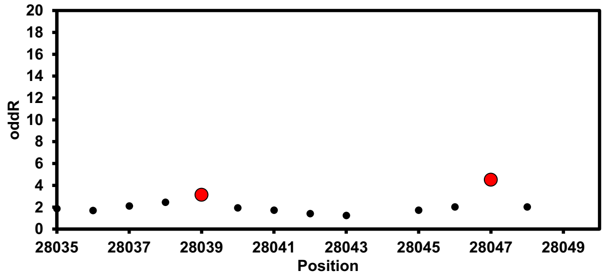

### U28759 in TRS-3a

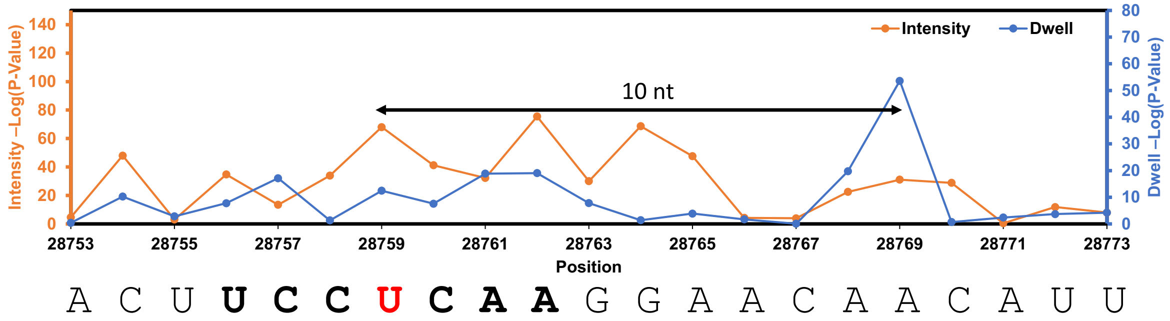


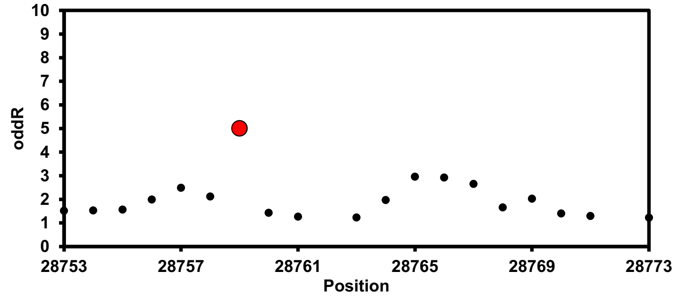


### U28927 in TRS-3a

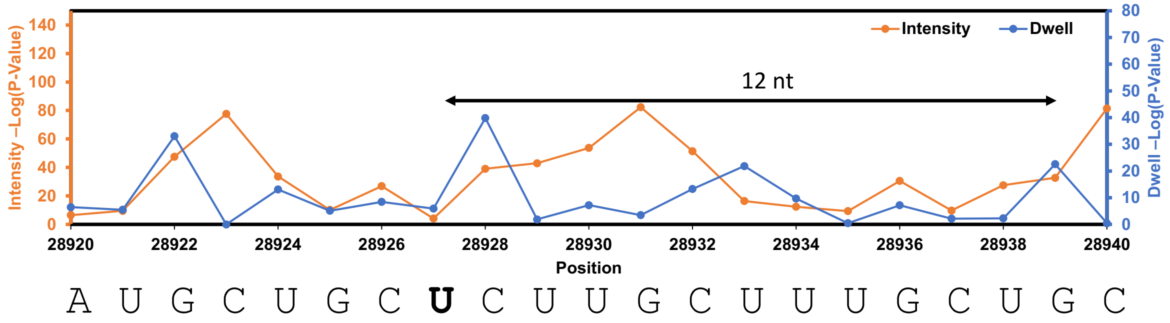


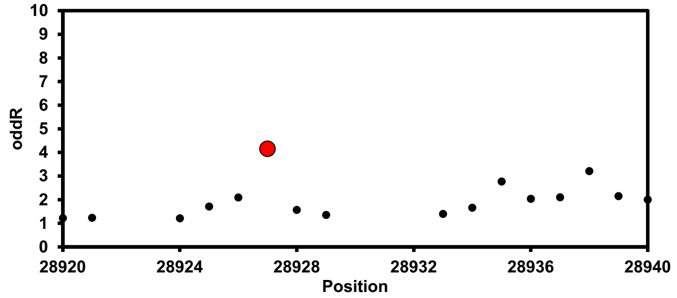

### A28993 in TRS-3a

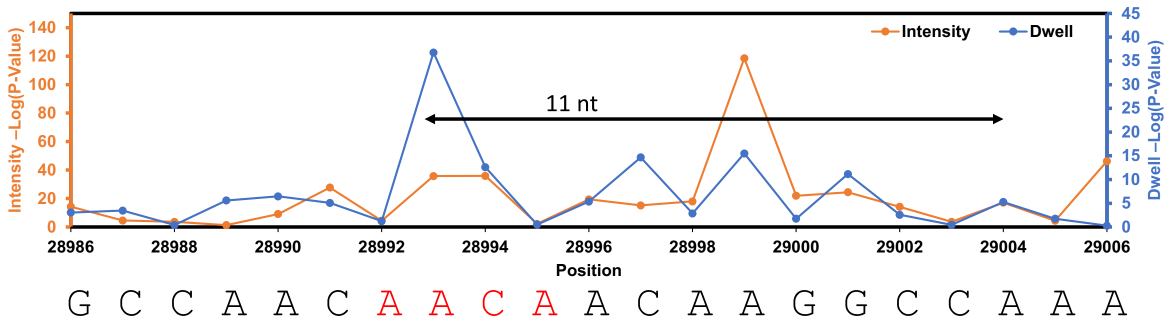


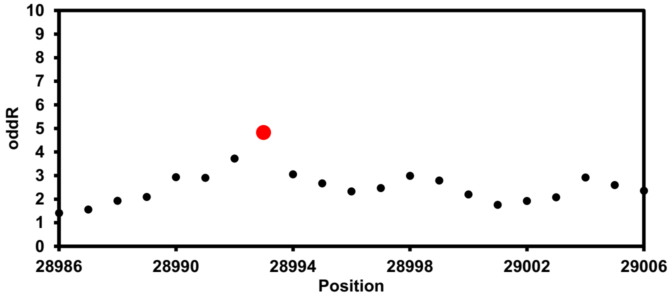


### A29285 in TRS-3a

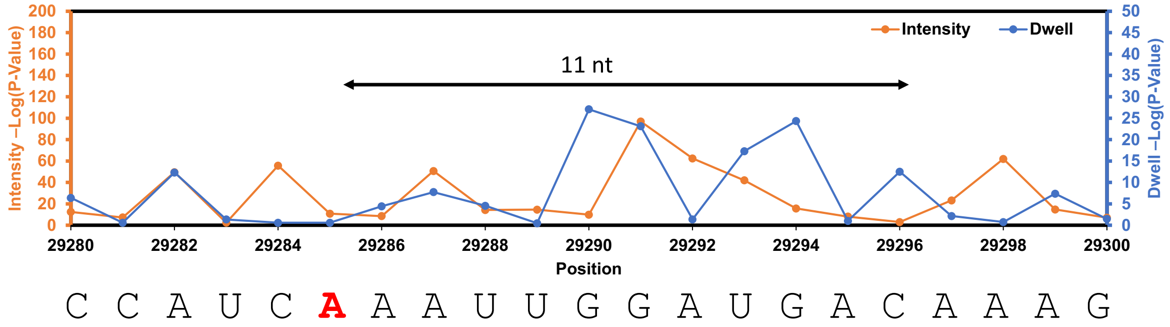


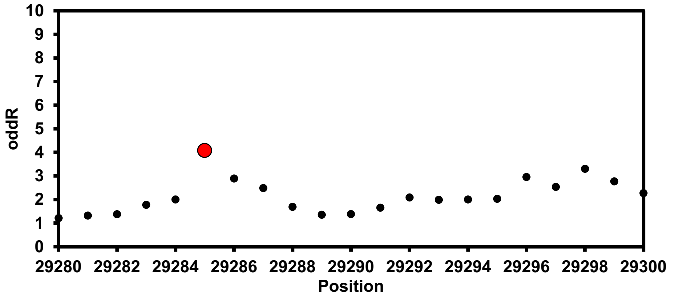

### U29418 in TRS-3a

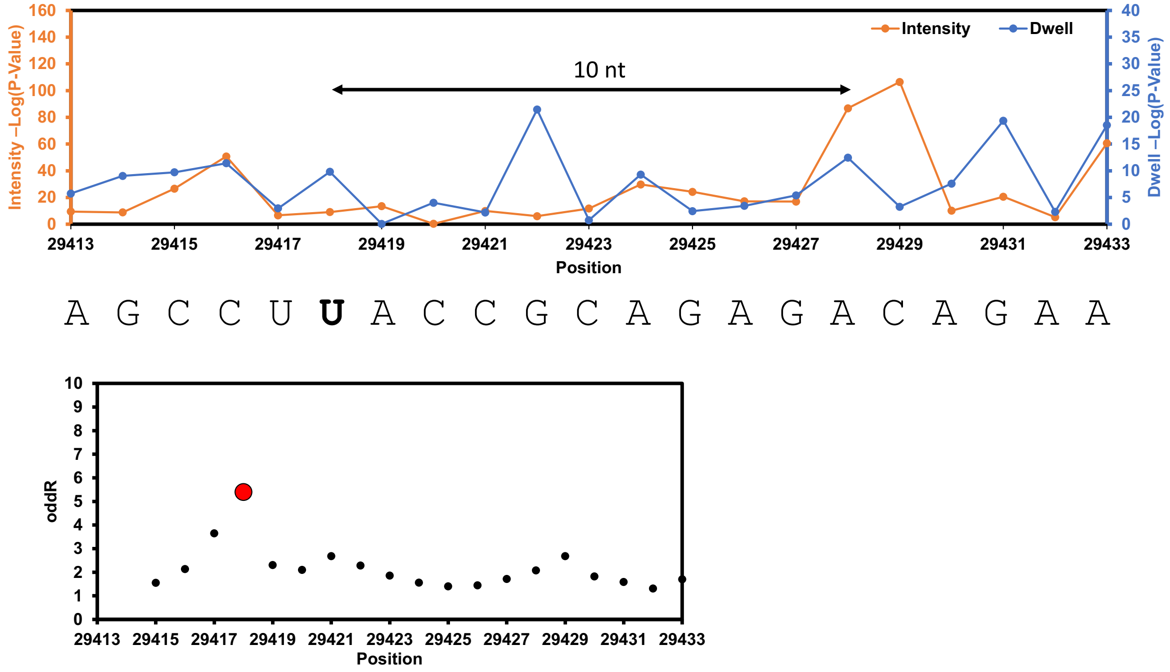

**Figure S13.** Data analysis for TRS-E from SARS-CoV-2.

In each block of data, the top plot is for the Nanocompore analysis and the bottom plot is for the Eligos2 analysis to compare the cellular SARS-CoV-2 RNA to the IVT RNA from publicly available data.[1,2] In the Nanocompore plots, the statistically significant differences in the current intensity are shown in orange and the dwell times are shown in blue. In the Eligos2 plots, the significant nucleotide is shown in the red data point. Each block of data is for a single site found by the Nanocompore analysis, long-range dwell impact (black arrow with distance marked), and Eligos2 analysis.

#### U27164 for TRS-E

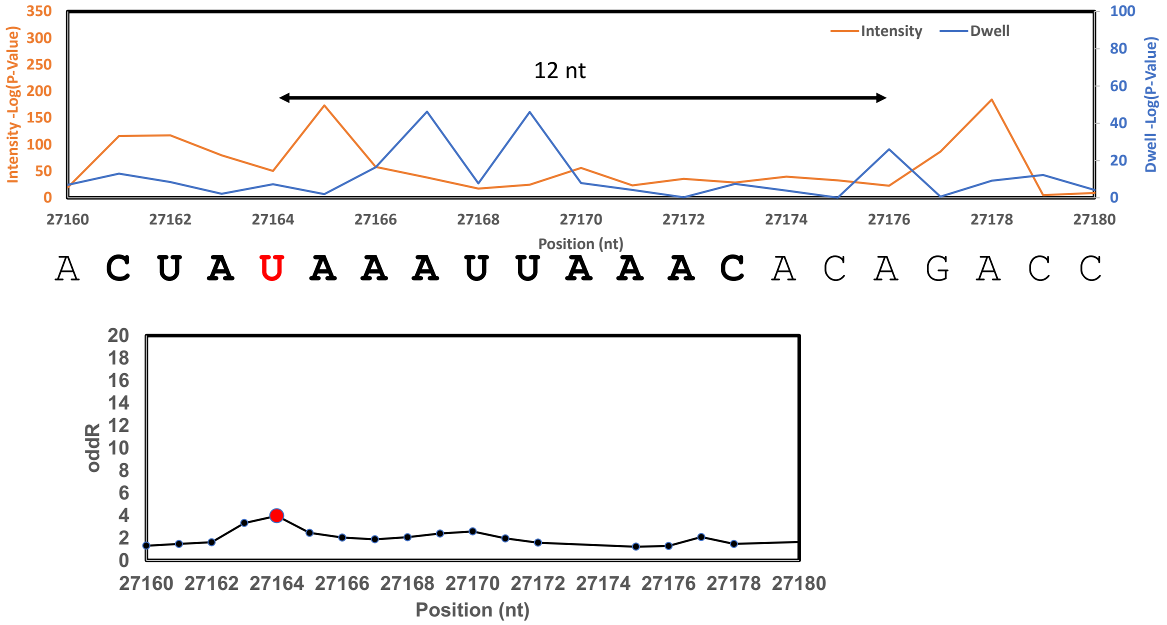


#### U28039 and A28047 for TRS-M

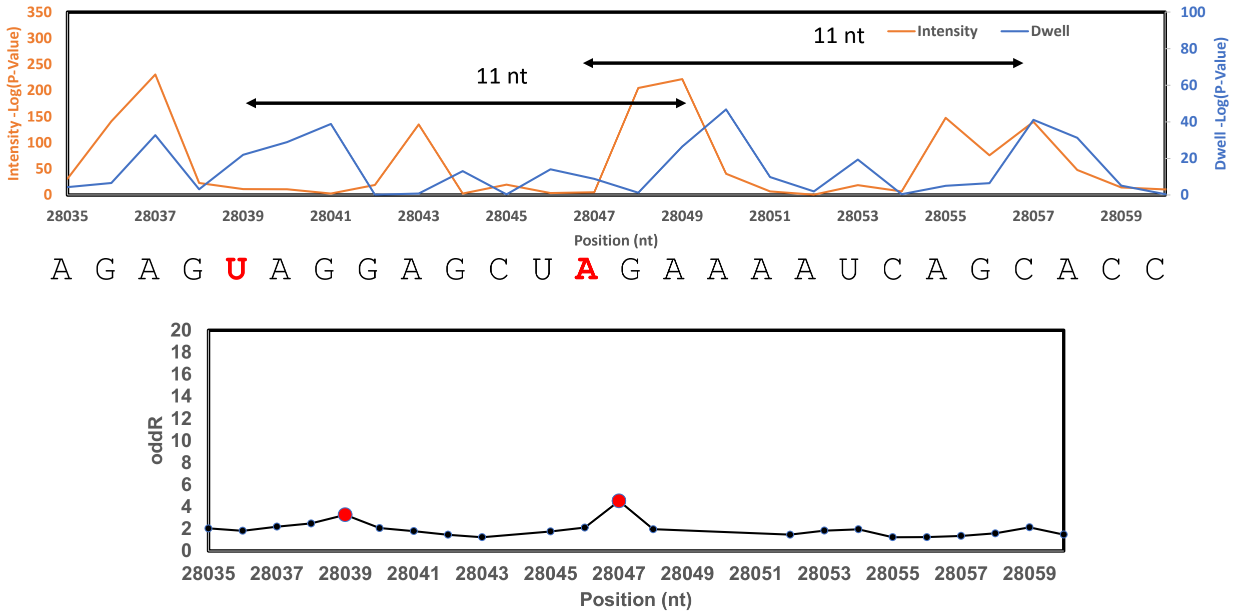

### U28759 for TRS-E

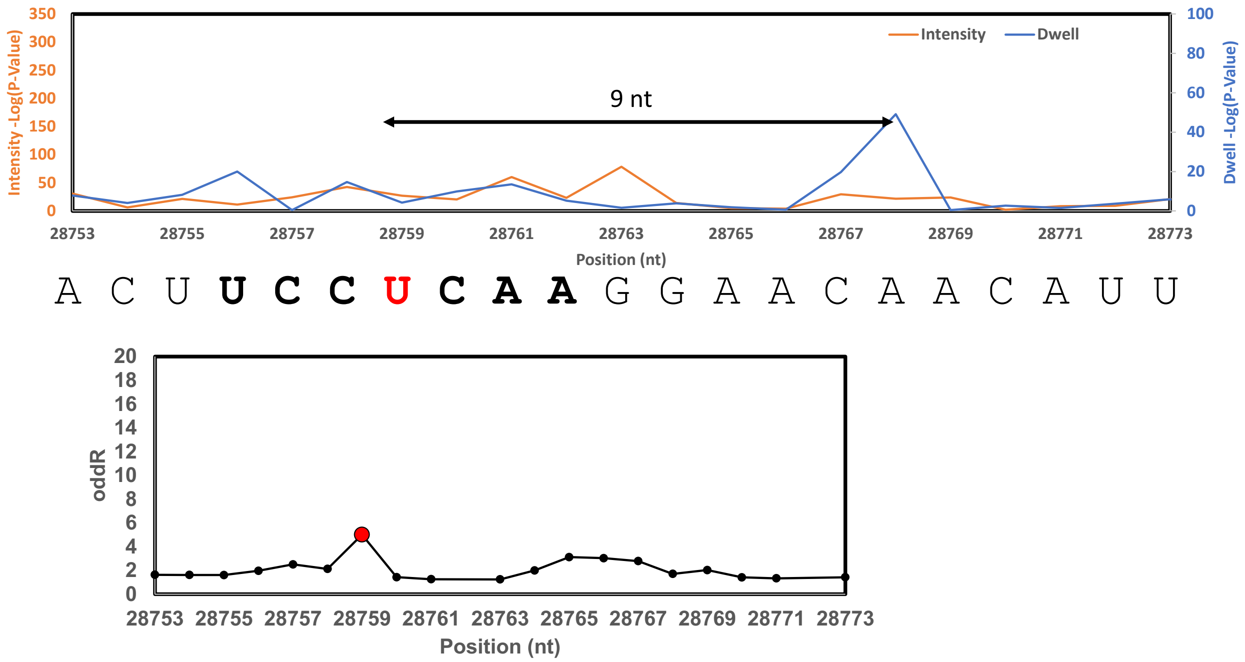


### U28927 for TRS-E

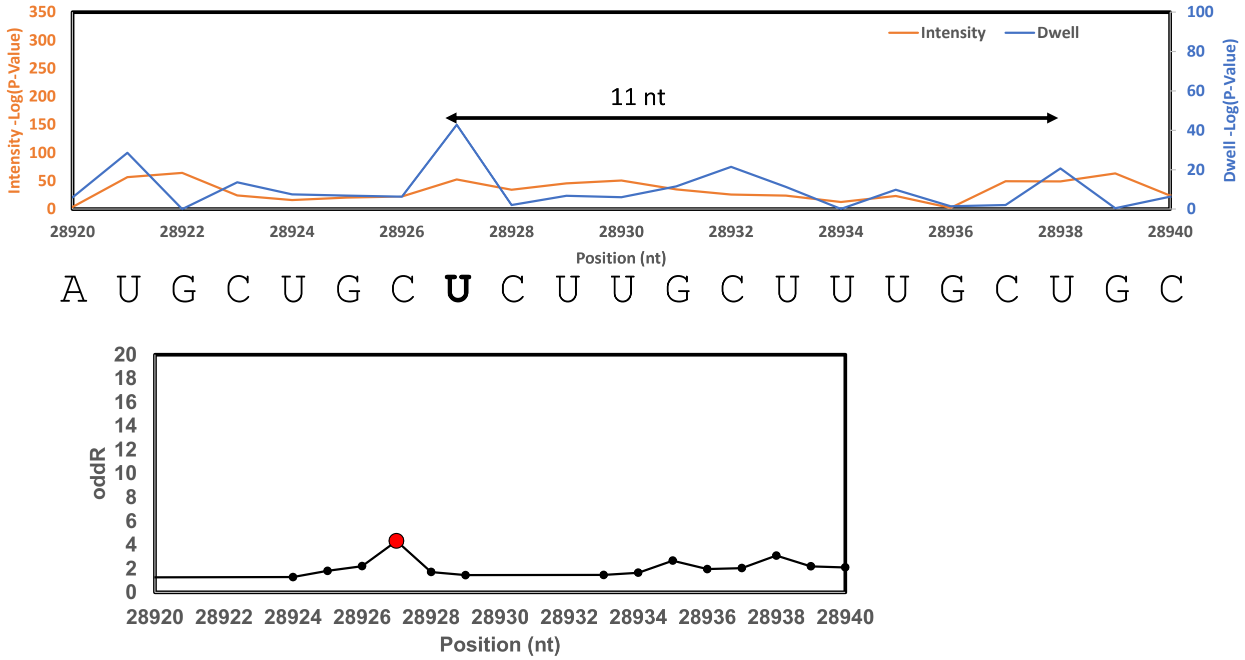

### U29418 for TRS-E

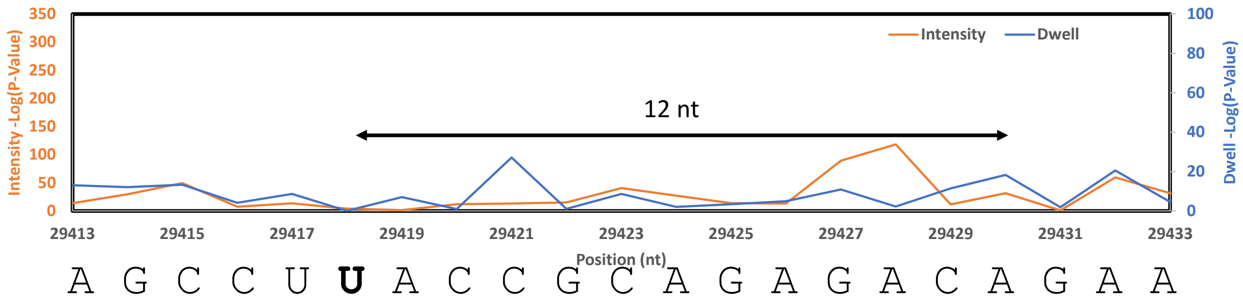


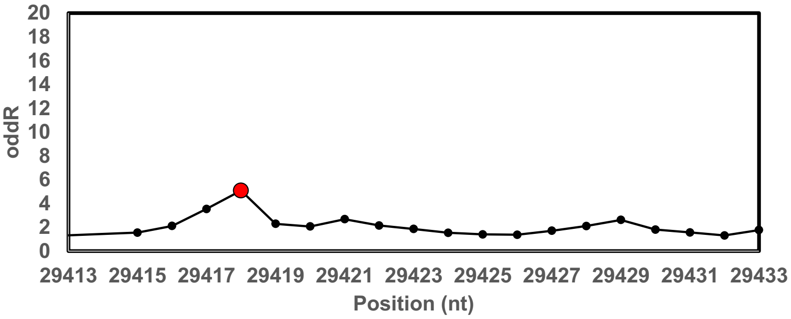

**Figure S14.** Data analysis for TRS-M from SARS-CoV-2.

In each block of data, the top plot is for the Nanocompore analysis and the bottom plot is for the Eligos2 analysis to compare the cellular SARS-CoV-2 RNA to the IVT RNA from publicly available data.[1,2] In the Nanocompore plots, the statistically significant differences in the current intensity are shown in orange and the dwell times are shown in blue. In the Eligos2 plots, the significant nucleotide is shown in the red data point. Each block of data is for a single site found by the Nanocompore analysis, long-range dwell impact (black arrow with distance marked), and Eligos2 analysis.

#### U27164 for TRS-M

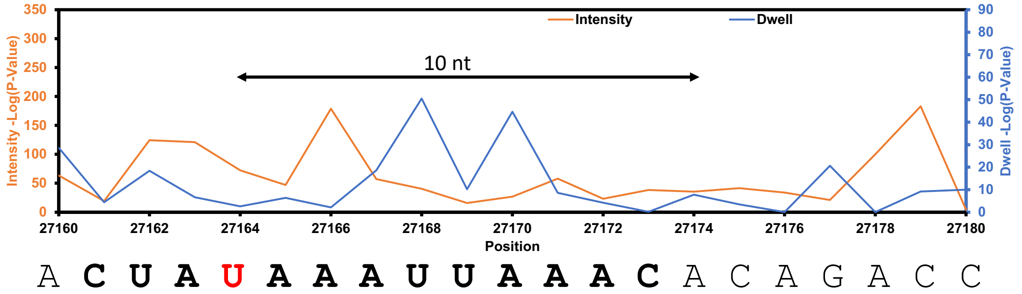


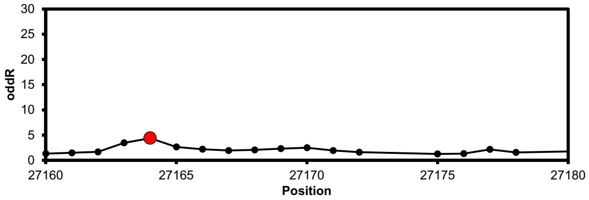


#### U28047 TRS-M

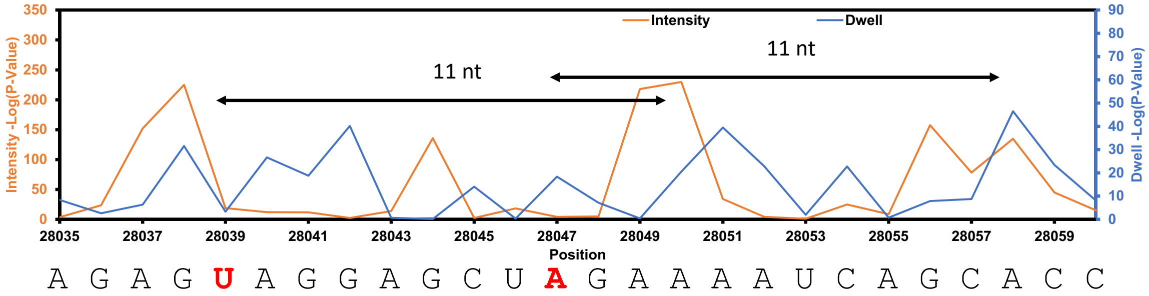


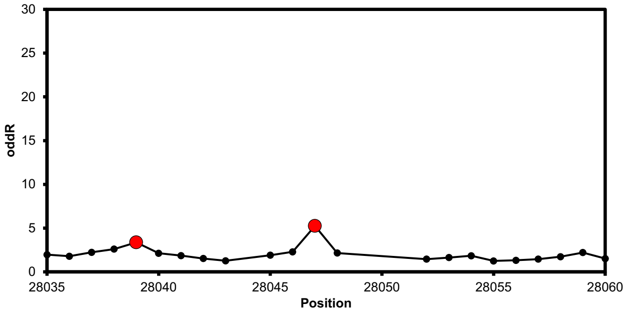

### U28759 for TRS-M

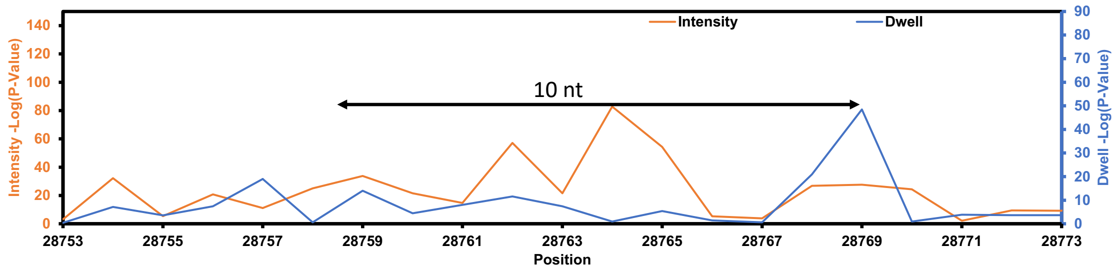


A C U **U C C** U **C A A** G G A A C A A C A U U

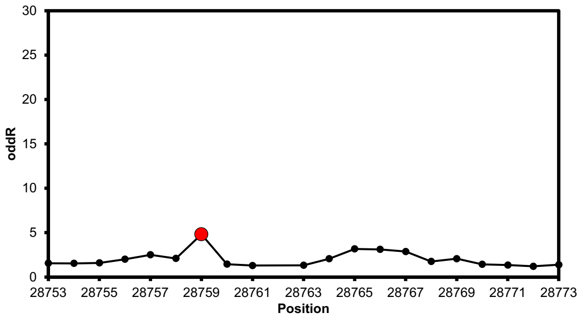


### U28927 for TRS-M

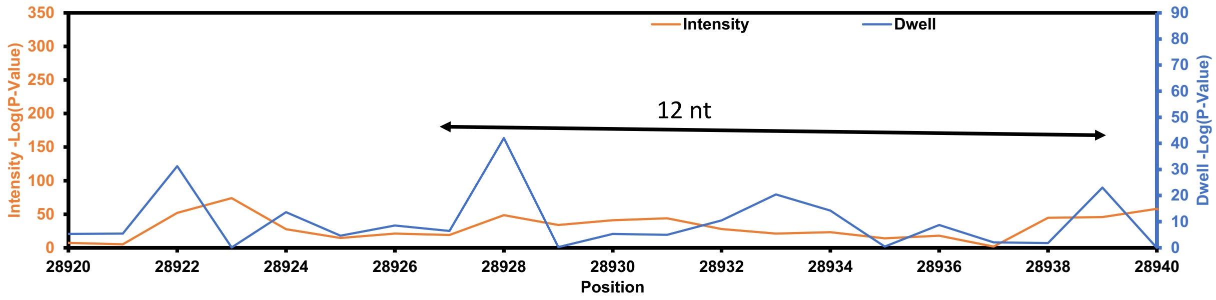


A U G C U G C **U** C U U G C U U U G C U G C

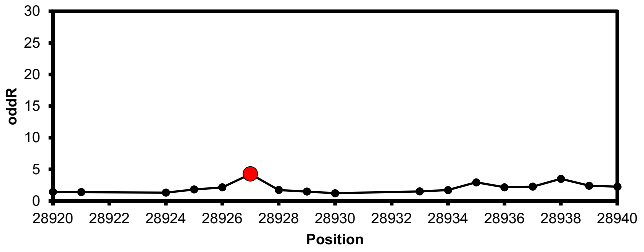

### U29418 for TRS-M

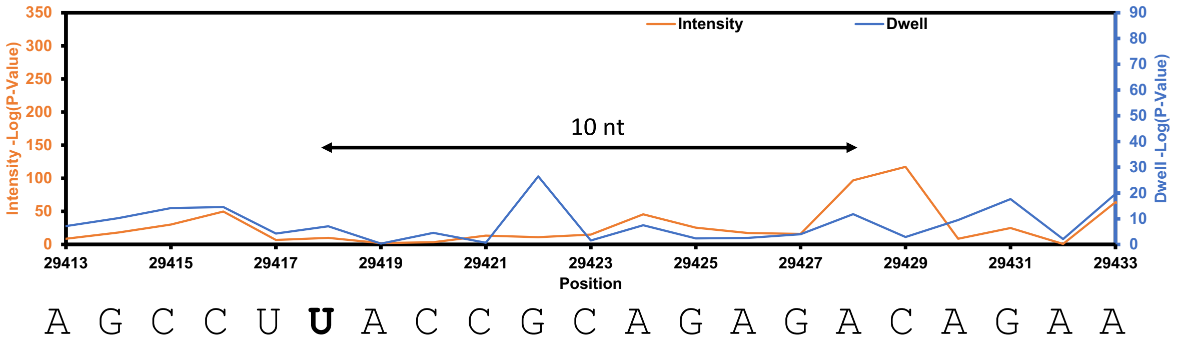


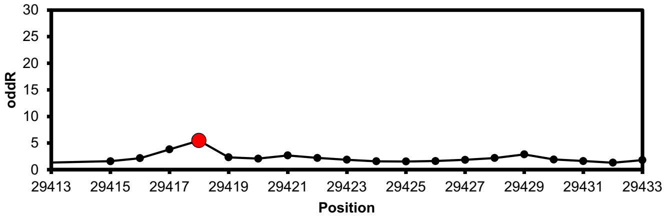

**Figure S15.** Data analysis for TRS-6 from SARS-CoV-2.

In each block of data, the top plot is for the Nanocompore analysis and the bottom plot is for the Eligos2 analysis to compare the cellular SARS-CoV-2 RNA to the IVT RNA from publicly available data.[1,2] In the Nanocompore plots, the statistically significant differences in the current intensity are shown in orange and the dwell times are shown in blue. In the Eligos2 plots, the significant nucleotide is shown in the red data point. Each block of data is for a single site found by the Nanocompore analysis, long-range dwell impact (black arrow with distance marked), and Eligos2 analysis.

#### U27164 for TRS-6

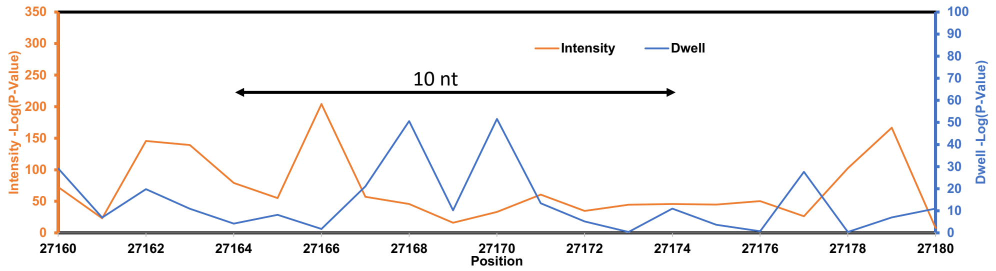


A C U A U A A A U U A A A C A C A G A C C

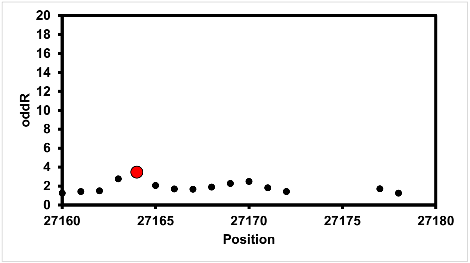


#### U28039 and A28047 for TRS-6

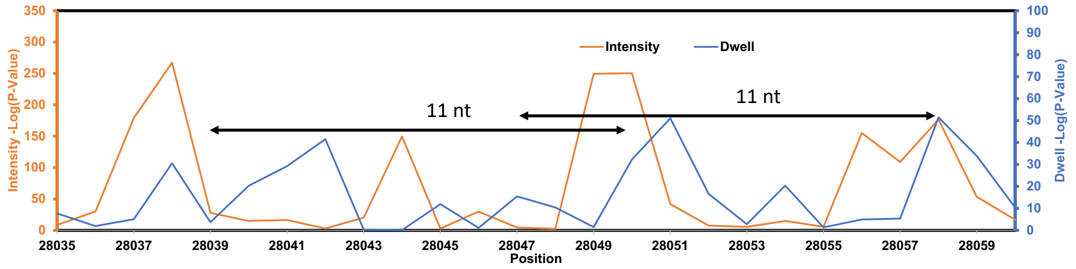


A G A G U A G G A G C U A G A A A A U C A G C A C C

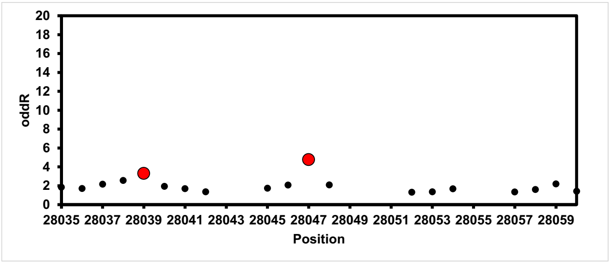

### U28759 for TRS-6

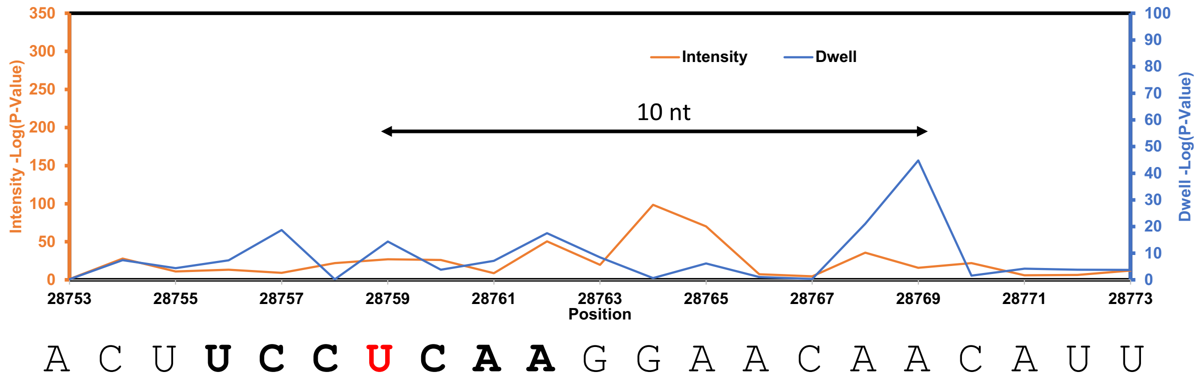


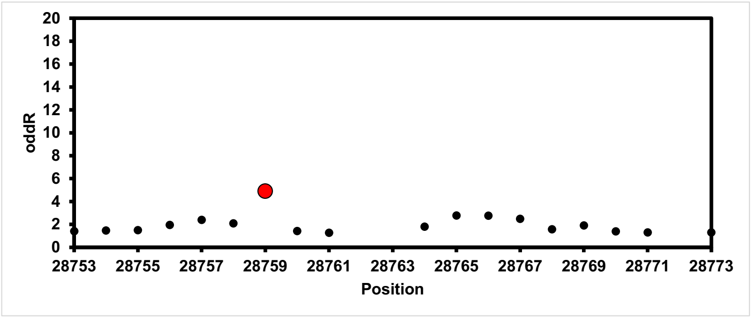


### U28927 for TRS-6

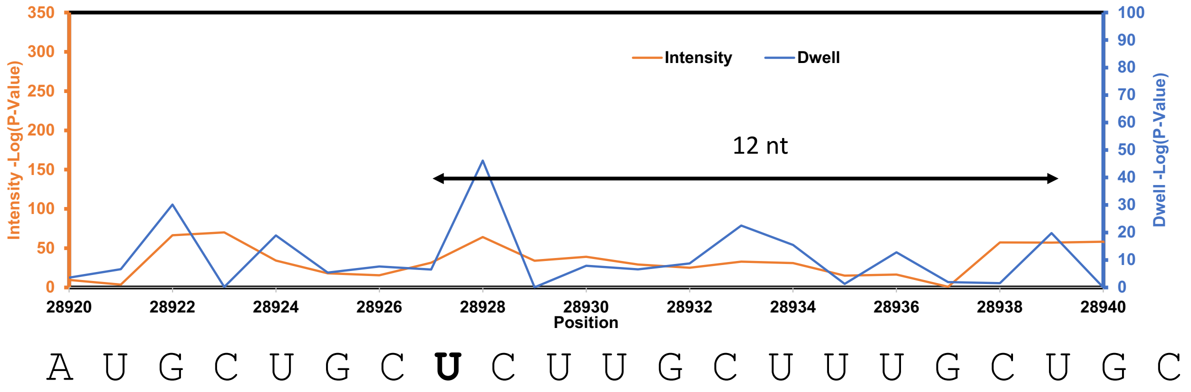


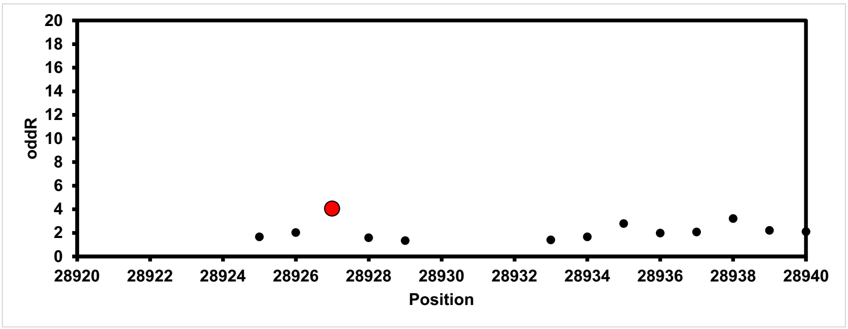

### U29418 for TRS-6

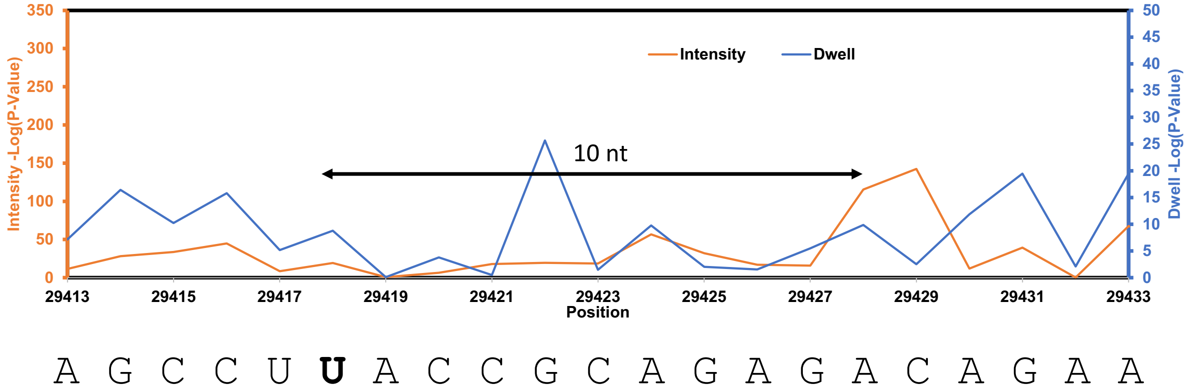


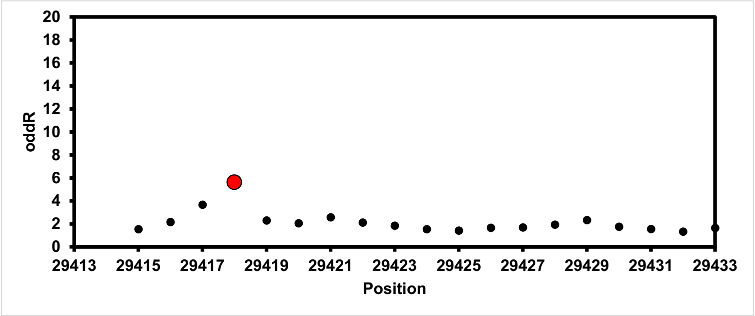

**Figure S16.** Data analysis for TRS-7a from SARS-CoV-2.

In each block of data, the top plot is for the Nanocompore analysis and the bottom plot is for the Eligos2 analysis to compare the cellular SARS-CoV-2 RNA to the IVT RNA from publicly available data.[1,2] In the Nanocompore plots, the statistically significant differences in the current intensity are shown in orange and the dwell times are shown in blue. In the Eligos2 plots, the significant nucleotide is shown in the red data point. Each block of data is for a single site found by the Nanocompore analysis, long-range dwell impact (black arrow with distance marked), and Eligos2 analysis.

#### U28039 and A28047 for TRS-7a

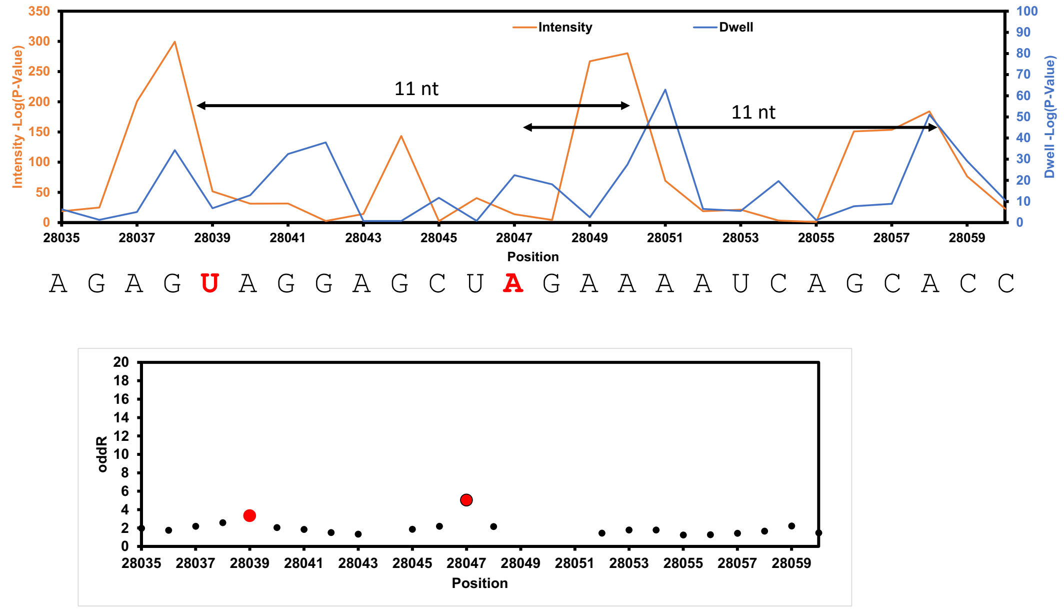


#### U28759 for TRS-7a

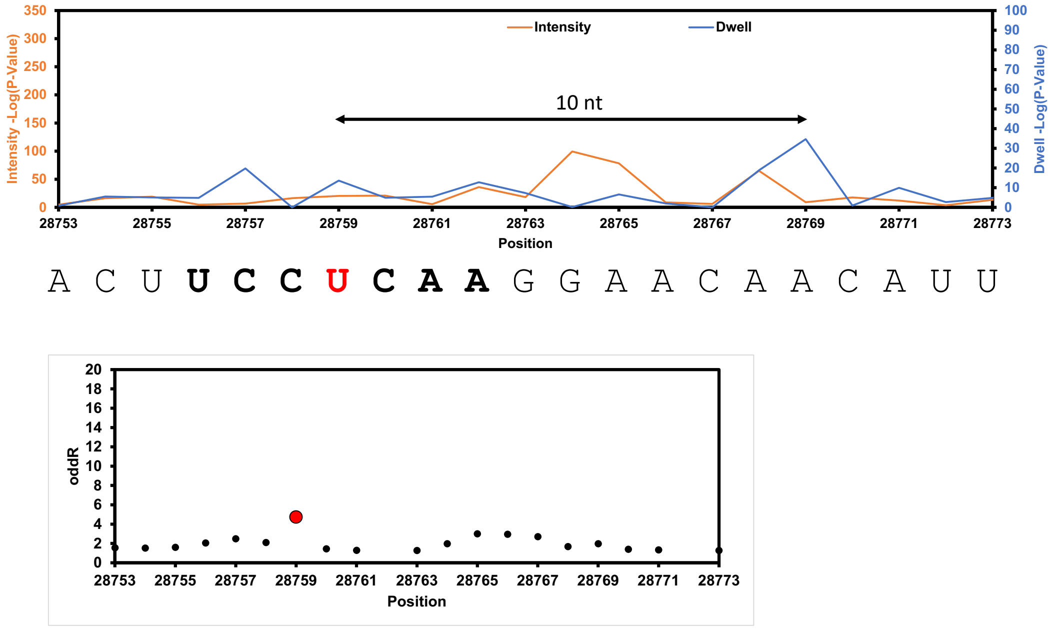

### U28927 for TRS-7a

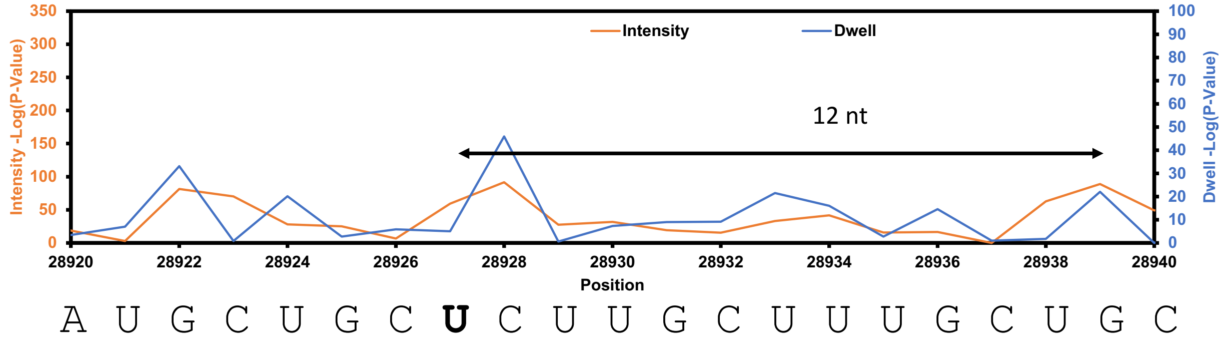


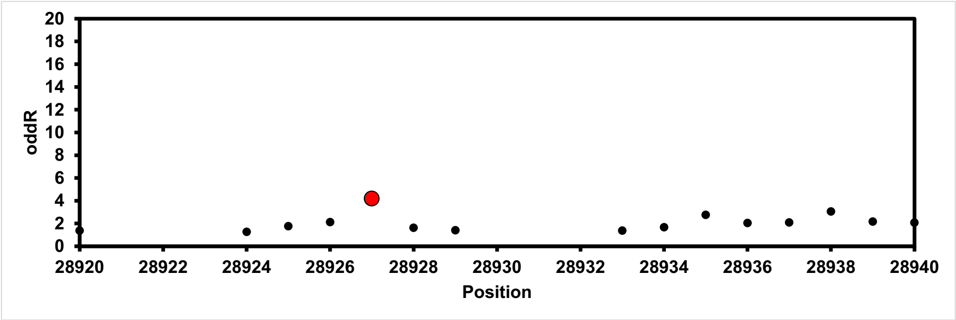


### U29418 for TRS-7a

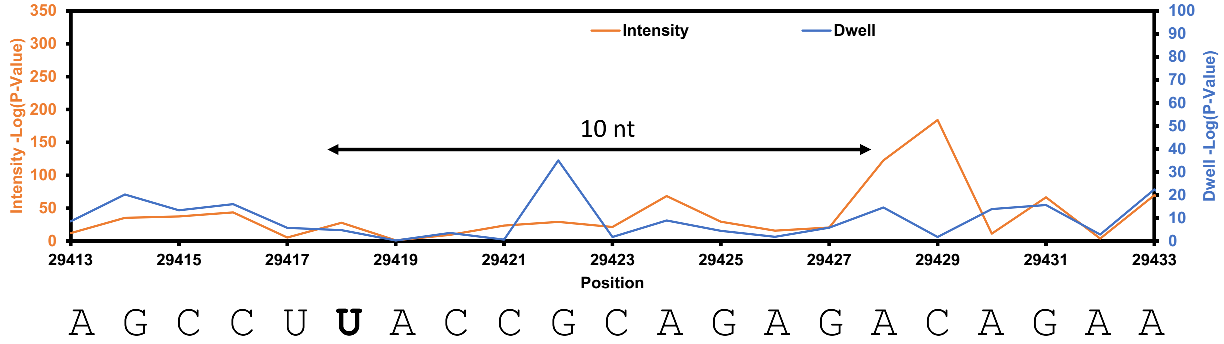


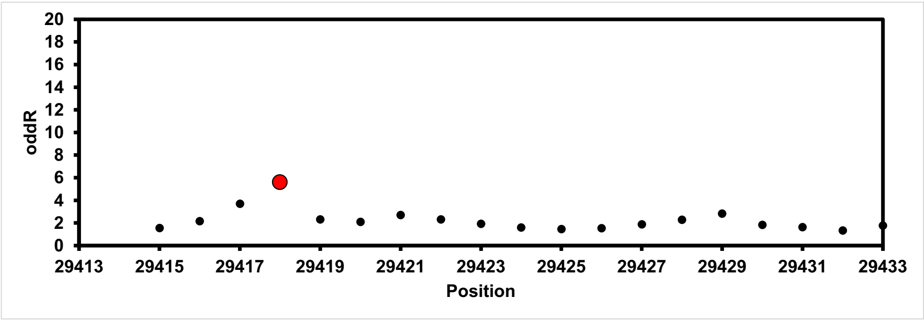

**Figure S17.** Data analysis for TRS-7b from SARS-CoV-2.

In each block of data, the top plot is for the Nanocompore analysis and the bottom plot is for the Eligos2 analysis to compare the cellular SARS-CoV-2 RNA to the IVT RNA from publicly available data.[1,2] In the Nanocompore plots, the statistically significant differences in the current intensity are shown in orange and the dwell times are shown in blue. In the Eligos2 plots, the significant nucleotide is shown in the red data point. Each block of data is for a single site found by the Nanocompore analysis, long-range dwell impact (black arrow with distance marked), and Eligos2 analysis.

#### U28039 and A28047 in TRS-7b

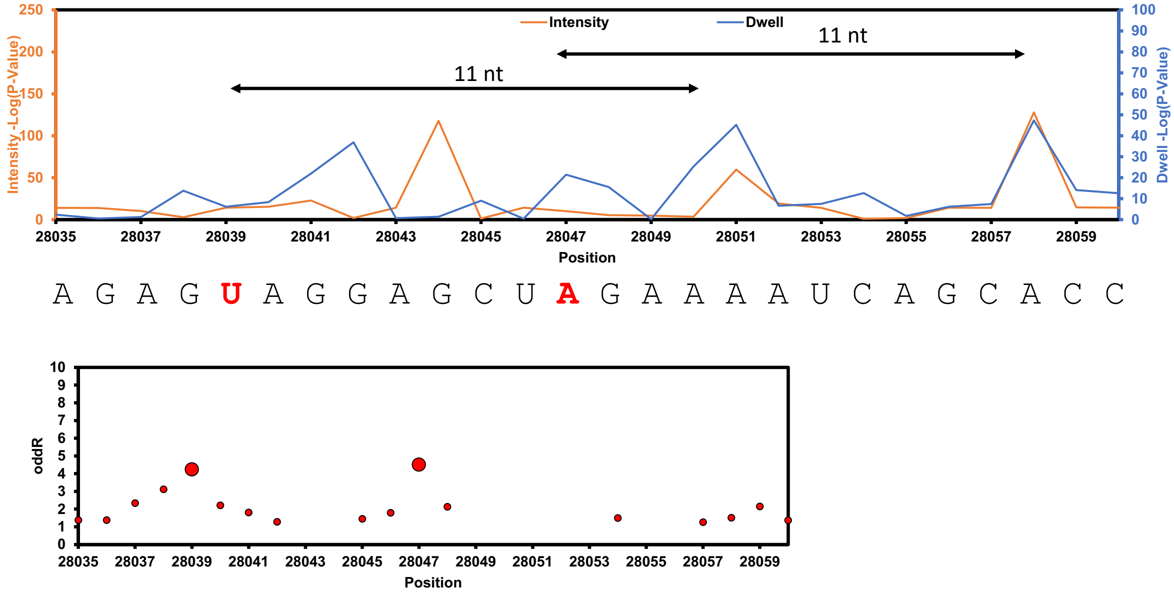


#### U28759 for TRS-7b

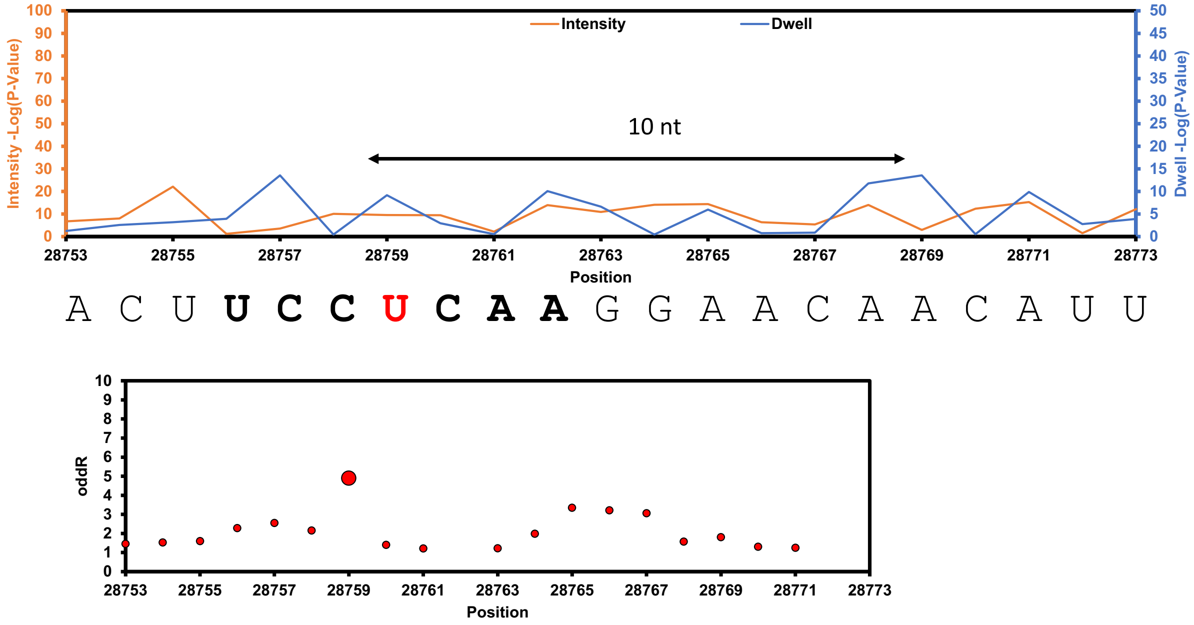

### U28927 for TRS-7b

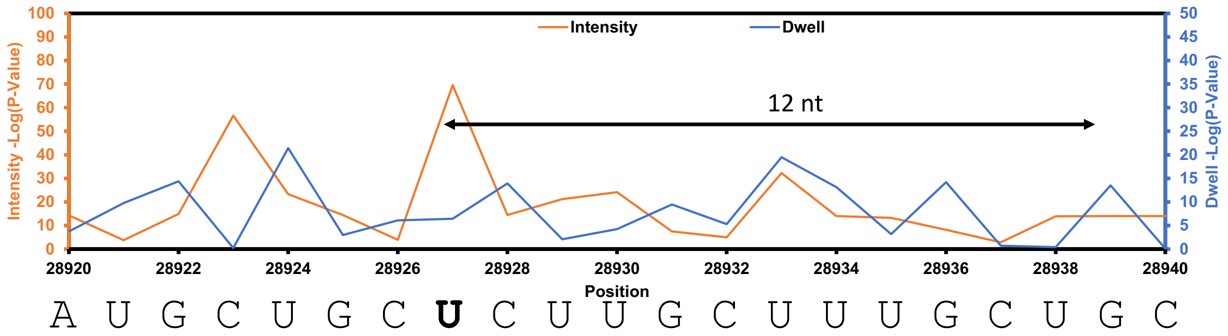


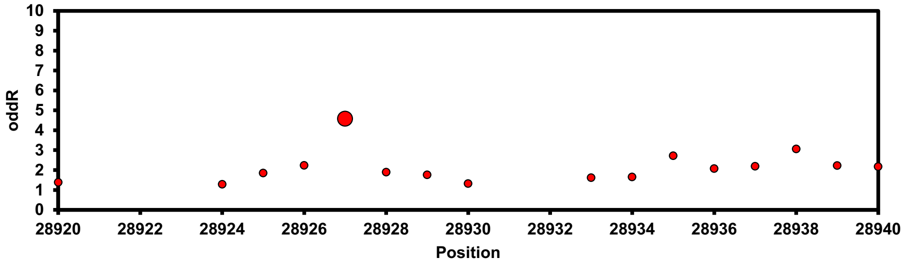


### U29418 for TRS-7b

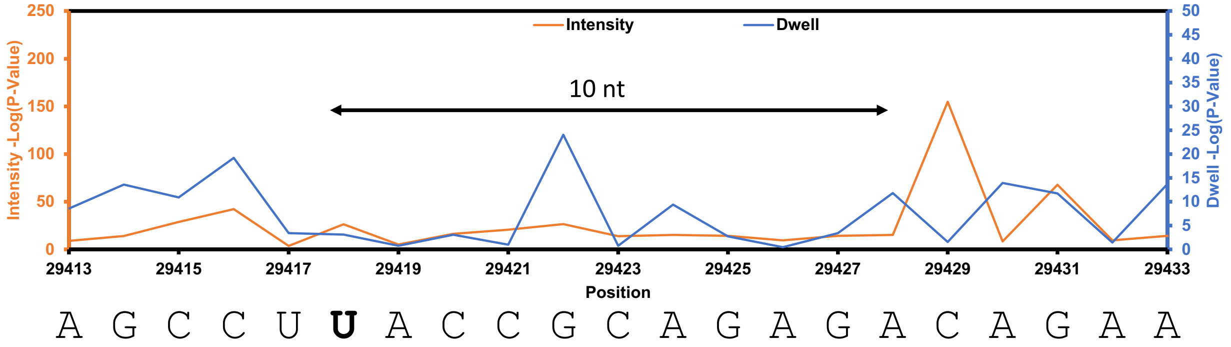


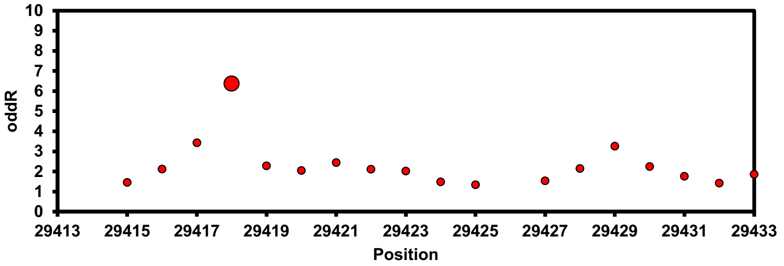

**Figure S18**. Data analysis for TRS-8 from SARS-CoV-2.

In each block of data, the top plot is for the Nanocompore analysis and the bottom plot is for the Eligos2 analysis to compare the cellular SARS-CoV-2 RNA to the IVT RNA from publicly available data.[1,2] In the Nanocompore plots, the statistically significant differences in the current intensity are shown in orange and the dwell times are shown in blue. In the Eligos2 plots, the significant nucleotide is shown in the red data point. Each block of data is for a single site found by the Nanocompore analysis, long-range dwell impact (black arrow with distance marked), and Eligos2 analysis.

#### U28039 and A28047 for TRS-8

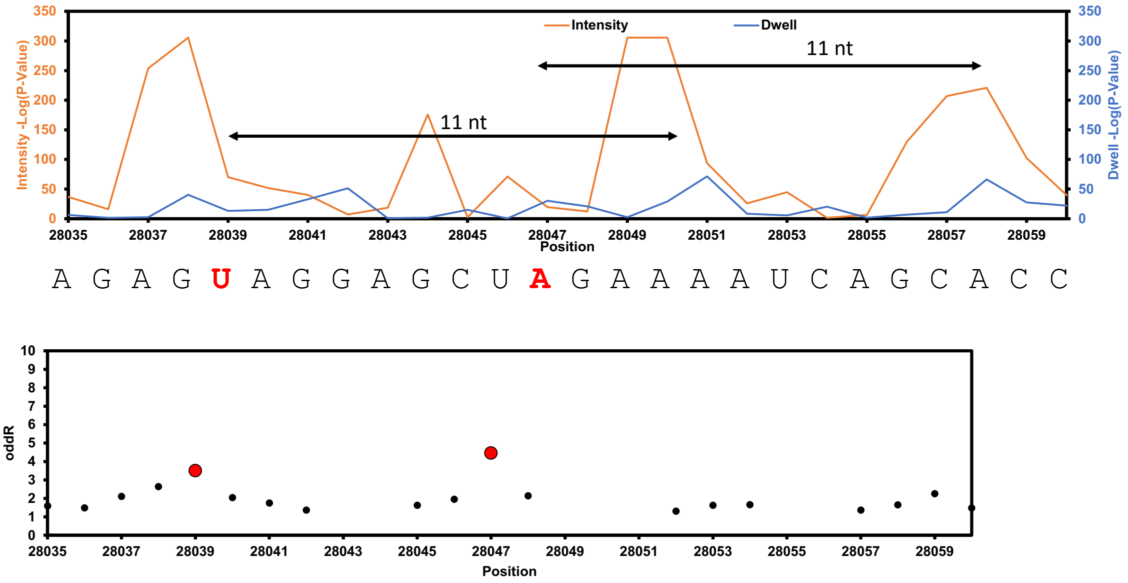


#### U28759 for TRS-8

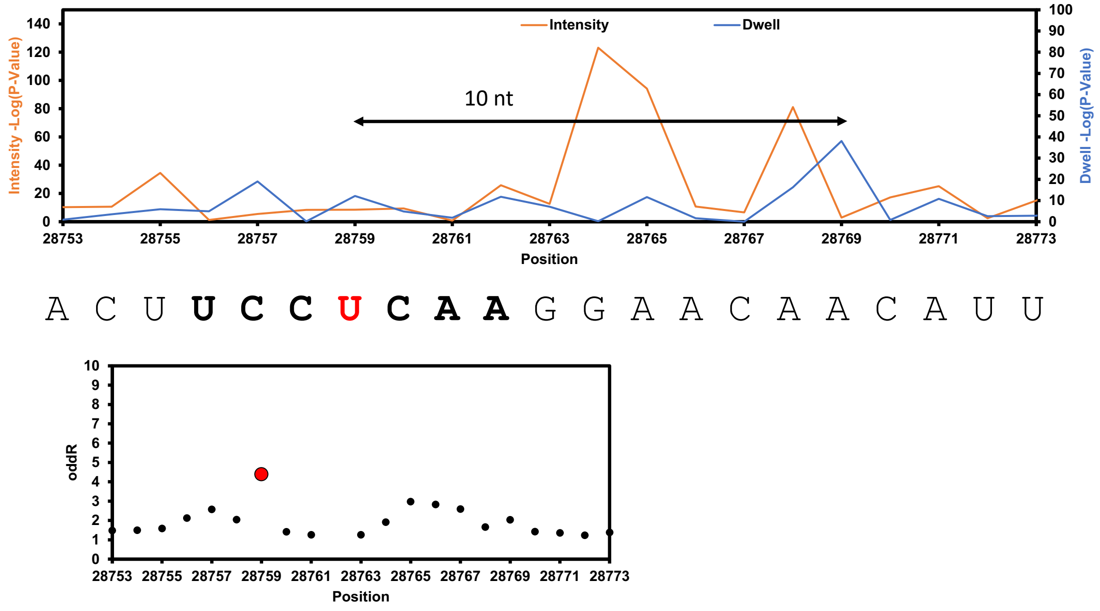

### U28927 for TRS-8

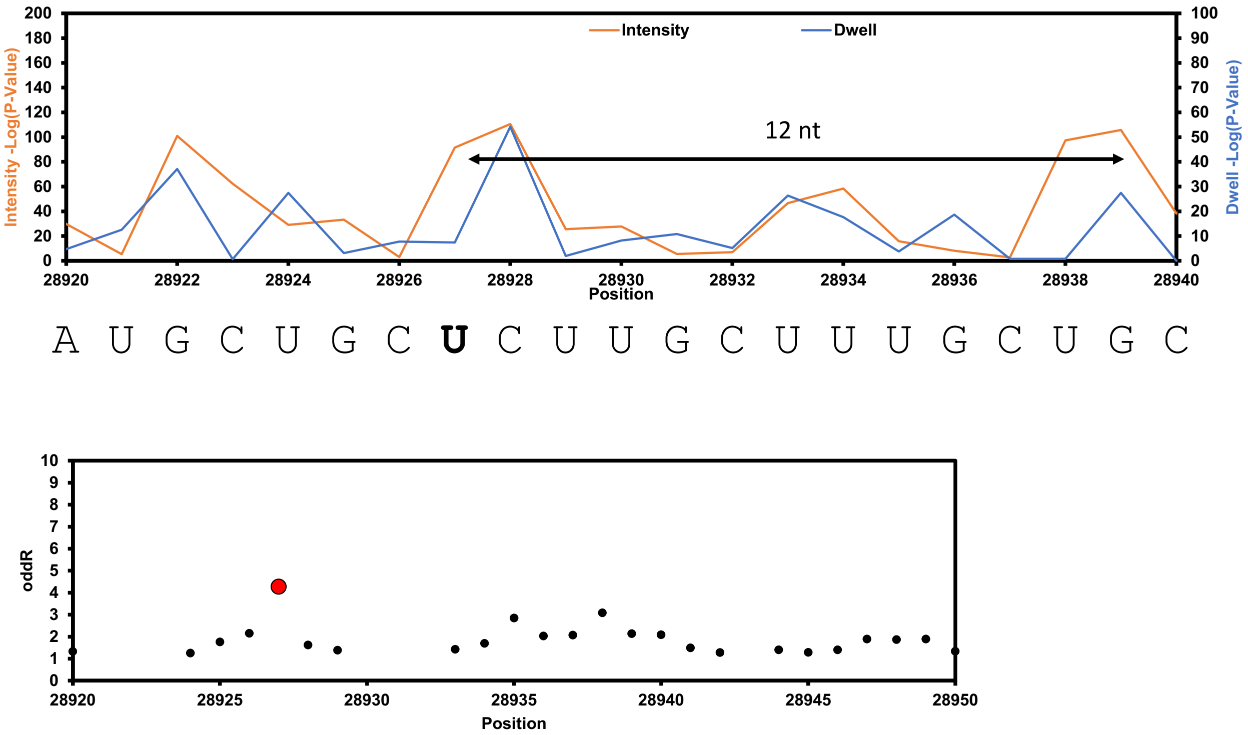


### U29418 in TRS-8

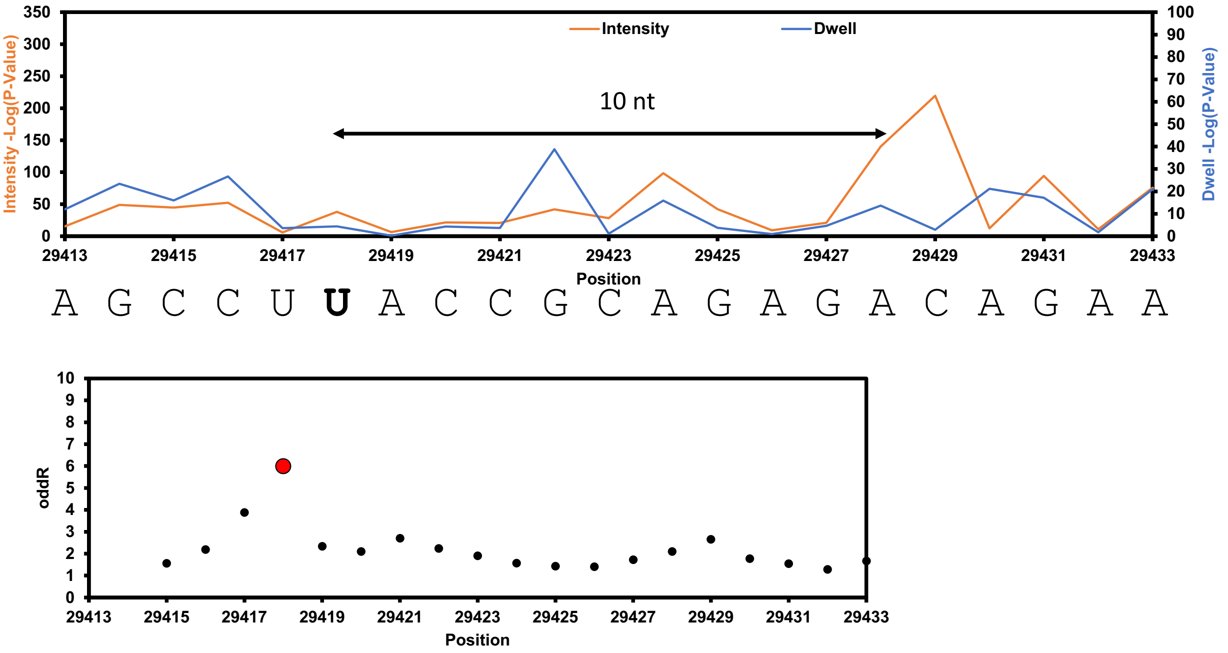

**Figure S19**. Data analysis for TRS-N from SARS-CoV-2.

In each block of data, the top plot is for the Nanocompore analysis and the bottom plot is for the Eligos2 analysis to compare the cellular SARS-CoV-2 RNA to the IVT RNA from publicly available data.[1,2] In the Nanocompore plots, the statistically significant differences in the current intensity are shown in orange and the dwell times are shown in blue. In the Eligos2 plots, the significant nucleotide is shown in the red data point. Each block of data is for a single site found by the Nanocompore analysis, long-range dwell impact (black arrow with distance marked), and Eligos2 analysis.

#### U28759 for TRS-N

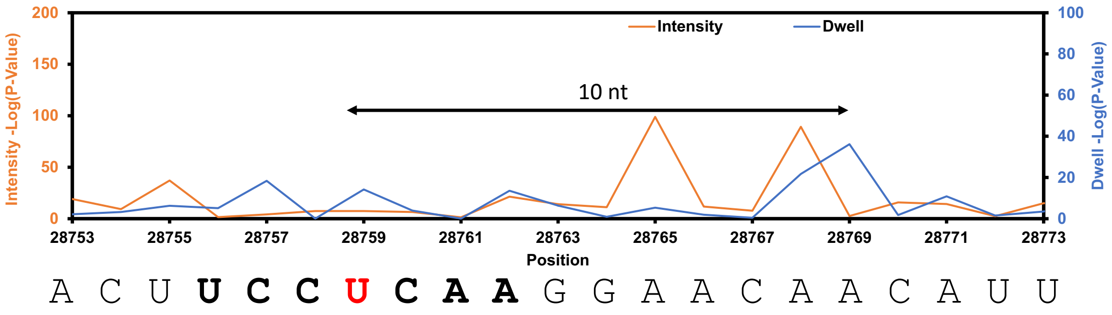


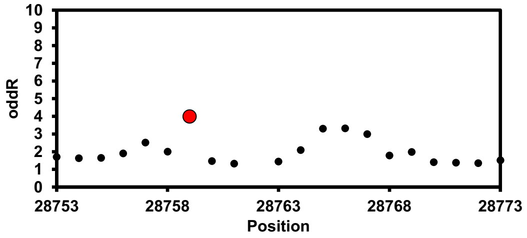


#### U28927 for TRS-N

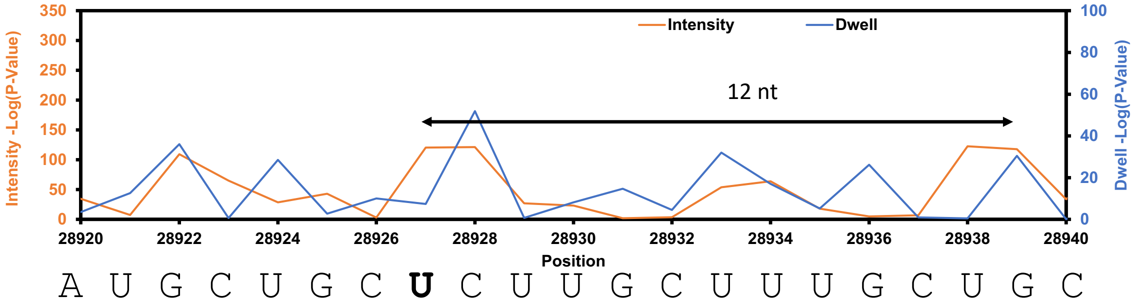


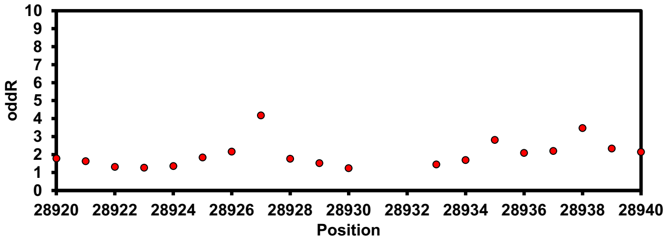

### U29418 for TRS-N

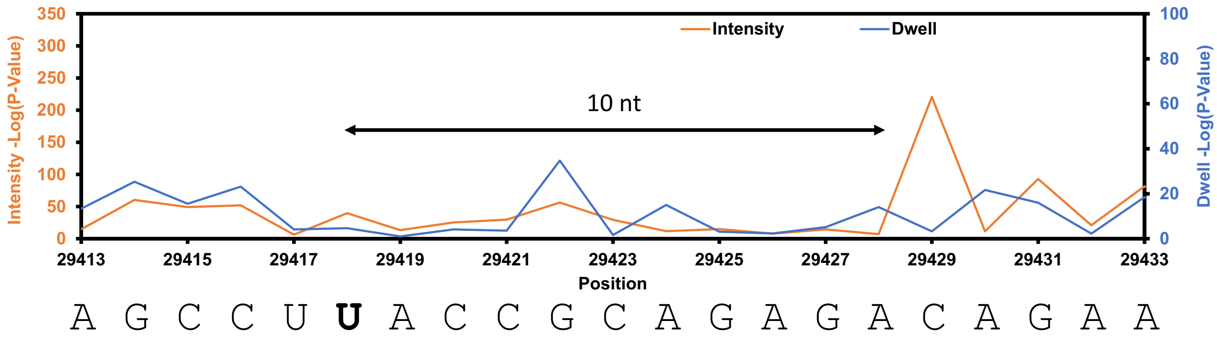


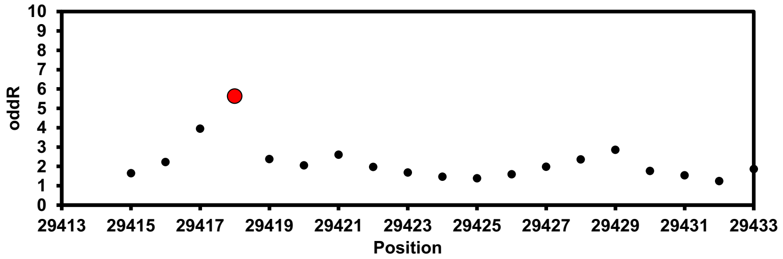

**Figure S20**. Summary of the Ψ epitranscriptomic sites found in SARS-CoV-2.

| Location | Sequence | TRS-S | TRS-3a | TRS-M | TRS-E | TRES-6 | TRS-7a | TRS-7b | TRS-8 | TRS-N |
|---|---|---|---|---|---|---|---|---|---|---|
| | Bold = | | | | | | | | | |
| | Site Modified | | | | | | | | | |
| U22322 | AU**U**CU | X | | | | | | | | |
| U23317 | CU**U**GA | X | | | | | | | | |
| U27164 | AG**U**GA | X | X | X | X | X | | | | |
| U28039 | AG**U**AG | | X | X | X | X | X | X | X | |
| U28417 | AA**U**AC | X | | | | | | | | |
| U28759 | CC**U**CA | X | X | X | X | X | X | X | X | X |
| U28927 | GC**U**CU | X | X | X | X | X | X | X | X | X |
| U29418 | CU**U**AC | X | X | X | X | X | X | X | X | X |

The TRSs from SARS-CoV-2 decrease in length from S, 3a, M, E, 6, 7a, 7b, 8, to N. The conserved modified sites are shown in red, and they were found in all TRSs that the length of the sub-genomic RNA would permit. The exceptions is U28039 that was not found in TRS-S. The table does not include complete analysis for all sites that gave the base calling error and long-range dwell time impact in TRS-6, 7a, 8, and N.

**Figure S21**. Predicted folding for the region flanking the conserved Ψ sites in the TRSs from SARS-CoV-2.

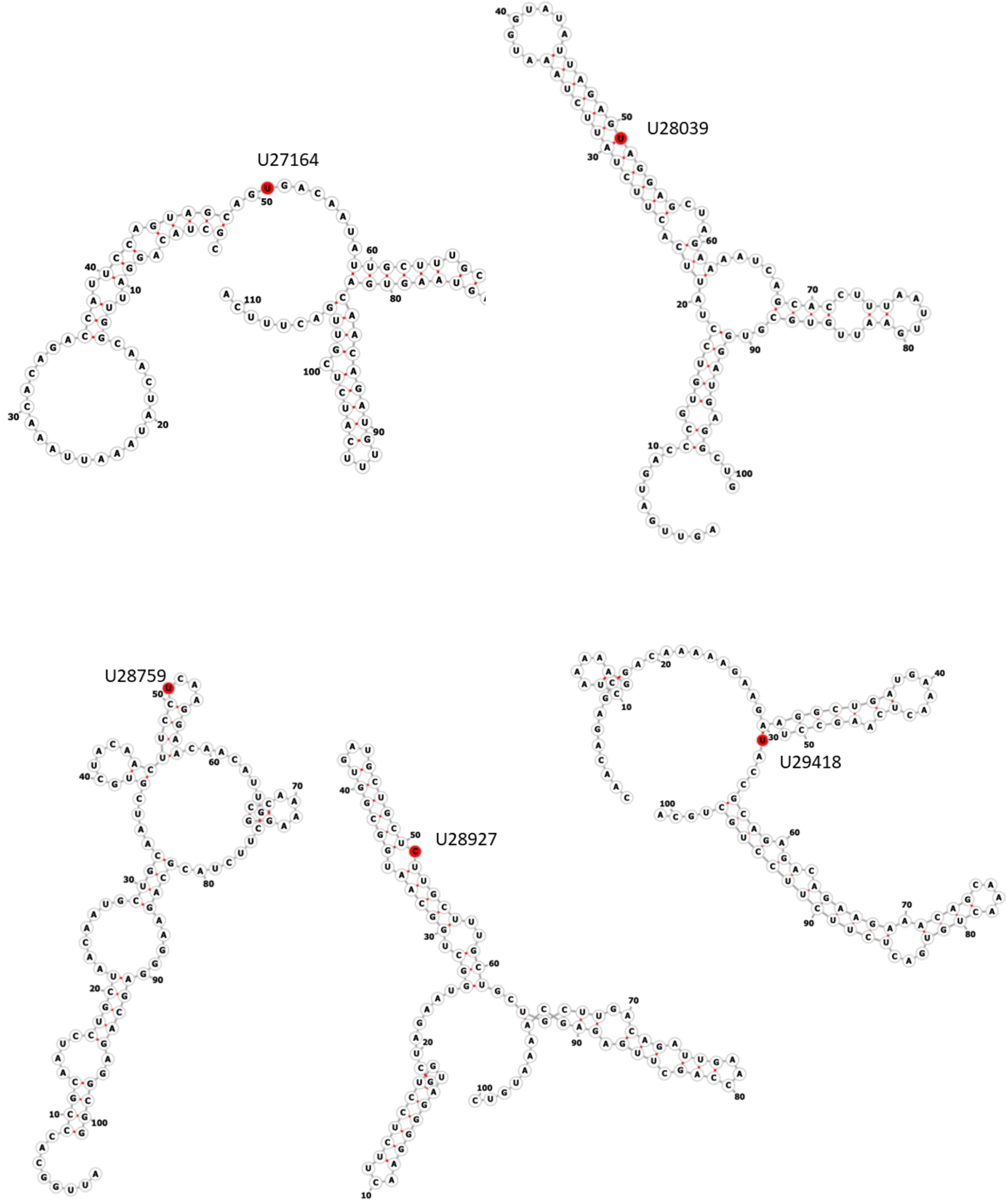
